# Commercial ChIP-Seq library preparation kits performed differently for different classes of protein targets

**DOI:** 10.1101/2022.04.19.488832

**Authors:** MS Simper, Della Coletta L, S Gaddis, K Lin, CD Mikulec, Y Takata, MW Tomida, D Zhang, DG Tang, MR Estecio, J Shen, Y Lu

## Abstract

**Background:** ChIP-Seq is a powerful method commonly used to study global protein-DNA interactions including both transcription factors and histone modifications. We have found that the choice of ChIP-Seq library preparation protocol plays an important role in overall ChIP-Seq data quality. However, very few studies have compared ChIP-Seq libraries prepared by different protocols using multiple targets and a broad range of input DNA levels.

**Results:** In this study, we evaluated the performance of four ChIP-Seq library preparation protocols [NEB NEBNext Ultra II, Roche KAPA HyperPrep, Diagenode MicroPlex, and Bioo (now PerkinElmer) NEXTflex] on three target proteins, chosen to represent the three typical signal enrichment patterns in ChIP-Seq experiments: sharp peaks (H3K4me3), broad domains (H3K27me3) and punctate peaks with a protein binding motif (CTCF). We also tested a broad range of different input DNA levels from 0.10 to 10 ng for H3K4me3 and H3K27me3 experiments.

**Conclusions:** Our results suggest that the NEB protocol may be better for preparing H3K4me3 (and potentially other histone modifications with sharp peak enrichment) libraries; the Bioo protocol may be better for preparing H3K27me3 (and potentially other histone modifications with broad domain enrichment) libraries, and the Diagenode protocol may be better for preparing CTCF (and potentially other transcription factors with well-defined binding motifs) libraries. For ChIP-Seq experiments using novel targets without a known signal enrichment pattern, the NEB protocol might be the best choice as it performed well for each of the three targets we tested across a wide array of input DNA levels.

## INTRODUCTION

Chromatin immunoprecipitation followed by high-throughput sequencing (ChIP-Seq) has become the method of choice for studying global protein-DNA interactions and chemical modifications of histone proteins. Since Keji Zhao and colleagues first coined the term “ChIP-Seq” in 2007 (1), more than 6500 publications have employed ChIP-Seq methods to: 1) survey interactions between protein and DNA to provide insight into mechanisms central to biological processes and disease states; 2) identify the genomic locations of chromatin-associated proteins; and 3) identify post-translational modifications affecting histones, chromatin modifying complexes, and other chromatin-associated proteins. Many factors must be considered and procedures optimized to complete successful ChIP-Seq experiments. These include, but are not limited to, the amount of tissue and/or number of cells available; the degree to which proteins are crosslinked to chromatin by formaldehyde; the method of chromatin fragmentation (e.g. sonication or enzymatic cleavage); the precision of fragment size selection; the quality of the antibody selected for immunoprecipitation; the chosen library preparation protocol; the cluster generation and sequencing platform; and post-sequencing data quality checks and subsequent data processing. To establish proper standards for the ChIP-Seq research community, the ENCODE and modENCODE consortia set guidelines for many of these factors, especially for validating antibodies, replicating experiments, required sequencing depth, and scoring and evaluating ChIP-Seq data (2).

Although many factors can introduce technical bias affecting next-generation sequencing (NGS) outcomes, the main sources of bias stem from chromatin structure, PCR amplification, and read-mapping effects (3). Over 10 years of operation as a next-generation core facility serving 120 laboratories, our experience indicates that the ChIP-Seq library preparation protocol plays an important role in overall ChIP-Seq outcomes, particularly when using ultra-low levels of input DNA. Smaller amounts of template require more PCR cycles to generate enough material for sequencing, but amplification bias increases with each PCR cycle (3). In cases with limited numbers of cells and/or amounts of input DNA, increased numbers of unmapped reads and PCR duplications are common (4). Over the past several years, many commercial kits have become available for ChIP-Seq library preparation, including kits developed for use with limited numbers of cells and/or ultralow DNA input. Sundaram et al. compared seven commercial and/or home-made ChIP-Seq library preparation methods at two input DNA levels (1 ng and 0.1 ng) using a PCR-free dataset as a reference (5). Each kit was evaluated by performing ChIP-Seq for H3K4me3 and then comparing unmapped reads, PCR amplification-derived duplicates, reproducibility, and sensitivity and specificity relative to the PCR-free reference dataset (5). They found that the Swift Biosciences Accel-NGS 2S ChIP-Seq library preparation method performed the best, the Rubicon Genomics ThruPLEX kit performed the second best, and the Sigma-Aldrich SeqPlex method performed poorly.

Earlier studies provided the research community with valuable information; however, they evaluated only a single target (4–6) and included neither commonly used ChIP-Seq library preparation kits (e.g. Next Ultra II kit from New England BioLabs (NEB) and KAPA HyperPrep kit from Roche) nor kits specificially designed for low-input samples (e.g. the Diagenode kit for low input). The Bioo NEXTflex ChIP-Seq kit (Perkin Elmer) was included in our evaluation because a number of our customers used its earlier version and generated good quality data. Here, we compared the performance of these four ChIP-Seq library preparation protocols (NEB, KAPA, Diagenode and Bioo) on three separate targets: histone H3 trimethylated on lysine 4 (H3K4me3), histone H3 trimethylated on lysine 27 (H3K27me3), and the transcription factor (TF) CCCTC-binding factor (CTCF). We carefully chose these targets to represent the three typical outcomes seen in ChIP-Seq experiments: targets that display sharp peaks, as expected for H3K4me3; targets that display broad peaks, as expected for H3K27me3; and targets that display discrete, punctate peaks, as expected for proteins that bind to specific DNA sequences, like CTCF (6). While CTCF is not representative of all TF given its abundant number of sites, it’s an example of narrow peak binding with the ready availability of well-validated antibodies. To test a broad range of input levels, we used six different amounts of input DNA, ranging from 0.1 to 10 ng, for H3K4me3 and H3K27me3 ChIP-Seq. All of our ChIP-Seq libraries were evaluated with respect to sequencing library complexity, reproducibility, and specific quality metrics suitable for the enrichment patterns of the three different protein targets. Our study indicates that the ChIP-Seq library preparation protocols performed differently for different classes of protein targets. The NEB protocol may be the best choice for H3K4me3 (and potentially other histone modifications with sharp peak enrichment); the Bioo protocol may be the best choice for H3K27me3 (and potentially other histone modifications with broad domain enrichment) though not at very low DNA levels; and the Diagenode protocol may be the best choice for CTCF (and potentially other transcription factors that bind to specific DNA sequence motifs). For ChIP-Seq experiments that target proteins with unknown signal enrichment patterns, the NEB protocol might be a good choice as it performed well with all three targets and the NEB libraries were behaved consistently across different input DNA levels.

## MATERIALS AND METHODS

### Cell culture and fixation

The androgen-sensitive human prostate adenocarcinoma cell line LNCaP was purchased from the American Type Culture Collection (ATCC) and cultured in RPMI 1640 (+) L-Glutamine medium from Gibco Life Technologies (Thermo Fisher Scientific, Waltham, MA) supplemented with 10% fetal bovine serum, 100 units/ml penicillin-streptomycin, and 1 mM sodium pyruvate. This line was authenticated regularly in our institutional CCSG Cell Line Characterization Core and also examined to be free of mycoplasma contamination. Cells were maintained in 5% CO2 at 37°C and cultured in 10 cm plates to a confluency of 70-80% before fixation. LNCaP cells were fixed in 1% methanol-free formaldehyde (Thermo Fisher Scientific) in RPMI for 10 min at room temperature followed by quenching for 5 min in 125 mM glycine (Sigma, St. Louis, MO) with low-speed shaking on an orbital platform. Plates were washed 2× with ice-cold phosphate-buffered saline (PBS) (pH 7.4) (Thermo Fisher Scientific) to eliminate any residual media, and 7 ml cold PBS containing 1× Complete Protease Inhibitors Cocktail, EDTA free (Roche, Basel, Switzerland) were immediately added. Plates were kept on ice while the cells were harvested by scraping. Cells were transferred to 15 ml conical tubes (2 plates/tube) and collected by centrifugation (4 min at 805 × g at 4°C). Cells were immediately used for chromatin preparation.

### Chromatin preparation

Fixed LNCaP cell pellets were resuspended by pipetting in sodium dodecyl sulfate (SDS) lysis buffer [1% SDS, 10mM EDTA, 50mM Tris, pH 8.1, and 1X Complete Protease Inhibitors Cocktail (Roche)]and incubated for at least 10 min on ice. Three hundred μl of SDS lysis buffer were added for every 2-3 million cells, and 300 μl of cell lysates were transferred to a 1.5 μl tube from Diagenode, Inc. (Denville, NJ) for sonication. Sonication was performed using a Diagenode Bioruptor Plus with 22 cycles of 30 sec high power (on) followed by 30 sec rest (off) at 4°C. After the intial 22 sonication cycles, the samples were allowed to rest on ice for 15 min, then subjected to an additional 22 cycles of sonication using the same conditions. This shearing protocol routinely resulted in a chromatin preparation with fragments sized between 200 bp and 700 bp. The cell lysates were cleared by centrifugation (10 min at 14,500 × g at 4°C), and the supernatant was transferred to a new tube. Two µl of the supernatant were diluted with 180 μl SDS lysis buffer and used for reverse-crosslinking. The DNA was purified using the QIAquick PCR Purification kit protocol (Qiagen, Hilden, Germany), quantified using a Qubit fluorometer, and 1 ng of purified DNA was loaded into an Agilent 2100 Bioanalyzer (Santa Clara, CA) using High Sensitivity (HS) DNA reagents to ensure that the desired size profile was obtained.

### Chromatin Immunoprecipitation

The following antibodies were used for chromatin immunoprecipitation: anti-histone H3 [ab1791] from Abcam (Cambridge, United Kingdom), and anti-H3K4me3 [17-614], anti-H3K27me3 [17-622], and anti-CTCF [07-729] from MilliporeSigma (St. Louis, MO). For each immunoprecipitation, 15 μl of Dynabeads Protein A magnetic beads (Invitrogen, Carlsbad, CA) were combined with 15 μl Dynabeads Protein G magnetic beads (Invitrogen) and washed 3 × in blocking solution (1× PBS/0.5% bovine serum albumin). After the final wash, the Dynabeads were resuspended in 250 μl blocking solution and 10 μg antibody were added for each 15-30 μg chromatin. The mixture was rotated overnight at 4°C and washed 3× with blocking solution before resuspension in 100 μl blocking solution. The chromatin lysate was diluted 1:10 using ChIP dilution buffer [16.7 mM Tris-HCl (pH 8.1), 167 mM NaCl, 1.2 mM EDTA, 0.01% SDS, 1.1% Triton X-100] with protease inhibitors, and 100 μl of diluted lysate were removed and saved as Input for comparison during analysis. One ml diluted lysate was then added to the antibody/bead mixture and incubated overnight on a rotator at 4°C. A magnet was used to collect the beads which were then washed 3× with 1 ml RIPA washing buffer (Teknova, Hollister, CA). Each supernatant was removed and transferred to a fresh tube. A final wash was performed with 1 ml 1× TE (Promega, Madison, WI) containing 50 mM NaCl, and the beads were collected by centrifugation (1 min at 960 × g at 4°C) before resuspension in 110 μl SDS lysis buffer. Crosslinking was reversed and DNA was purified from both the immunoprecipitated and the input samples before analysis using a Bioanalyzer. To validate the ChIP experiments prior to library preparation and sequencing, vendor-provided primers were used for qPCR as follows: hGAPDH-promoter region specific primers for H3K4me3, hAlpha satellite control primers for H3K27me3, and hSCN4A primers for CTCF.

### NEBNext^®^ Ultra II™ DNA Library Prep Kit for Illumina^®^

ChIP-Seq libraries were prepared using the NEBNext^®^ Ultra II™ DNA Library Prep Kit for Illumina^®^ (New England BioLabs, Inc., Ipswich, MA) following the manufacturer’s protocol. Briefly, ng (1, 5, and 10 ng) and sub-ng (100, 250, and 500 pg) amounts of fragmented DNA were used to repair the ends before ligation to a NEBNext Adapter for Illumina sequencing. The adapter-ligated DNA was then enriched using PCR [1 cycle at 98°C for 30 sec; followed by a specific number of cycles depending on the amount of input DNA (Table 1), at 98°C for 10 sec, 65°C for 75 sec; and 1 cycle at 65°C for 5 min]. DNA was purified using AMPure XP beads (Beckman Coulter, Brea, CA) after adapter ligation and PCR enrichment.

**Table 1:**
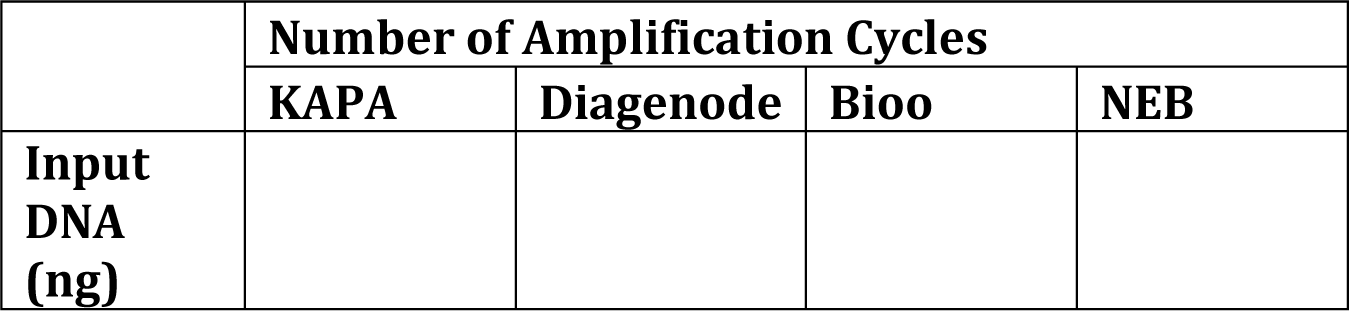
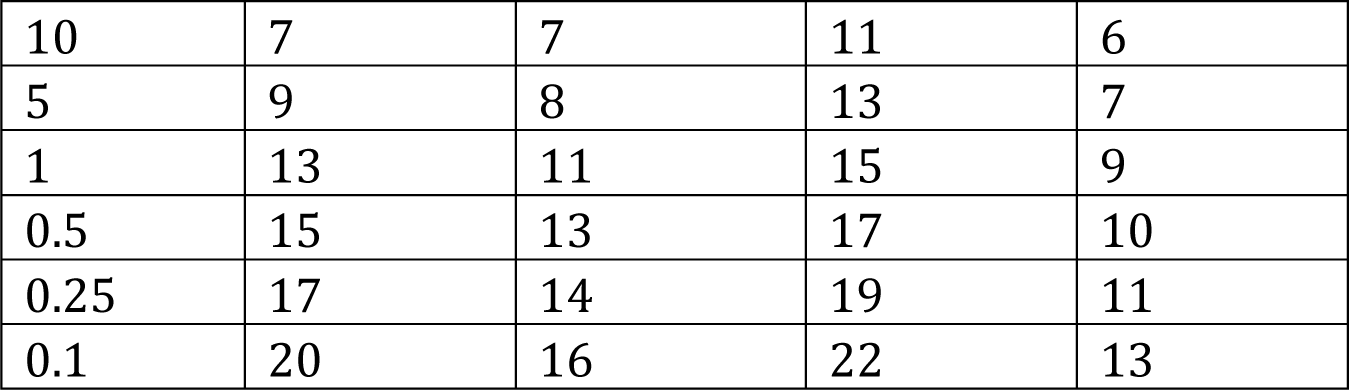
PCR amplification cycle number specific to kit and input levels.

|  | Number of Amplification Cycles |  |  |  |
| --- | --- | --- | --- | --- |
|  | KAPA | Diagenode | Bioo | NEB |
| Input DNA (ng) |  |  |  |  |
| 10 | 7 | 7 | 11 | 6 |
| 5 | 9 | 8 | 13 | 7 |
| 1 | 13 | 11 | 15 | 9 |
| 0.5 | 15 | 13 | 17 | 10 |
| 0.25 | 17 | 14 | 19 | 11 |
| 0.1 | 20 | 16 | 22 | 13 |

### KAPA HyperPrep Kit

ChIP libraries were prepared using the KAPA HyperPrep Kit (Roche) following the manufacturer’s protocol. Briefly, ng (1, 5, and 10 ng) and sub-ng (100, 250, and 500 pg) amounts of fragmented DNA were used to repair the ends before ligation to a diluted NEXTflex DNA adapter (PerkinElmer, Waltham, MA). The adapter-ligated DNA was then enriched using PCR [1 cycle at 98°C for 45 sec; followed by a specific number of cycles depending on amount of input DNA (Table 1), at 98°C for 15 sec, 60°C for 30 sec, 72°C for 30 sec; and 1 cycle at 72°C for 1 min]. DNA was purified using AMPure XP beads (Beckman Coulter) after adapter ligation and PCR enrichment.

### Diagenode MicroPlex Library Preparation Kit v2

ChIP libraries were prepared using the Diagenode MicroPlex Library Preparation Kit v2 (Diagenode) following the manufacturer’s protocol. Briefly, ng (1, 5, and 10 ng) and sub-ng (100, 250, and 500 pg) amounts of DNA were repaired to create blunt ends for ligation to a MicroPlex stem-loop adapters. The adapter-ligated DNA was then enriched and indexed using PCR (1 cycle at 72°C for 3 min; 1 cycle at 85°C for 2 min; 1 cycle at 98°C for 2 min; 4 cycles at 98°C for 20 sec, 67°C for 20 sec, 72°C for 40 sec; followed by a specific number of cycles depending on the amount of DNA input [see Table 1], at 98°C for 20 sec, 72°C for 50 sec). The total number of PCR cycles listed in Table 1 does not include the four cycles of stage 1, and is just the number of stage 2 cycles. DNA was purified using AMPure XP beads (Beckman Coulter) after adapter ligation and PCR enrichment.

### NEXTflex ChIP-Seq Library Prep Kit for Illumina Sequencing (Bioo)

ChIP libraries were prepared using the NEXTflex ChIP-Seq Kit (Bioo Scientific, now PerkinElmer) following the manufacturer’s protocol. Briefly, ng (1, 5, and 10 ng) and sub-ng (100, 250, and 500 pg) amounts of fragmented DNA were end-repaired and adenylated prior to ligation to a NEXTflex ChIP adapter. The adapter-ligated DNA was then enriched using PCR [1 cycle at 98°C for 2 min; followed by a specific number of cycles depending on amount of DNA input (Table 1) at 98°C for 30 sec, 65°C for 30 sec, 72°C for 1 min; and 1 cycle at 72°C for 4 min]. DNA was purified using AMPure XP beads (Beckman Coulter) after repair, adapter ligation, and PCR enrichment.

### Sequencing

Each ChIP-Seq library was checked for quality using a 2200 TapeStation (Agilent Technologies). A KAPA Library Quantification Kit (Roche) was used to quantify the libraries for pooling, and a final concentration of 1.5 nM was loaded onto an Illumina cBot for cluster generation before sequencing with an Illumina HiSeq 3000 using a single read 50 bp run.

### Bioinformatics analysis

#### Mapping

In order to make meaningful comparisons with our own existing data (data not shown) all ChIP-Seq reads were trimmed to 36 bp and mapped to the human genome (hg19). Mapping was performed using Bowtie (version 1.1.2) (7), allowing for no more than two mismatches and retaining only reads that were mapped to unique positions. 91-96% of reads were mapped to the human genome, with 78-82% being uniquely mapped. To avoid PCR bias, when multiple reads were mapped to the same genomic position, only one read was retained for analysis.

#### Peak/domain calling

H3K4me3 and CTCF peaks were detected using MACS (version 1.4.2, p-value cutoff 1e-5 and window size 300 bp) (8). Peaks that overlapped ENCODE blacklisted regions (6) were removed. To avoid any possible effects of total H3 library quality, H3K4me3 peak calling was performed without a control library reference. CTCF peaks in each library were called by taking the corresponding total input library as a control. The H3K27me3 enriched domains were initially identified using the enriched domain detector (EDD version 1.1.16) (9), without any significance threshold (i.e. the FDR cutoff for EDD was set to be larger than 1). The EDD gap penalty and bin size were set to 10 and 20 kb, respectively. After peak calling with EDD, for each 20 kb bin, the z-score was calculated as: 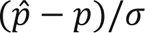, where 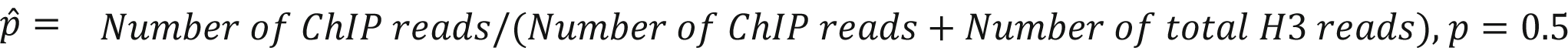 and 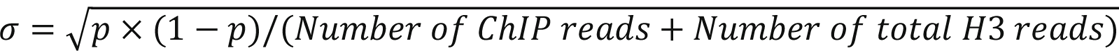. The p-value for each candidate domain was calculated using Stouffer’s Z-score method by combining all the bins within the putative domain. The p-values were corrected using the method of Benjamini & Hochberg (BH). The domains with FDR ≤ 0.05 were called as significantly enriched with H3K27me3. To minimize the effect of total H3 library quality, a combined total H3 was used as a control for the detection of H3K27me3 enriched domains with 15M reads from the total H3 library prepared by each protocol at the 10 ng level.

### Signal landscape

For the H3K4me3 and CTCF libraries, each read was extended by 150 bp (i.e. the expected average fragment size) to its 3’ end. The number of reads mapping to each genomic position was normalized to a total of 10 M mapped reads, averaged over every 10 bp window, and displayed in the UCSC Genome Browser (http://genome.ucsc.edu/) (10). For each H3K27me3 library, the log2 ratio of the number of reads in each 20 kb window over the combined total H3 following normalization, based on the total number of mapped reads, was calculated and displayed in UCSC Genome Browser.

### Gene annotation

Genes from GENCODE Release 29 (11) were used to annotate H3K4me3 peaks and H3K27me3 enriched domains. The promoter region was defined as -1000 bp to +500 bp of a transcription start site (TSS).

### RNA-Seq analysis

For gene expression data, two RNA-Seq libraries were generated from LNCaP cells and sequenced (2 x 75 bp paired-end protocol) using an Illumina HiSeq 2500 instrument. Although it was the same cell line, the RNA-Seq was performed at a different time from a different set of cells. Each pair of reads represents a cDNA fragment from the library. The reads were mapped to the human genome (hg19) using TopHat (version 2.0.10) (12). The number of fragments corresponding to each known gene in GENCODE (Release 29) was enumerated using htseq-count (HTSeq package version 0.6.0) (13). Genes shorter than 200 bp and those coding for rRNAs were removed prior to analysis. The FPKM (number of Fragments Per Kilobase per Million fragments) value for each gene was calculated and averaged over the two replicates.

### Quality of peak calling for H3K4me3

The quality of peak calling for H3K4me3 was measured as the percentage of peaks in promoter regions vs. the percentage of expressed genes marked by peaks. The promoter region was defined as -1000 bp to +500 bp of a TSS. To identify the expressed genes, we plotted log10(FPKM) values for all of the genes in a histogram, which revealed a bimodal shape. The histogram was fitted to a density curve using the density function in R. Genes with FPKM values larger than the point of minimal density between the two density peaks were defined as expressed genes.

### H3K4me3 aggregated signal around TSS/enhancer

The 3,000 bases 5’ and 3’ of a TSS as well as the 3000 bases 5’ and 3’ from the center of an enhancer were subdivided into 100 bp bins. For each H3K4me3 library, the RPKM (reads per kilobase per million mapped reads) value was calculated for each bin and averaged over all TSSs or enhancers. Enhancers were defined as DNaseI hypersensitive sites that are separated by at least 10 kb from a TSS. DNaseI hypersensitivity data was obtained from the UCSC genome browser (http://genome.ucsc.edu/cgi-bin/hgTrackUi?db=hg19&g=wgEncodeUwDnase).

### Correlation of H3K27me3 signal with gene expression

For each library, the log2 ratio was calculated from the number of H3K27me3 reads within -2000 bp to +1000 bp of each gene’s TSS over the number of total H3 reads. Log2 ratio values vs. gene expression values (based on FPKM) were compared using Spearman’s correlation. The TSS of the longest transcript was used to represent each gene’s TSS.

### Correlation of H3K27me3 signal with chromatin accessibility

For each library, the log2 ratio was calculated from the number of H3K27me3 reads in each 20 kb window over the number of total H3 reads. Log2 ratio values vs. chromatin accessibility scores were compared in 20 kb windows using Spearman’s correlation. Chromatin accessibility scores were calculated as RPKM values based on DNaseI-Seq data for LNCaP cells downloaded from the UCSC genome browser (http://genome.ucsc.edu/cgi-bin/hgTrackUi?db=hg19&g=wgEncodeUwDnase). As H3K27me3 is typically enriched in facultative heterochromatin (14), windows lacking a TSS for a gene were excluded from the analysis.

### Identification of CTCF motifs in CTCF peaks

The identification of canonical CTCF motifs near CTCF peak summits used the FIMO (15) component of the MEME suite (version 4.10.2) (16) with default parameter settings. The canonical CTCF motif was defined as in JASPAR (17) (Matrix ID MA0139.1).

### Reproducibility curves

For comparing H3K4me3 peaks for each pair of H3K4me3 libraries, the peaks were merged and ranked from the strongest to the weakest based on RPKM values.The percentage of peaks common to both libraries from the same number of top peaks was calculated. A similar approach was applied for comparing H3K27me3 enriched 20 kb windows, except that the windows were ranked by z-scores.

### Number of reads used

For all comparisons, the same number of reads was used for all libraries shown on a single plot; however, the number of reads per plot varies across the plots. The number of reads used in each plot is summarized in Table S2.

## Results

### Experimental design and data quality metrics

The experimental design used for testing the four ChIP-Seq library preparation protocols (Bioo, Diagenode, KAPA, and NEB) is outlined in Figure 1. ChIP DNAs from LNCaP cells were immunoprecipitated with H3K4me3, H3K27me3, and CTCF antibodies before library preparation. These antibodies have been widely used for ChIP (18–23). For the H3K4me3 and H3K27me3 ChIP-Seq analyses, we tested six different amounts of input DNA for each protocol: 10, 5, and 1 ng and 500, 250, and 100 pg. CTCF DNA was undetectable by fluorometry Qubit following immunoprecipitation, therefore; multiple CTCF immunoprecipitations were combined and then divided equally across the four protocols to normalize the amount of input DNA used for each. To ensure reproducibility, two independent sets of ChIP-Seq experiments were performed to assess the H3K4me3 (H3K4me3 sets 1 and 2) and H3K27me3 (H3K27me3 sets 1 and 2) libraries, and three independent sets of ChIP-Seq experiments were performed to assess the CTCF libraries (CTCF sets 1-3). All libraries were sequenced on an Illumina HiSeq 3000 using a single read 50 base pair run (see Table S1 for number of reads generated and mapping rate for each library and Figure S1 for screenshots of signal landscape and peaks called for each library). To assess our ChIP-Seq libraries, we evaluated library complexity for all libraries and reproducibility for the H3K4me3 and H3K27me3 libraries. Based on the known enrichment patterns for our three protein targets, we employed different quality metrics for each: H3K4me3 libraries were evaluated by the fraction of reads in peaks (FRiP), the fraction of reads in promoter regions, and the precision and sensitivity of peak calling; H3K27me3 libraries were evaluated by their correlation with gene expression and chromatin accessibility; and CTCF libraries were evaluated by the number of peaks, the percentage of peaks that contained the CTCF motif, and the distance between the CTCF motif and the peak summit. To avoid batch effect, we made the decision of evaluating reproducibility by comparing neighboring input amounts rather than between the two sets (sets 1 and 2 for both K3K4me3 and H3K27me3) because the two sets were performed more than one year apart.

**Figure 1.**
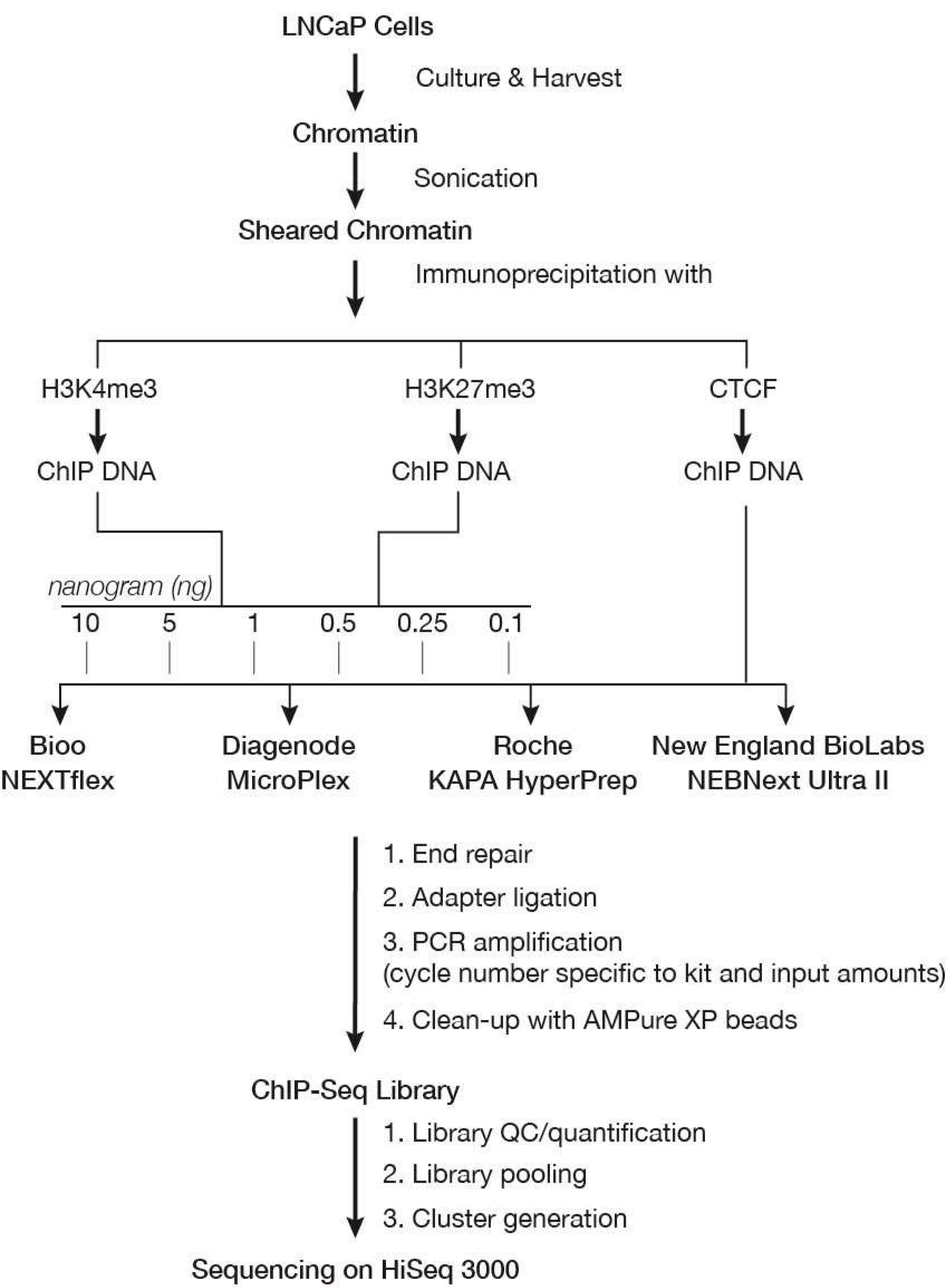
Experimental design. Diagram outlining the experimental design for comparing the Bioo NEXTflex, Diagenode MicroPlex, Roche KAPA HyperPrep, and New England BioLabs NEBNext Ultra II ChIP-Seq library preparation protocols. Two independent ChIP-Seq experiments (sets 1 and 2) were carried out to assess H3K4me3 and H3K27me3 libraries, and three independent ChIP-Seq experiments (sets 1-3) were performed to evaluate CTCF libraries. For H3K4me3 and H3K27me3, six different amounts of input DNA (10, 5, and 1 ng and 500, 250, and 100 pg), were used for library construction; however, due to low recovery of CTCF ChIP DNA, only a single DNA input was used for library construction. The number of amplification cycles for each protocol (Table 1) followed the manufacturers’ recommendations based on the amount of input DNA.

### Library complexity

Library complexity is commonly used to evaluate the quality of ChIP-Seq libraries (2, 3, 24). Low complexity libraries often result from insufficient amounts of starting DNA or over-amplification by PCR and generate less useful data than do high complexity libraries. To evaluate library complexity, we measured the NRF (Non-Redundant Fraction), defined as the ratio of the number of positions in the genome to which uniquely mappable reads and the total number of uniquely mapped reads (2) (Figure 2 and S2). The Bioo libraries yielded the lowest overall complexity, especially at lower amounts of input DNA for both H3K4me3 (Figure 2A and S2A) and H3K27me3 (Figure 2B and S2B). As expected, library complexity decreased with lower amounts of input DNA. The Bioo libraries were the most variable in terms of NRFs, whereas the NEB libraries were the most consistent across all input amounts. Notably, at the 100 pg level, the KAPA libraries showed a much more dramatic drop in complexity than did the Diagenode and NEB libraries, with the exception of H3K4me3 set 2. The results were similar when assessing the CTCF libraries (Figure 2C). Again, Bioo typically generated the lowest complexity library, NEB generated the highest complexity library, and KAPA performance was variable at lower input amounts.

**Figure 2.**
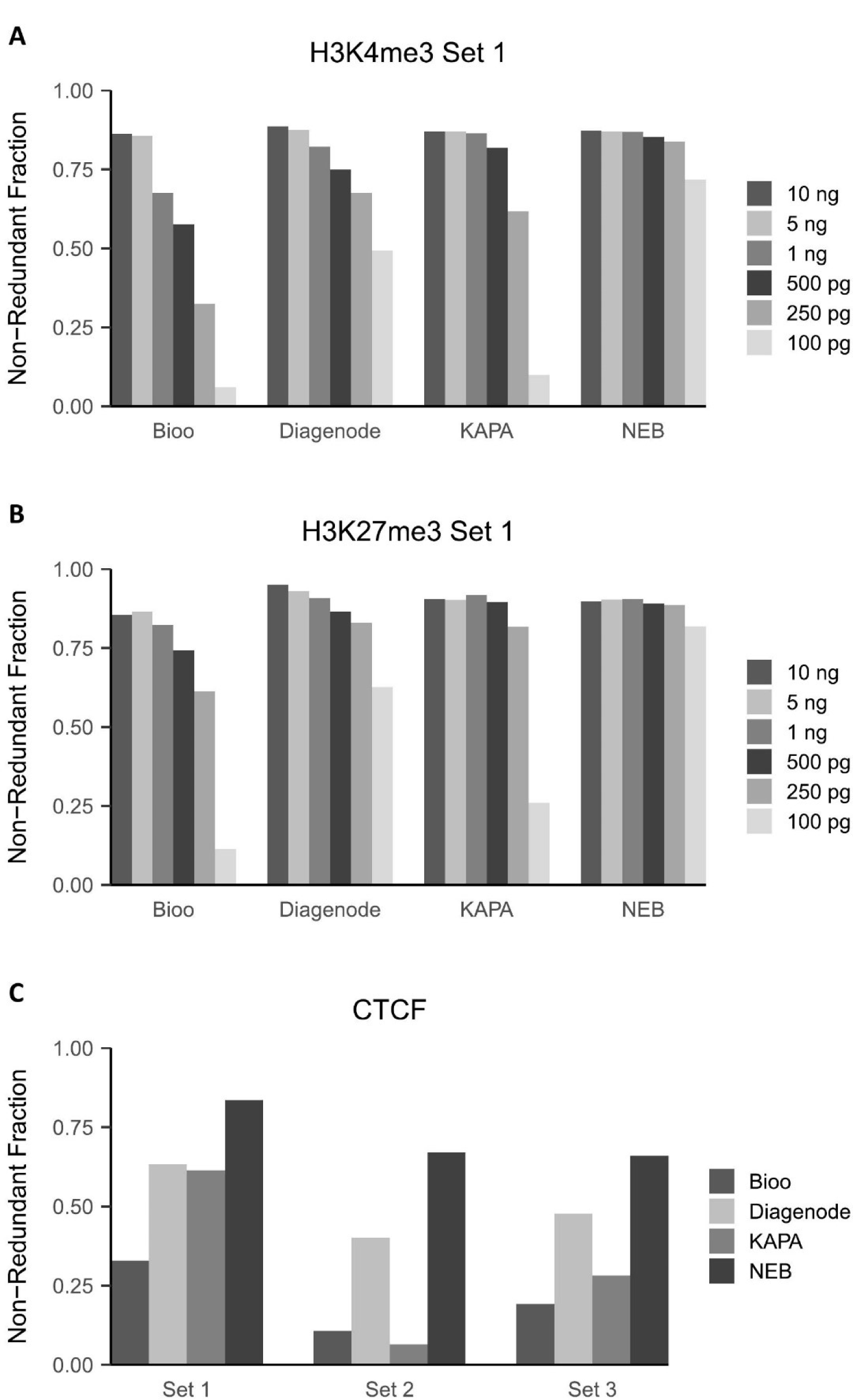
Library complexity measured as Non-Redundant Fraction (NRF). **A.** NRF for H3K4me3 set 1 libraries. **B.** NRF for H3K27me3 set 1 libraries. **C.** NRF for libraries from CTCF sets 1-3.

### H3K4me3 ChIP-Seq

#### Signal portion

To evaluate the portion of true H3K4me3 signal in our libraries, we plotted the fraction of reads in peaks (FRiP) vs. sequencing depth at different DNA inputs (Figure 3 and S3). Given that H3K4me3 is enriched near transcription start sites (TSSs) (1), we also plotted the fraction of reads in promoter regions at each DNA input (Figure S4). Of the tested protocols, the NEB protocol worked well at all DNA input levels, and outperformed the other protocols at the 10 and 5 ng levels; and the Diagenode protocol resulted in the best signal at the picogram DNA levels. The Bioo libraries generated the lowest signal at almost all input levels except the 100 pg level in H3K4me3 set 1, for which, the KAPA libraries generated the lowest signal (Figure 3A, S3A and S4A-B). Surprisingly, although libraries prepared using higher levels of input DNA were expected to have better quality, a general trend within each protocol indicated that the portion of H3K4me3 signal in the library increased when the amount of input DNA decreased (Figure 3B, S3B and S4C-D). The finding that H3K4me3 signal increased as DNA input decreased was also seen both in the aggregated signal curves around TSSs (Figure S5) and by visual inspection of the signal landscape (Figure S1A). This inverse relationship is consistent with a prior study showing H3K4me3 signal over TSSs decreased with increasing amounts of input DNA (5). Regardless, the H3K4me3 signal from the NEB libraries was the most consistent across all input levels.

**Figure 3.**
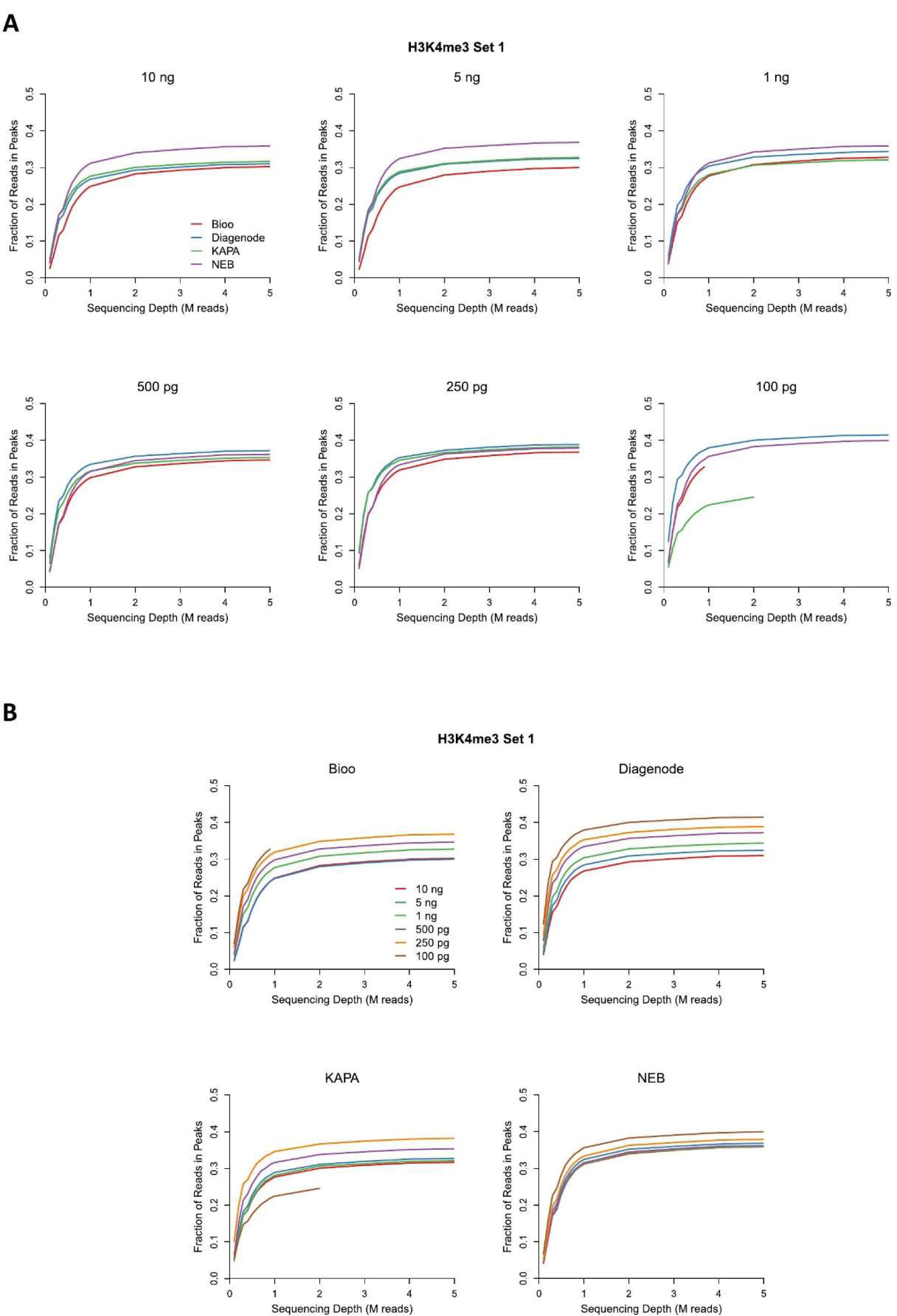
FRiP (fraction of reads in peaks) for H3K4me3 plotted against sequencing depth for H3K4me3 set 1 libraries. **A.** Graphs displayed to highlight differences in protocol performance at specific DNA inputs. **B.** Graphs displayed to facilitate comparisons of a single protocol at different DNA inputs.

#### Peak calling

H3K4me3 is a common histone modification that generally correlates with the promoters of transcriptionally active genes (1). To evaluate the quality of peak calling for H3K4me3 from our libraries, we used the percentage of H3K4me3 peaks in promoter regions as a measurement of precision, and we used the percentage of expressed genes marked by those peaks as a measurement of sensitivity (Figure 4 and S6). Overall, Bioo libraries produced the lowest percentages for both precision and sensitivity across all DNA inputs (Figure 4A and S6A). The differences between the Bioo libraries compared to the others became more obvious with each decrease in input DNA. Peak calling was similar across the other three protocols at all DNA inputs except that the KAPA libraries were inferior at the 100 pg level in H3K4me3 set 1. The quality of peak calling, at different amounts of input DNA within single protocols (Figure 4B and S6B), was clearly decreased at the 250 pg and 100 pg levels in H3K4me3 sets 1 and 2 for the Bioo libraries and at 100 pg in H3K4me3 set 1 for the KAPA libraries but remained stable across all inputs for the Diagenode and NEB libraries.

**Figure 4.**
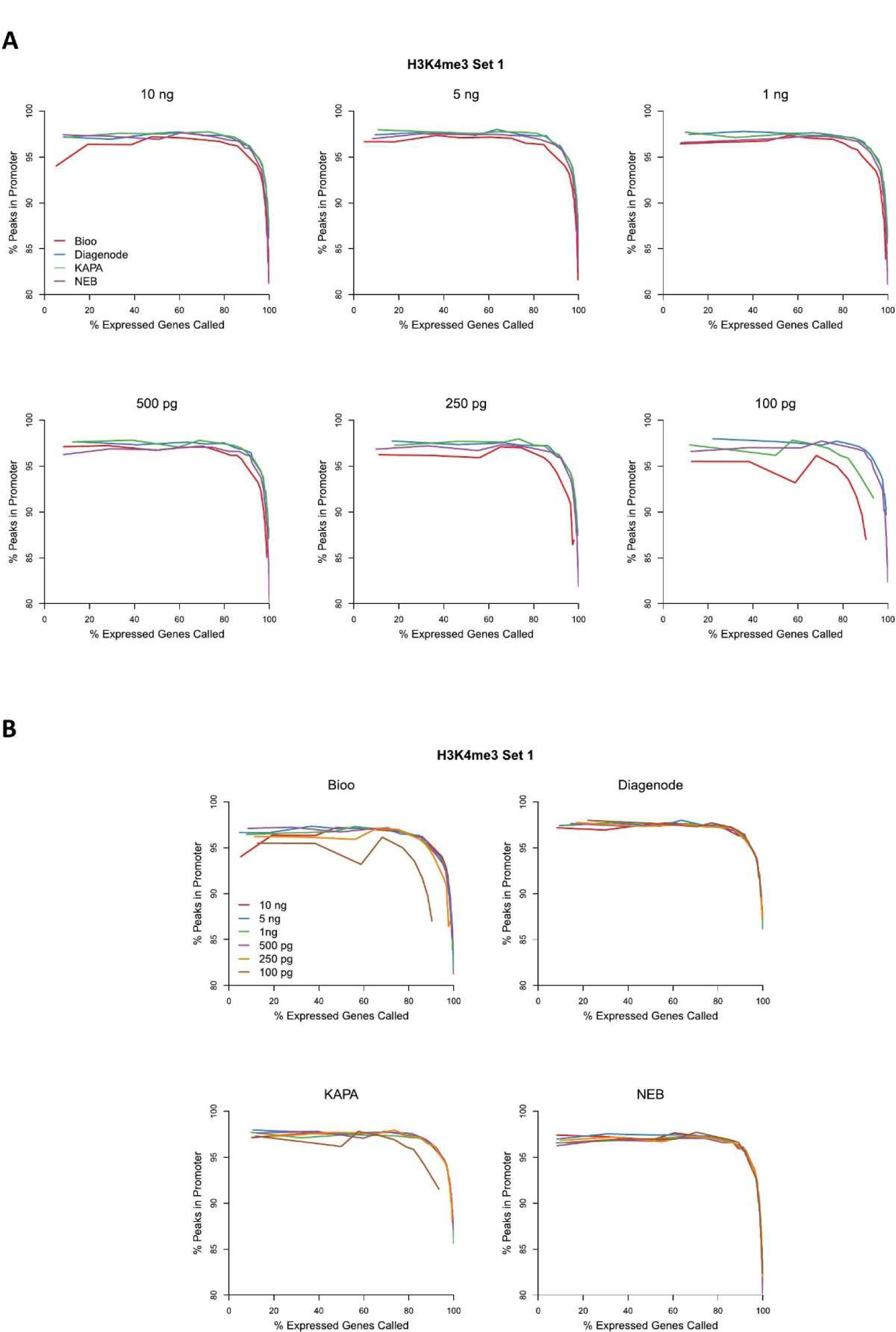
The peak calling quality for H3K4me3 measured as the percentage of peaks in promoter regions vs. the percentage of expressed genes marked by peaks at varying sequencing depth for H3K4me3 set 1 libraries. **A.** Graphs displayed to highlight diffences in the performance of each protocol at specific DNA inputs. **B.** Graphs displayed to facilitate comparisons of a single protocol at different DNA inputs.

### H3K27me3 ChIP-Seq

#### Correlation with gene expression

H3K27me3 is a histone mark typically associated with gene repression (1, 14, 25). We evaluated H3K27me3 ChIP by comparing this mark across the genome in our ChIP-Seq data set to our own RNA-Seq gene expression data for this cell line (LNCaP) (Figure 5 and S7). As expected, H3K27me3 correlated negatively with gene expression in each of our libraries. However, while the KAPA libraries in H3K27me3 set 2 gave rise to the strongest negative correlation at five of the six input DNA levels (250 pg-10 ng), KAPA set 1 libraries had the weakest negative correlation with gene expression regardless of DNA input. This exhibits the inconsistency in the KAPA kit performance between libraries prepared at different times. Among the remaining three protocols, Bioo libraries had the best negative correlation between the presence of H3K27me3 and gene expression except for the 100 pg library in H3K27me3 set 1. NEB was better than or similar to Diagenode at all input DNA levels (Figure 5A and S7A). As expected, H3K27me3 ChIP-Seq libraries prepared from higher amounts of input DNA generally had better negative correlation with gene expression than did ChIP-Seq using lower amounts of DNA (Figure 5B and S7B).

**Figure 5.**
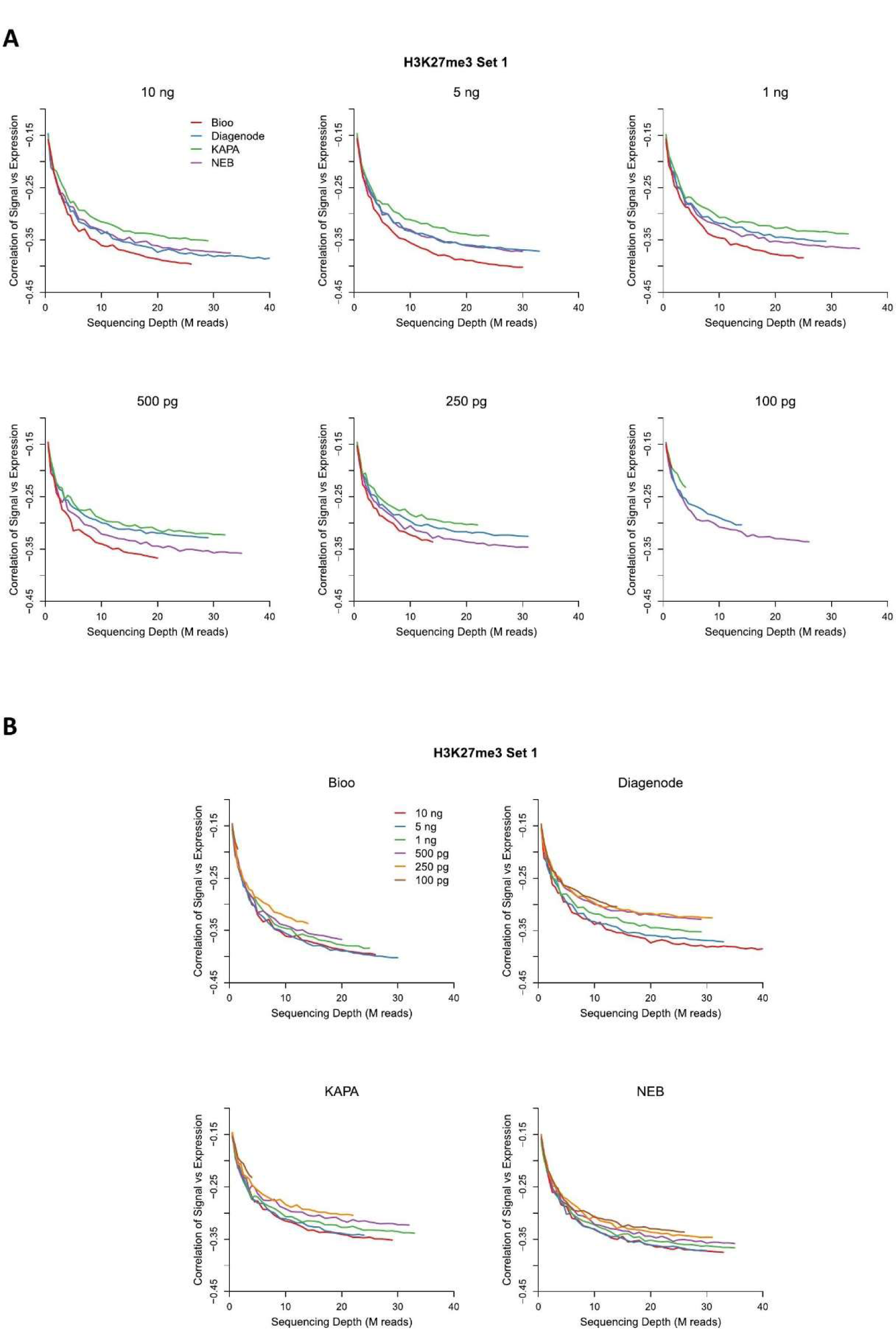
Spearman correlation of H3K27me3 signal intensity versus gene expression plotted against increasing sequencing depth for H3K27me3 set 1 libraries. The lower the curve, the better the negative correlation with gene expression. **A.** Graphs displayed to highlight diffences in the performance of each protocol at specific DNA inputs. **B.** Graphs displayed to facilitate comparisons of a single protocol at different DNA inputs.

#### Correlation with chromatin accessibility

As the H3K27me3 histone mark is associated with heterochromatin (14, 25), we also evaluated the correlation of H3K27me3 signal in our ChIP-Seq libraries with the signal of chromatin accessibility in LNCaP cells as determined by DNaseI-Seq data available through the UCSC genome browser (Figure 6 and S8). Overall, the negative correlation between H3K27me3 signal vs. chromatin accessibility agreed with the negative correlation with gene expression. Among the libraries we tested, KAPA libraries were the least inconsistent between H3K27me3 set 1 and set 2 when comparing H3K27me3 signal to chromatin accessibility. Among the remaining three protocols, Bioo libraries had the best negative correlation between H3K27me3 signal and chromatin accessibility, except for the 100 pg library in H3K27me3 set 1. NEB libraries always performed better than Diagenode libraries at all input DNA levels (Figure 6A and S8A). As expected, H3K27me3 ChIP-Seq using higher levels of input DNA generally had better negative correlation with chromatin accessibility than did ChIP-Seq using lower levels of input DNA (Figure 6B and S8B).

**Figure 6.**
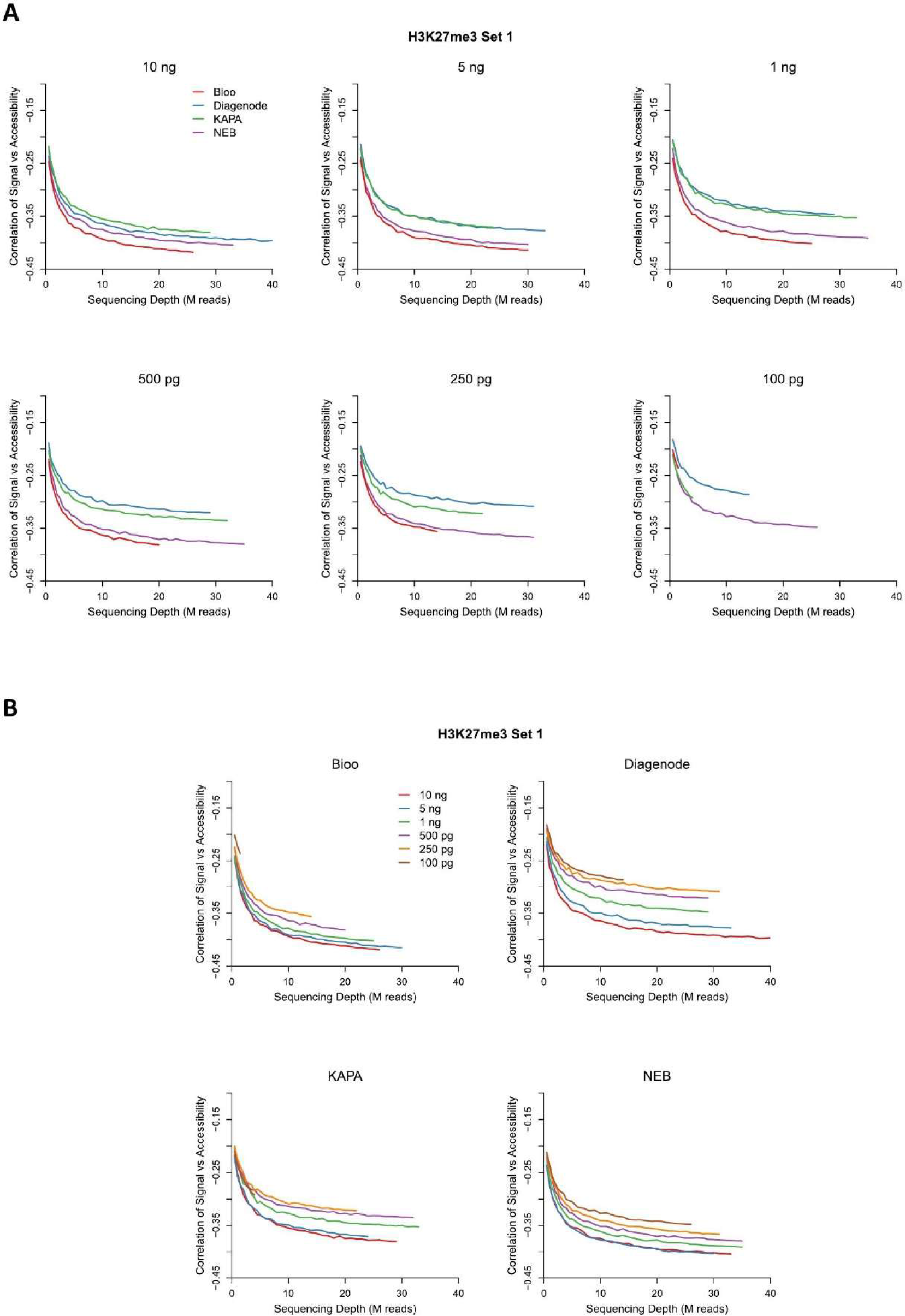
Spearman correlation of H3K27me3 signal intensity versus chromatin accessibility plotted against increasing sequencing depth for H3K27me3 set 1 libraries. The lower the curve, the better the negative correlation with gene expression. **A.** Graphs displayed to highlight diffences in the performance of each protocol at specific DNA inputs. **B.** Graphs displayed to facilitate comparisons of a single protocol at different DNA inputs.

### CTCF ChIP-Seq

To evaluate the performance of the four protocols for determining the genome-wide distribution of the transcription factor CTCF, we assessed the number of peaks called, the percentage of peaks that contained the canonical CTCF binding motif, and the distance between the CTCF motif and the peak summit (Figure 7). Overall the Diagenode libraries always yielded a greater number of called peaks, and the NEB libraries yielded a lesser number of peaks for all CTCF sets (Figure 7A). With respect to identifying a CTCF motif within a peak, the Bioo libraries had the highest percentage of peaks lacking a CTCF motif, whereas the Diagenode libraries had the highest percentage of peaks containing a CTCF motif (Figure 7B). Consistently, the offset between a CTCF motif and the peak summit was widest for the Bioo libraries and narrowest for the Diagenode libraries (Figure 7C). These findings indicate that the Diagenode libraries might more accurately capture the genome-wide distribution of CTCF, and potentially other transcription factors, whereas the Bioo libraries might less accurately reflect the true distribution of CTCF.

**Figure 7.**
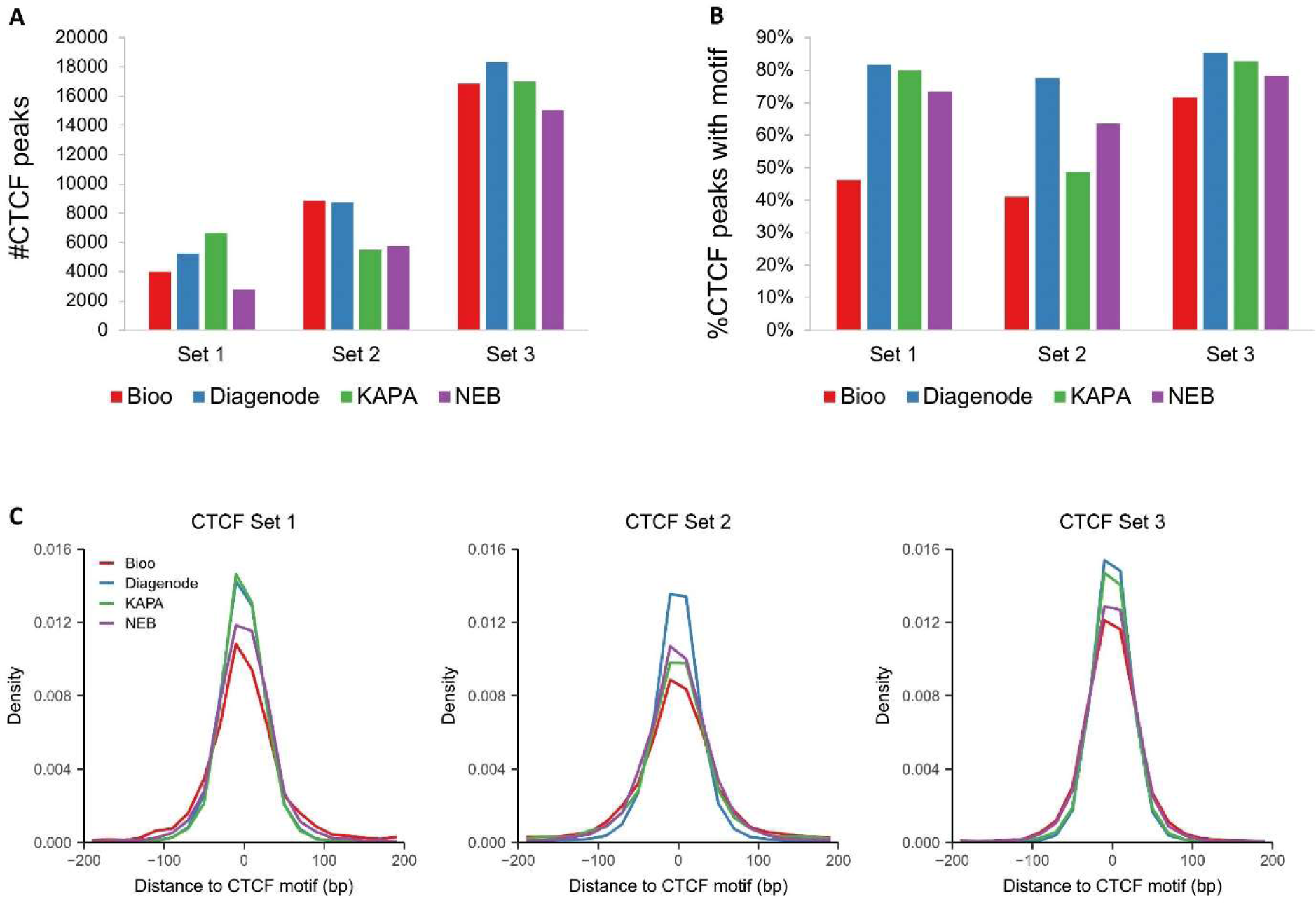
Performance of protocols for CTCF ChIP-Seq. Libraries for CTCF sets 1-3 are presented. **A.** Number of peaks called in each library. **B.** Percentage of peaks with CTCF motifs within 100 bp of peak summits. **C.** Histogram of distance between peak summit and CTCF motif.

### Reproducibility

To evaluate the reproducibility of the ChIP-Seq protocols, we calculated either the percentage of top peaks (H3K4me3) or enriched 20 kb windows (H3K27me3) shared between two libraries created from the same protocol but differing by an incremental change in the amount of input DNA (i.e. 10 ng vs. 5 ng, 5 ng vs. 1 ng, 1 ng vs. 500 pg, 500 pg vs. 250 pg, and 250 pg vs. 100 pg) and then compared these across protocols (Figure 8 and S9). As observed in many of our analyses, the KAPA libraries were inconsistent in set 1 and set 2 for both H3K4me3 and especially H3K27me3. Of the three remaining protocols, overall, Bioo libraries had the lowest reproducibility and the Diagenode libraries had the highest reproducibility. When we compared reproducibility between all incremental changes in DNA input for each individual protocol, we found that H3K4me3 and H3K27me3 libraries prepared at lower DNA inputs tended to have higher reproducibity (Figure 9 and S10). This may be because low input libraries captured only strong signals, which usually have high reproducibility.

**Figure 8.**
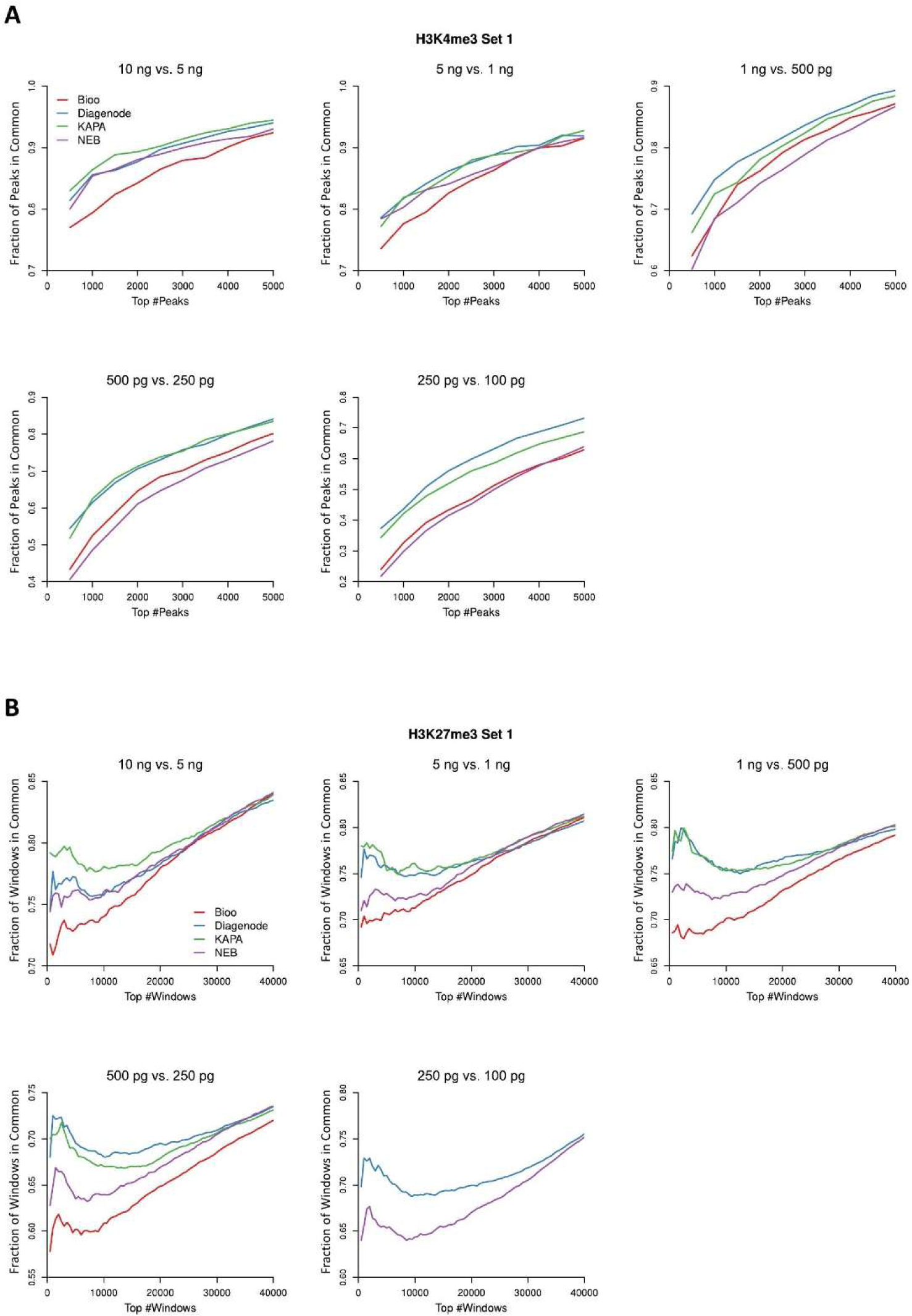
Reproducibility curves displayed to highlight differences in the performance of each protocol. The X-axis represents the number of top peaks (H3K4me3) or enriched 20 kb windows (H3K27me3) in pairwise comparisions of libraries produced from incremental changes in DNA input (i.e. 10 ng vs. 5 ng, 5 ng vs. 1 ng, 1 ng vs. 500 pg, 500 pg vs. 250 pg, and 250 pg vs. 100 pg). The Y-axis represents the fraction of top peaks or enriched windows that are in common between the two libraries that were compared. H3K4me3 (**A**) and H3K27me3 (**B**) set 1 data are plotted.

**Figure 9.**
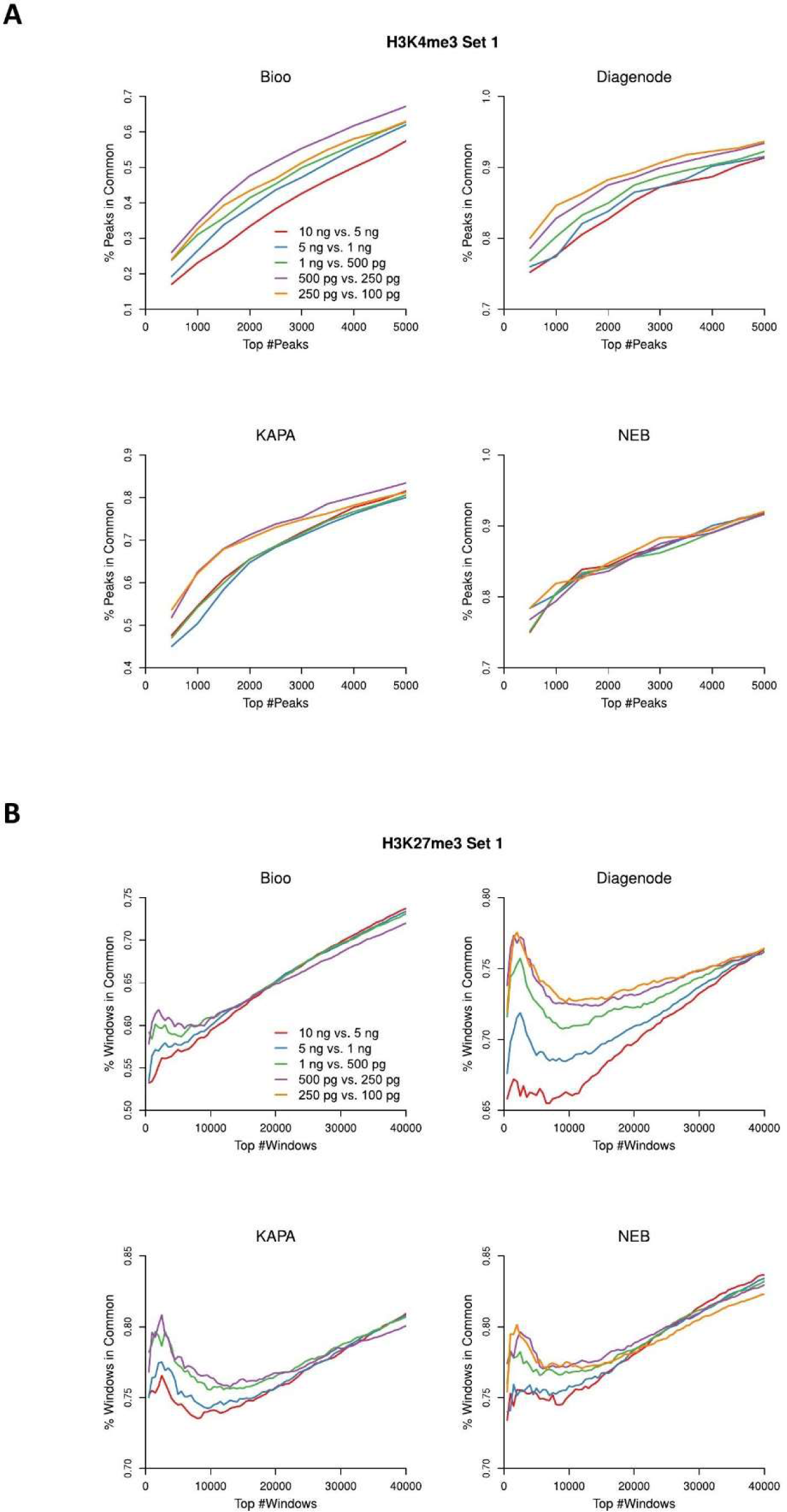
Reproducibility curves displayed to facilitate single protocol comparisons at incremental changes in DNA input. The X-axis represents the number of top peaks (H3K4me3) or enriched 20 kb windows (H3K27me3) in incremental, pair-wise comparisions of DNA input levels. The Y-axis represents the fraction of top peaks or enriched windows that are in common between the two libraries that were compared. H3K4me3 (**A**) and H3K27me3 (**B**) set 1 data are plotted.

## Discussion

### Study design

In this study, we evaluated the performance of four ChIP-Seq library preparation protocols (Bioo, Diagenode, KAPA, and NEB) on three separate targets, to represent the three typical signal enrichment patterns in ChIP-Seq experiments: sharp peaks (H3K4me3), broad domains (H3K27me3), and punctate peaks over sequences bearing a protein binding motif (CTCF). We also tested a broad range of different input DNA levels ranging from 0.10 to 10 ng for creating our H3K4me3 and H3K27me3 ChIP-Seq libraries. To ensure the reproducibility of our results, two independent sets of ChIP-Seq experiments were carried out for H3K4me3 and H3K27me3, and three independent sets of ChIP-Seq experiments were carried out for CTCF. To the best of our knowledge, our study is the most comprehensive evaluation of ChIP-Seq protocols across a wide range of input DNA amounts for three distinct protein targets associated with three different types of signal enrichment pattern in ChIP-Seq. Previous studies were either based on a single target or a smaller range of input DNA levels (4–6).

To assess our ChIP-Seq libraries, we evaluated library complexity for all libraries and reproducibility for the H3K4me3 and H3K27me3 libraries. Based on the known enrichment patterns for our three protein targets, we employed different quality metrics for each: H3K4me3 libraries were evaluated by the fraction of reads in peaks (FRiP), the fraction of reads in promoter regions, and the precision and sensitivity of peak calling; H3K27me3 libraries were evaluated by their correlation with gene expression and chromatin accessibility; and CTCF libraries were evaluated by the number of peaks, the percentage of peaks that contained the CTCF motif, and the distance between the CTCF motif and the peak summit.

### Protocol Performance

An important lesson from our study is that no single protocol consistently outperformed the others for all protein targets across all quality measurements. Overall, the Bioo libraries had lower library complexity, lower reproducibility, and lower performance than the others for preparation of the H3K4me3 and CTCF libraries. However, Bioo libraries performed better than the others for H3K27me3 libraries. Comparatively, the NEB libraries generally had higher library complexity and better H3K4me3 signal, especially at ng DNA levels and were generally the most consistent across different DNA levels for several metrics (i.e. library complexity, signal portion in H3K4me3 libraries, and reproducibility). The Diagenode protocol performed well for preparing CTCF libraries, with a good number of peaks called, the highest percentage of peaks with the CTCF motif, and the shortest distance between the peak summit and CTCF motif. The KAPA libraries were more variable and inconsistent compared to the others particularly for library complexity at low input DNA levels, the signal portion in H3K4me3 libraries prepared at low input DNA levels, the correlation between H3K27me3 signal and gene expression/chromatin accessibility, the number of CTCF peaks, and reproducibility. A summary of each protocol’s performance based on specific quality metrics is shown in Table 2.

**Table 2.**
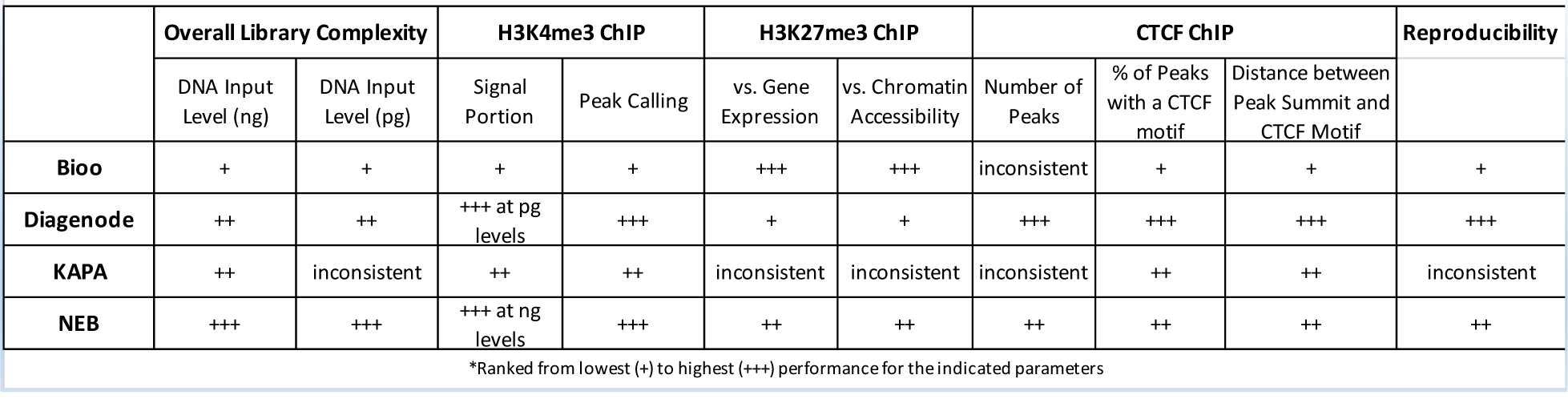
Performance of Library Preparation Protocols*.

|  | Overall Library Complexity |  | H3K4me3 ChIP |  | H3K27me3 ChIP |  | CTCF ChIP |  |  | Reproducibility |
| --- | --- | --- | --- | --- | --- | --- | --- | --- | --- | --- |
|  | DNA Input Level (ng) | DNA Input Level (pg) | Signal Portion | Peak Calling | vs. Gene Expression | vs. Chromatin Accessibility | Number of Peaks | % of Peaks with a CTCF motif | Distance between Peak Summit and CTCF Motif |  |
| **Bioo** | + | + | + | + | +++ | +++ | inconsistent | + | + | + |
| **Diagenode** | ++ | ++ | +++ at pg levels | +++ | + | + | +++ | +++ | +++ | +++ |
| **KAPA** | ++ | inconsistent | ++ | ++ | inconsistent | inconsistent | inconsistent | ++ | ++ | inconsistent |
| **NEB** | +++ | +++ | +++ at ng levels | +++ | ++ | ++ | ++ | ++ | ++ | ++ |

\*Ranked from lowest (+) to highest (+++) performance for the indicated parameters

In summary, our study indicates that commercial library preparation kits performed differently for different classes of protein targets. For example, the NEB protocol may be the best choice for H3K4me3 (and potentially other histone modifications with sharp peak enrichment), Bioo may be the best choice for H3K27me3 (and potentially other histone modifications with broad domain enrichment) though not at very low DNA levels, and Diagenode may be the best choice for CTCF (and potentially other transcription factors that bind a specific DNA binding motif). For ChIP-Seq experiments that target proteins with unknown signal enrichment patterns, the NEB protocol might be a good choice as it performed well with all three targets and the NEB libraries produced consistent results across different input DNA levels.

### Performance of different input DNA levels and potential problems of some quality metrics

Although libraries prepared using higher levels of input DNA were expected to have better quality (because of more representative DNA material and fewer PCR cycles), only the following metrics increased with increasing DNA input: library complexity, peak calling precision and sensitivity for H3K4me3, and correlation vs. gene expression/chromatin sensitivity for H3K27me3. In contrast, the fraction of reads in peaks (FRiP) and the fraction of reads in promoter regions for H3K4me3, and reproducibility for both H3K4me3 and H3K27me3 decreased with increasing DNA input. A previous study from Sundaram et al. also reported an inverse correlation between H3K4me3 signal over TSSs and DNA input (6). To further investigate why some metrics inversely correlated with input DNA level, we generated scatter plots of H3K4me3 signal intensity in peaks for each input level vs. the 10 ng level (Figure S11). Our results suggested that the signal from libraries created with lower amounts of DNA was skewed toward stronger peaks and may explain why the calculated signal portion in low-input libraries tended to be higher than in high-input libraries. Among all protocols, NEB was less skewed toward strong peaks. Given that the NEB protocol also gave the highest portion of H3K4me3 signal in libraries at 10 and 5 ng, we also compared the other protocols to the NEB protocol at 10 and 5 ng levels using scatter plots of H3K4me3 signal intensity in peaks (Figure S12). Interestingly, we found that instead of being skewed toward strong peaks, the signal from the NEB libraries increased more evenly compared to the others. That is, the signal in weak peaks was also stronger. Together, our data indicate that some metrics such as FRiP that are commonly used to evaluate the quality of ChIP-Seq may not always be reliable. It is possible that samples with better on-target specificity, such as the 10 ng level samples in our study, receive a worse value because the sample captures real but weak signals that are incorrectly classified as background noise. Indeed, in contrast to the H3K4me3 signal around TSSs, higher input DNA levels tended to have stronger signals than lower input levels around enhancers (Figure S13), where H3K4me3 is weakly enriched (1, 26). Based on our investigation, we suggest examining multiple metrics to determine ChIP-Seq library quality, and include an additional visual verification such as a signal intensity scatter plot as used in this study.

In summary, our study confirmed the common belief that the libraries prepared using higher levels of input DNA have a better quality.

### Effect of PCR cycles on our study

In this study, different numbers of PCR cycles were used to amplify DNA fragments in ChIP-Seq libraries according to the manufacturer’s recommendation, with Bioo requiring the most cycles followed by, Diagenode, KAPA and NEB [Table 1]. We chose to follow the manufacturer’s recommendations as we anticipate that is what most users would do and therefore, makes our results more meaningful for their experiments. However, PCR introduces bias that can affect the metrics we used to evaluate the ChIP-Seq library quality. For example, library complexity was directly impacted by the number of PCR cycles and correlated perfectly with the number of PCR cycles. On the other hand, none of the other quality metrics we used had a perfect correlation with the number of PCR cycles. Although PCR is regarded as the most important cause of GC bias (27) and GC content generally increases with cycle number (28), the Bioo libraries, which underwent the highest number of PCR cycles, tended to have lower GC content (Figure S14). Our results suggest that although the quality of the ChIP-Seq library can be influenced by the number of PCR cycles, it is more influenced by other factors intrinsic to each specific protocol.

### Limitations and future work

There are limitations to this study that could be addressed in the future. For example, here we only investigated the representative protein targets of two major regulatory mechanisms: transcription factors and posttranslational modifications of histones. In the future, we may investigate the third major mechanism: higher-order chromatin organization, that might be addressed by targeting nuclear lamina proteins (29). Lamina-associated domains range from 80 kb to 30 Mb in human fibroblasts. These domains are even broader than those associated with H3K27me3, thus the protocols’ performance on lamina-associated domains may be different from our current study. Future work should also include using paired-end sequencing data to help determine whether the preference to a range of fragment size is a factor influencing the protocol performance.

## Supporting information

full resolution Figure 2

full resolution Figure 3

full resolution Figure 4

full resolution Figure 5

full resolution Figure 6

full resolution Figure 7

full resolution Figure 8

full resolution Figure 9

## Abbreviations

ChIP-Seq: Chromatin immunoprecipitation followed by high-throughput sequencing
CTCF: CCCTC-binding factor, a highly conserved zinc finger protein and a transcription factor
TF: transcription factor
PBS: Phosphate-buffered saline
SDS: Sodium dodecyl sulfate
FPKM: Number of Fragments Per Kilobase per Million fragments
EDD: Enriched domain detector
RNA-Seq: Ribonucleic acid sequencing
RPKM: Reads per kilobase per million mapped reads
TSS: Transcription start site
NRF: Non-Redundant Fraction
FRiP: Fraction of reads in peaks
LNCaP: Androgen-sensitive human prostate adenocarcinoma cells

## Acknowledgements

We thank Dr. Briana Dennehey for editorial assistance, Dr. Sharon Dent and Dr. Michael MacLeod for their critical reading of the manuscript, and Joi Holcomb for her help with the preparation of figure 1.

## Authors’ contributions

JS, YL, MSS and LDC conceived the project, designed experiments, interpreted data and wrote the manuscript; MSS, LDC, CDM, YT, MWT, DZ performed experiments; YL, SG and KL conducted bioinformatics analyses; ME and DGT helped with data interpretation; SG helped with writing the manuscript. All authors read and approved the manuscript.

## Availability of data and materials

The raw datasets for the ChIP-Seq protocols have been deposited in GEO and can be accessed by the accession number GSExxx.

## Ethics approval and consent to participate

Not applicable.

## Consent for publication

Not applicable

## Figure legends

**Figure S1.** Representative screenshots showing the overall signal landscape for each ChIP-Seq library. **A.** H3K4me3 Set 1 libraries. **B.** H3K4me3 Set 2 libraries. **C.** H3K27me3 Set1 libraries. **D.** H3K27me3 Set2 libraries. **E.** CTCF libraries. The small bars on top of each track indicate the peaks/broad domains called in each library. The number of reads used to generate the signal landscape of each library is indicated at the end of track name.

**Figure S2.** Library complexity measured as Non-Redundant Fraction (NRF). **A.** NRF for H3K4me3 set 2 libraries. **B.** NRF for H3K27me3 set 2 libraries.

**Figure S3.** FRiP (fraction of reads in peaks) for H3K4me3 plotted against sequencing depth for H3K4me3 set 2 libraries. **A.** Graphs displayed to highlight differences in protocol performance at specific DNA inputs. **B.** Graphs displayed to facilitate comparisons of a single protocol at different DNA inputs.

**Figure S4.** Fraction of H3K4me3 reads in promoter regions. **A-B.** Bar graphs displayed to highlight differences in protocol performance at specific DNA inputs. **C-D.** Graphs displayed to facilitate comparisons of a single protocol at different DNA inputs. H3K4me3 set 1 libraries are presented in A and C, and H3K4me3 set 2 libraries are presented in B and D. Figure S5. Aggregated signal curves around transcription start sites (TSSs) for H3K4me3 libraries organized by protocol.

**Figure S6.** The peak calling quality for H3K4me3 measured as the percentage of peaks in promoter regions vs. the percentage of expressed genes marked by peaks at varying sequencing depth for H3K4me3 set 2 libraries. **A.** Graphs displayed to highlight diffences in the performance of each protocol at specific DNA inputs. **B.** Graphs displayed to facilitate comparisons of a single protocol at different DNA inputs.

**Figure S7.** Spearman correlation of H3K27me3 signal intensity versus gene expression plotted against increasing sequencing depth for H3K27me3 set 2 libraries. The lower the curve, the better the negative correlation with gene expression. **A.** Graphs displayed to highlight diffences in the performance of each protocol at specific DNA inputs. **B.** Graphs displayed to facilitate comparisons of a single protocol at different DNA inputs.

**Figure S8.** Spearman correlation of H3K27me3 signal intensity versus chromatin accessibility plotted against increasing sequencing depth for H3K27me3 set 2 libraries. The lower the curve, the better the negative correlation with gene expression. **A.** Graphs displayed to highlight diffences in the performance of each protocol at specific DNA inputs. **B.** Graphs displayed to facilitate comparisons of a single protocol at different DNA inputs.

**Figure S9.** Reproducibility curves displayed to highlight differences in the performance of each protocol. The X-axis represents the number of top peaks (H3K4me3) or enriched 20 kb windows (H3K27me3) in pairwise comparisions of libraries produced from incremental changes in DNA input (i.e. 10 ng vs. 5 ng, 5 ng vs. 1 ng, 1 ng vs. 500 pg, 500 pg vs. 250 pg, and 250 pg vs. 100 pg). The Y-axis represents the fraction of top peaks or enriched windows that are in common between the two libraries that were compared. H3K4me3 (**A**) and H3K27me3 (**B**) set 2 data are plotted.

**Figure S10.** Reproducibility curves displayed to facilitate single protocol comparisons at incremental changes in DNA input. The X-axis represents the number of top peaks (H3K4me3) or enriched 20 kb windows (H3K27me3) in incremental, pair-wise comparisions of DNA input levels. The Y-axis represents the fraction of top peaks or enriched windows that are in common between the two libraries that were compared. H3K4me3 (**A**) and H3K27me3 (**B**) set 2 data are plotted

**Figure S11.** Scatter plots showing H3K4me3 signal intensity in peaks for each amount of input DNA (y-axis) compared to 10 ng of input DNA (x-axis) for each protocol. H3K4me3 set 1 (**A**) and set 2 (**B**) library data are presented as scatter plots with smoothed color density, and with red lines representing LOESS-fitted curves.

**Figure S12.** Scatter plots showing H3K4me3 signal intensity in peaks for each protocol (y-axis) compared to the NEB protocol (x-axis) at DNA inputs of 10 and 5 ng. H3K4me3 set 1 (**A**) and set 2 (**B**) library data are presented as scatter plots with smoothed color density, and with red lines representing LOESS-fitted curves.

**Figure S13.** Aggregated signal curves around enhancers for H3K4me3 organized by protocol.

**Figure S14.** Bar graphs indicating the average GC content of the DNA sequences under peaks uniquely called to each protocol at specific DNA inputs. H3K4me3 set 1 and set 2 libraries are presented in **A** and **B**, respectively. H3K27me3 set 1 and set 2 libraries are presented in **C** and **D**, respectively. H3K27me3 Bioo 100 ng and KAPA 100 ng libraries didn’t have enough reads for a meaningful peak calling, thus were skipped.

