## Supplementary figures and images for "Commercial ChIP-Seq library preparation kits performed differently for different classes of protein targets"

### full resolution Figure 2

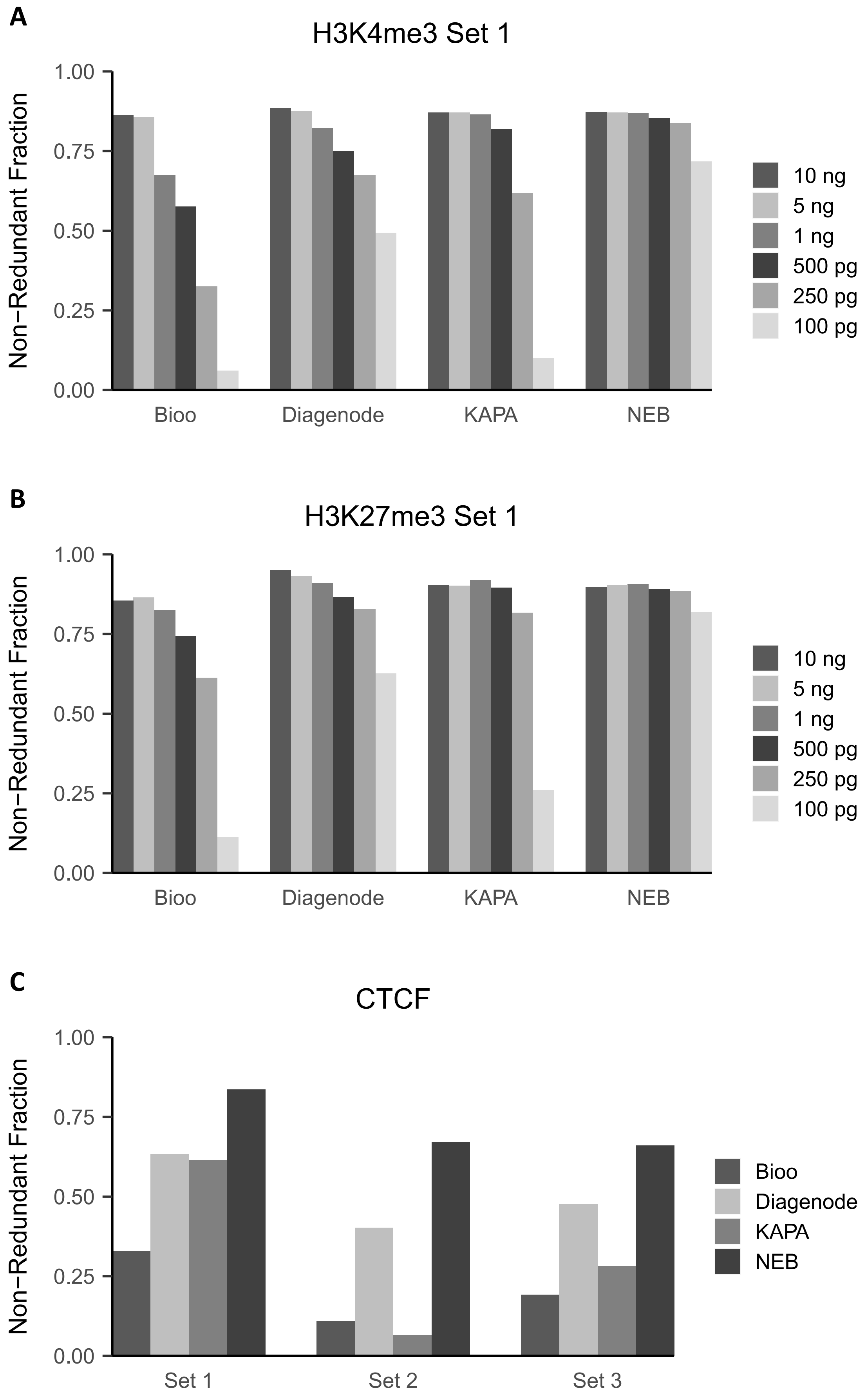

### full resolution Figure 3

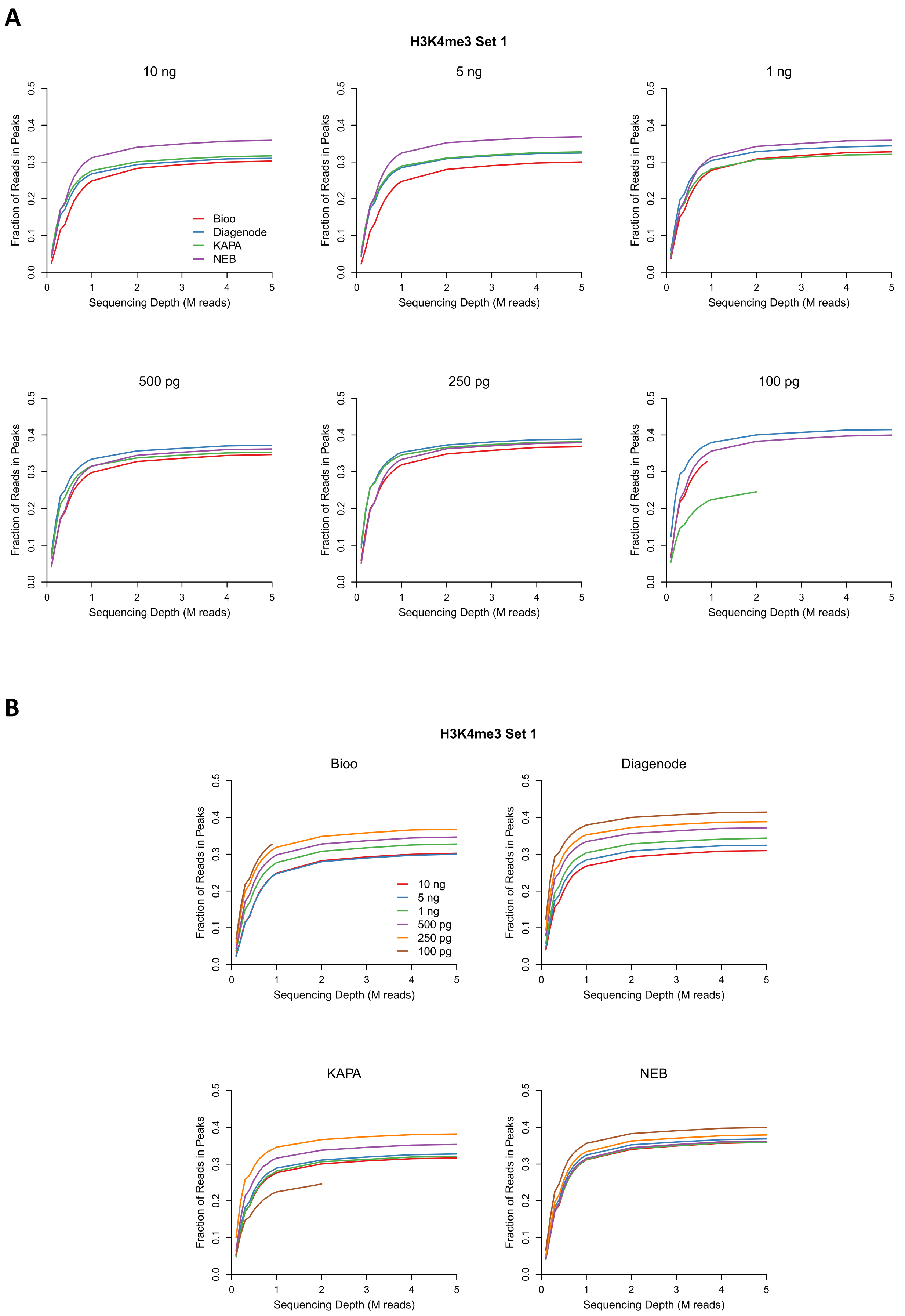

### full resolution Figure 4

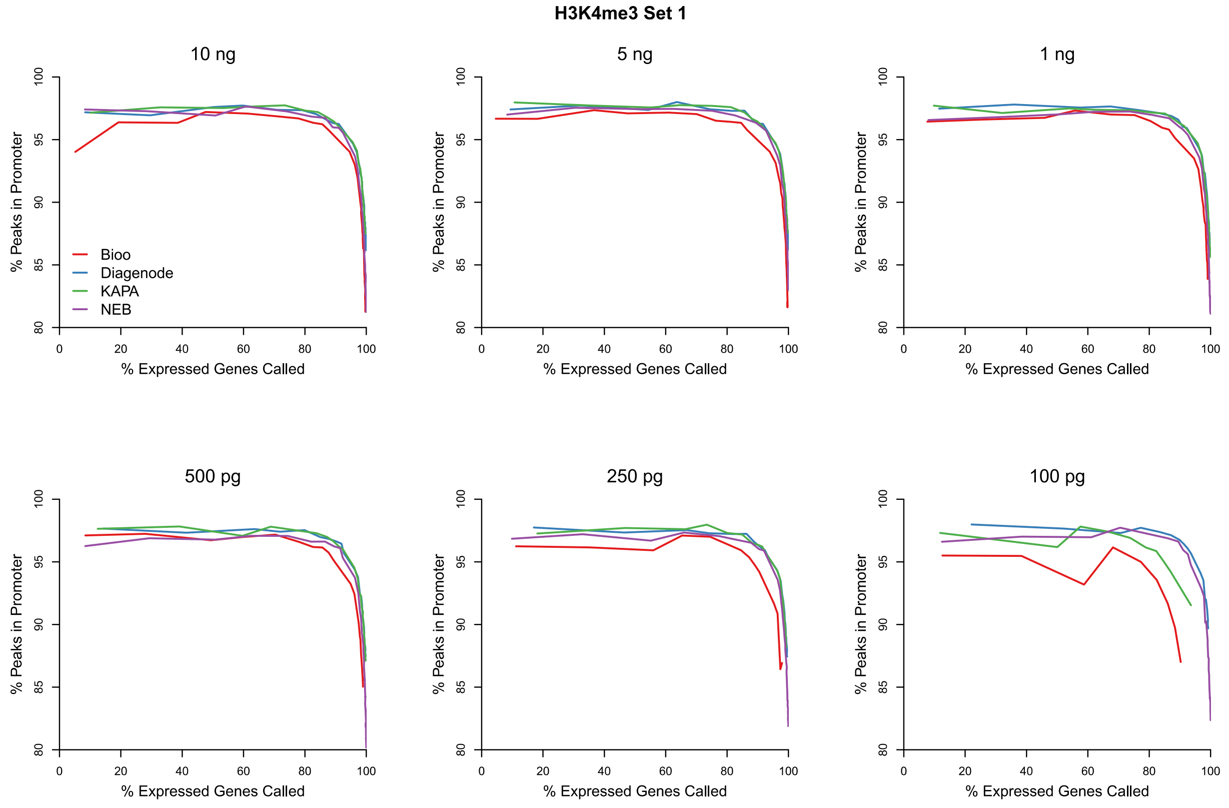

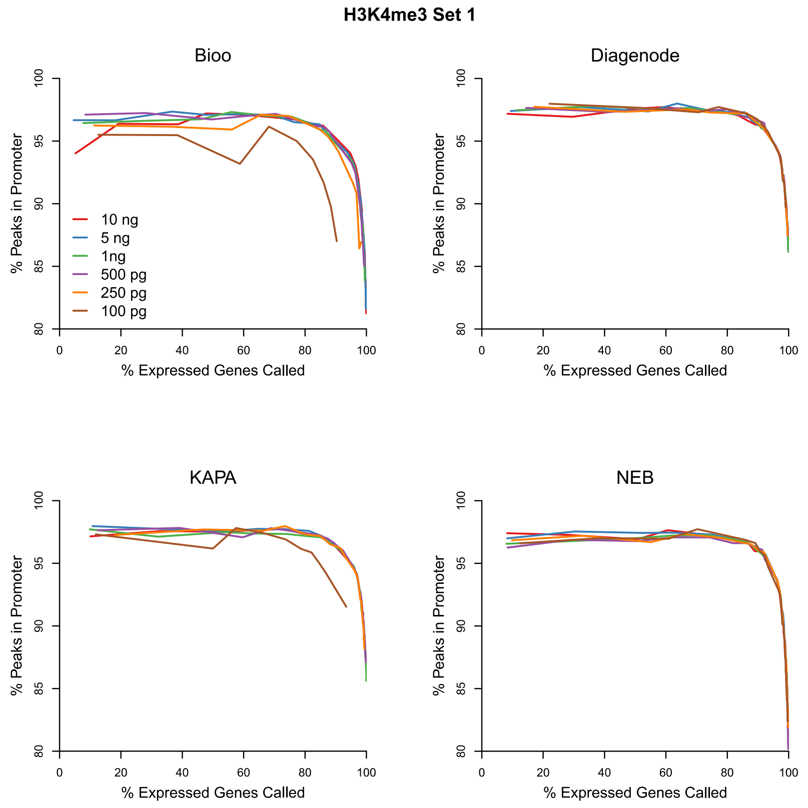

### full resolution Figure 5

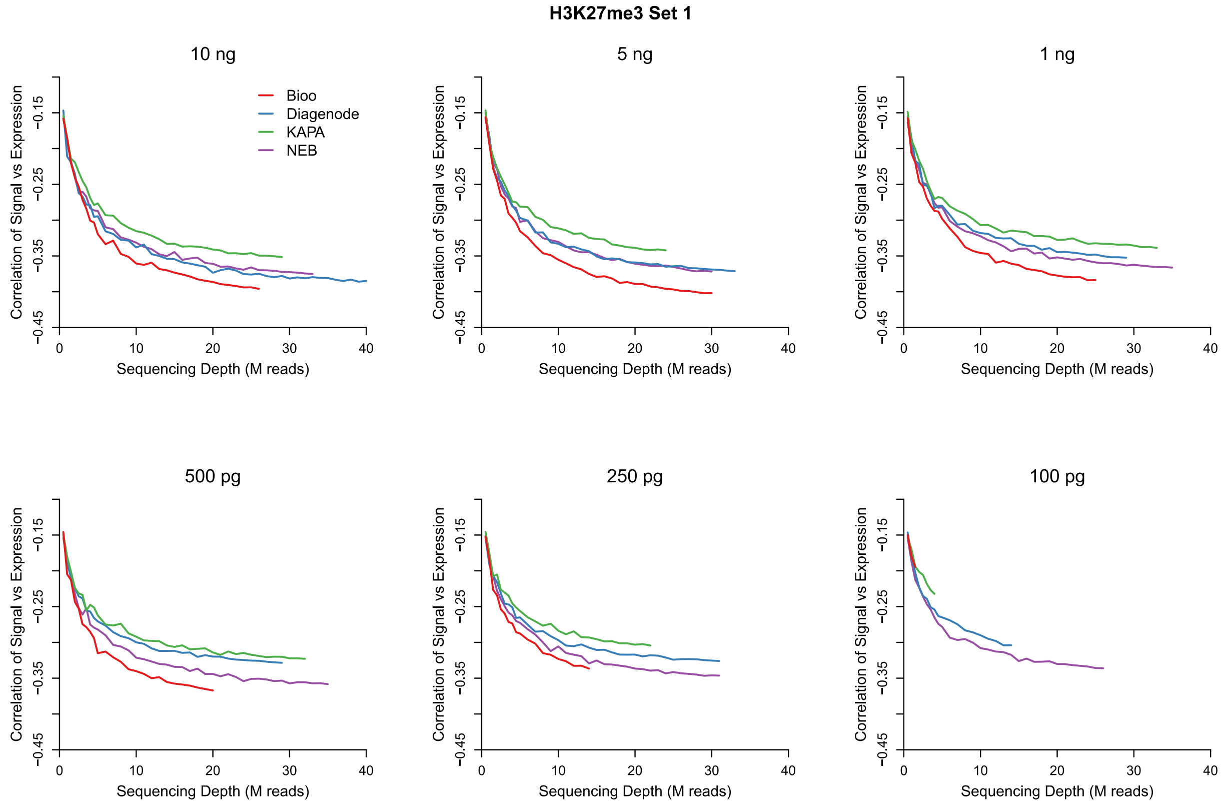

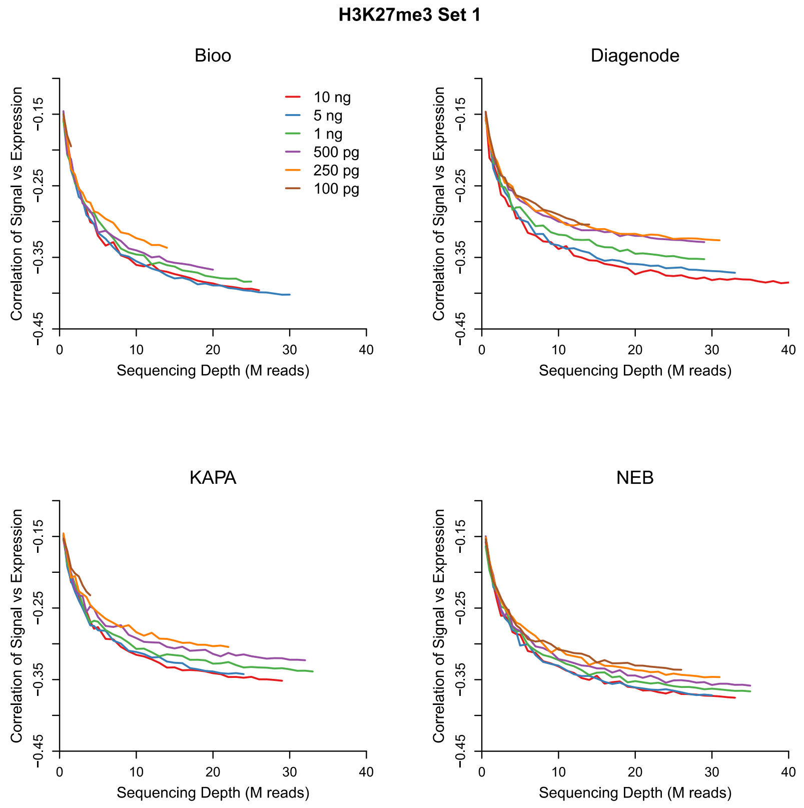

### full resolution Figure 6

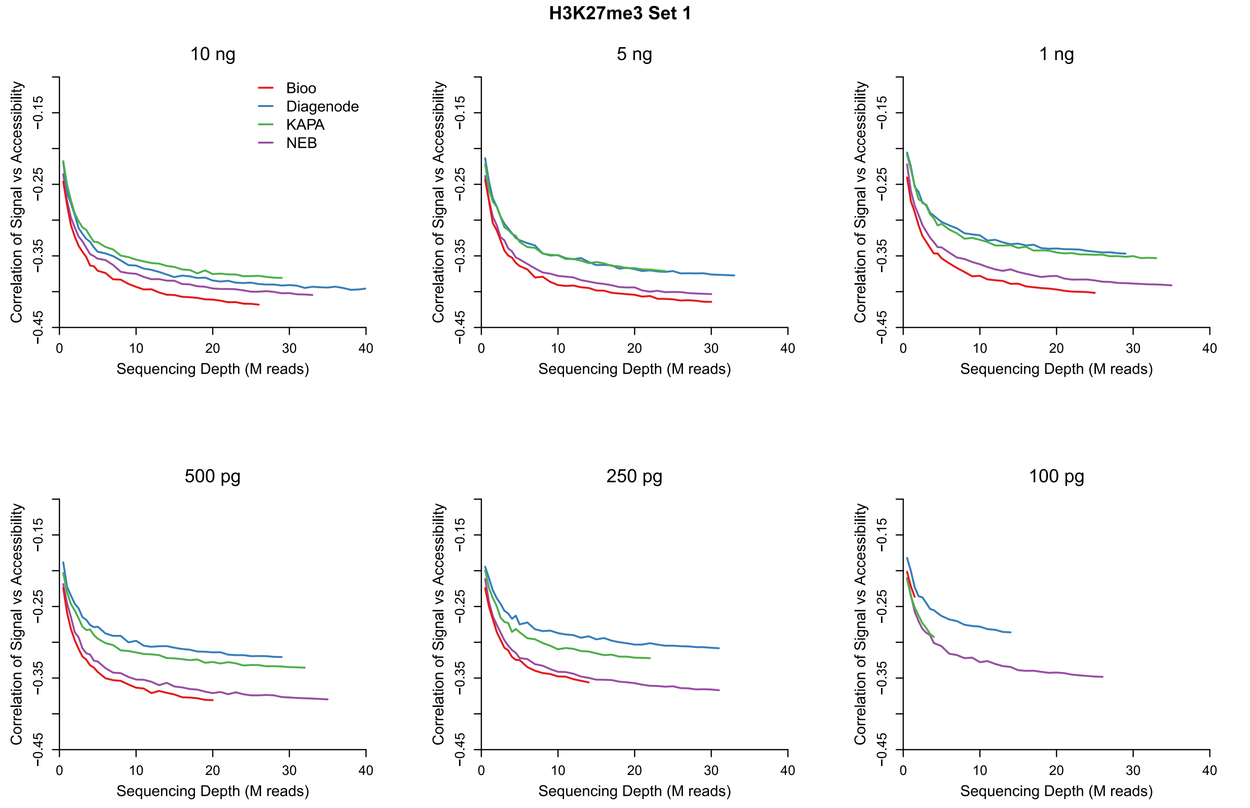

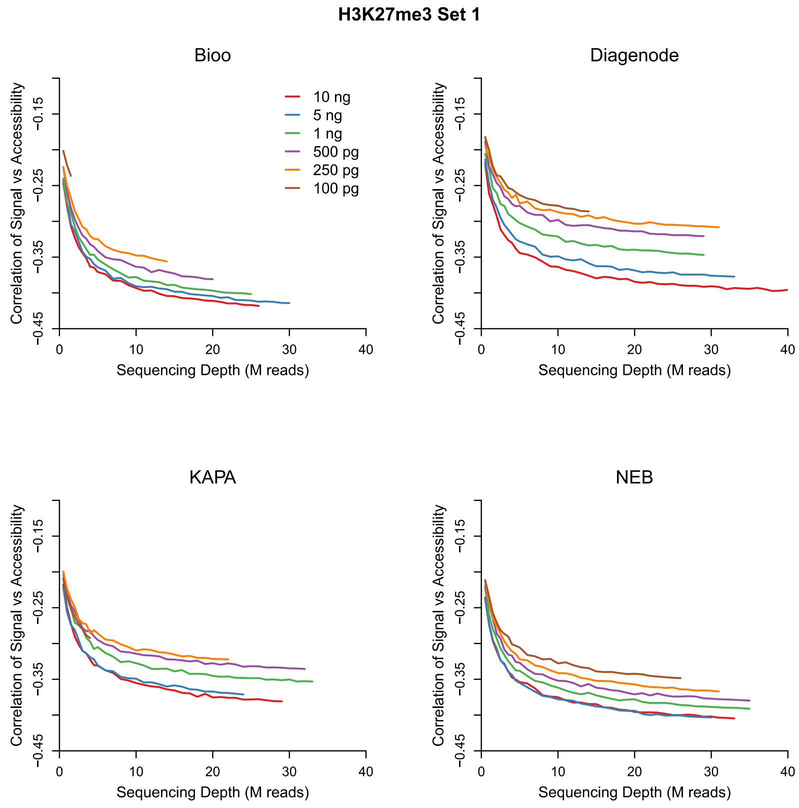

### full resolution Figure 7

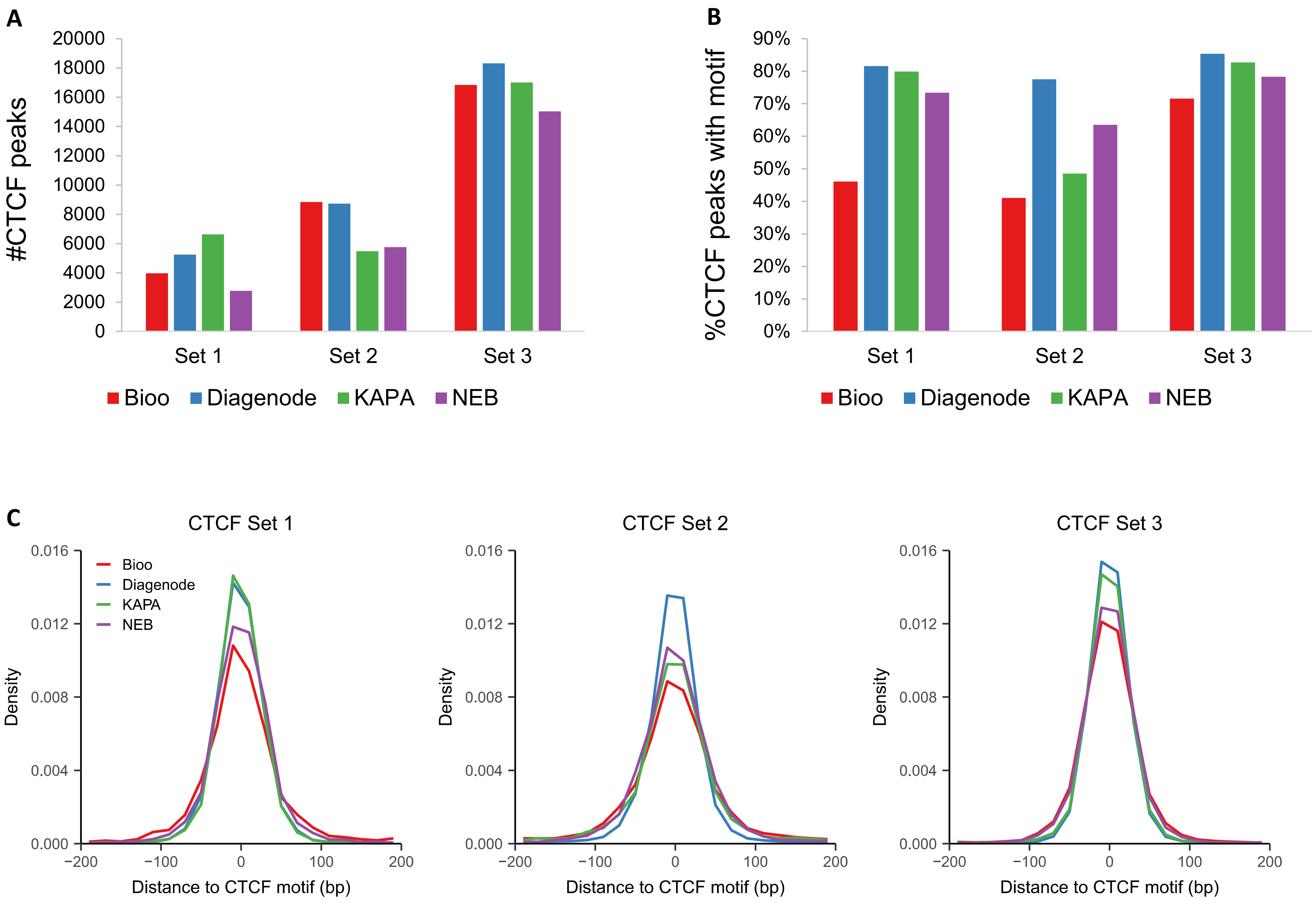

### full resolution Figure 8

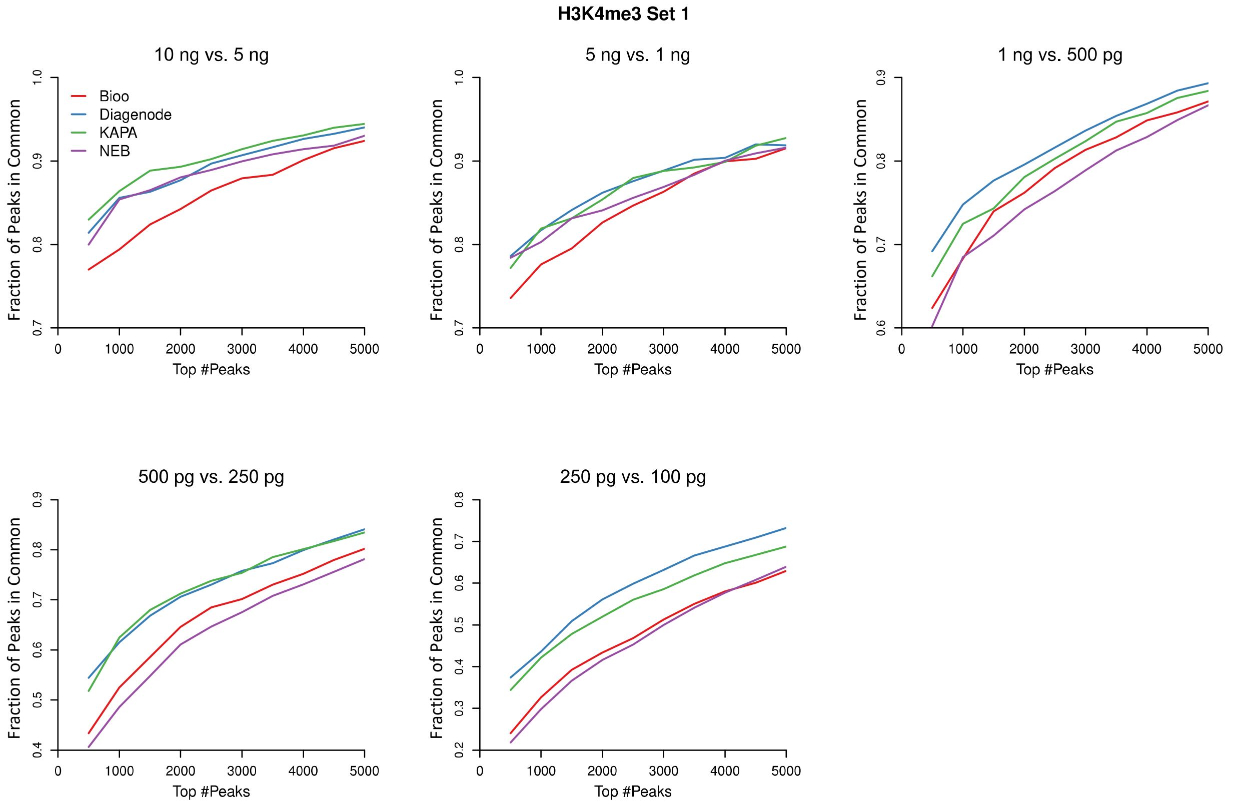

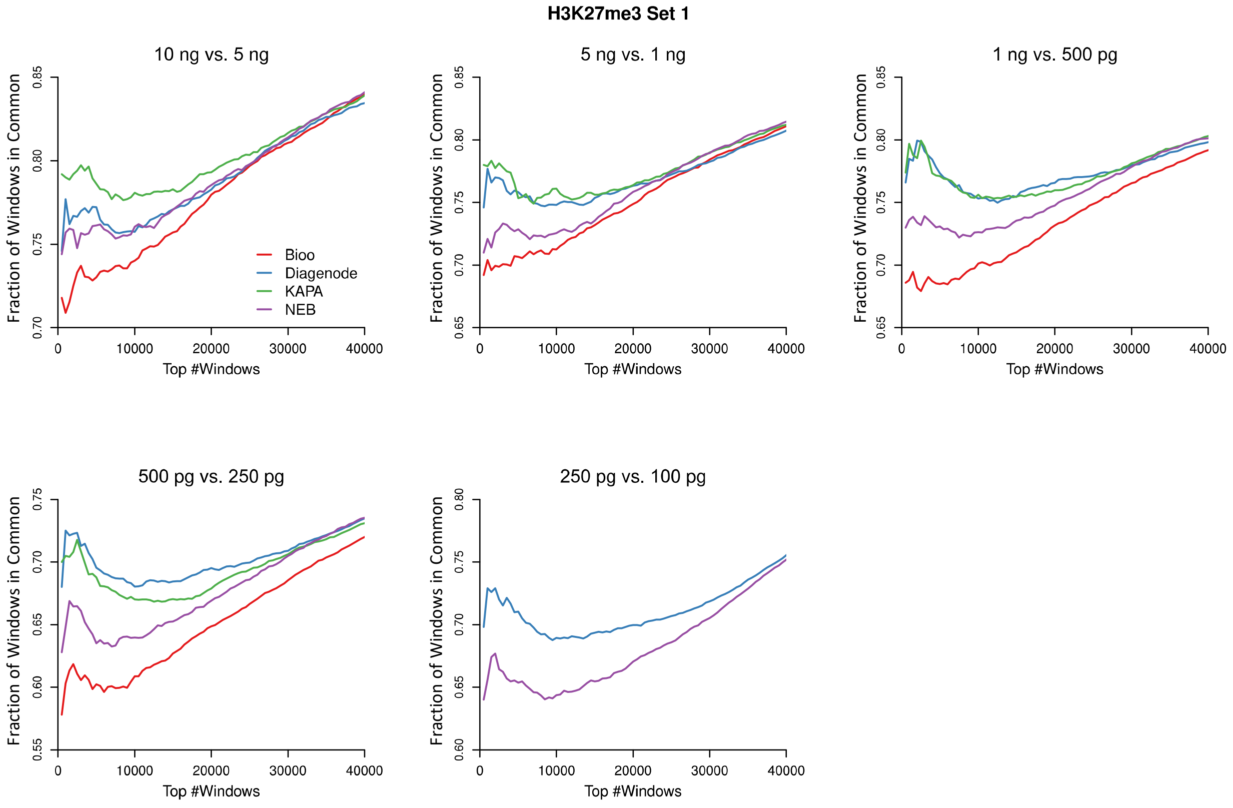

### full resolution Figure 9

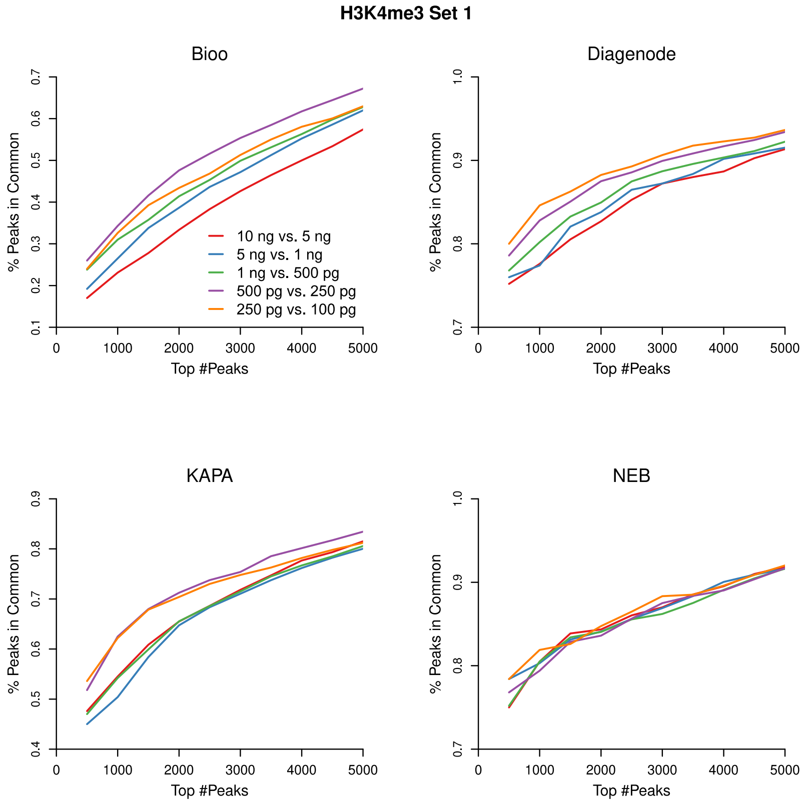

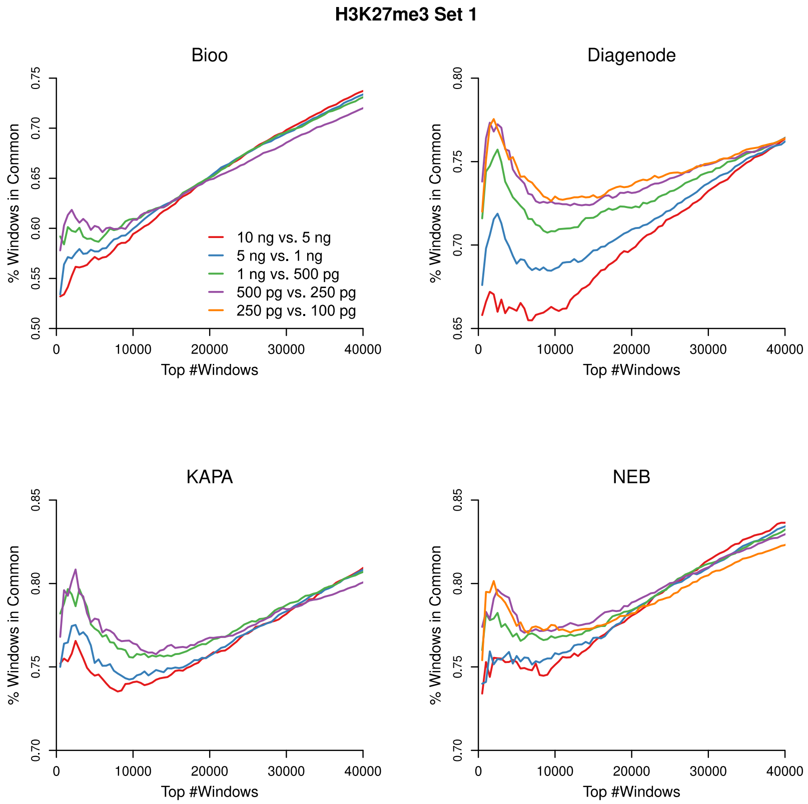

### Supplemental Figure 13

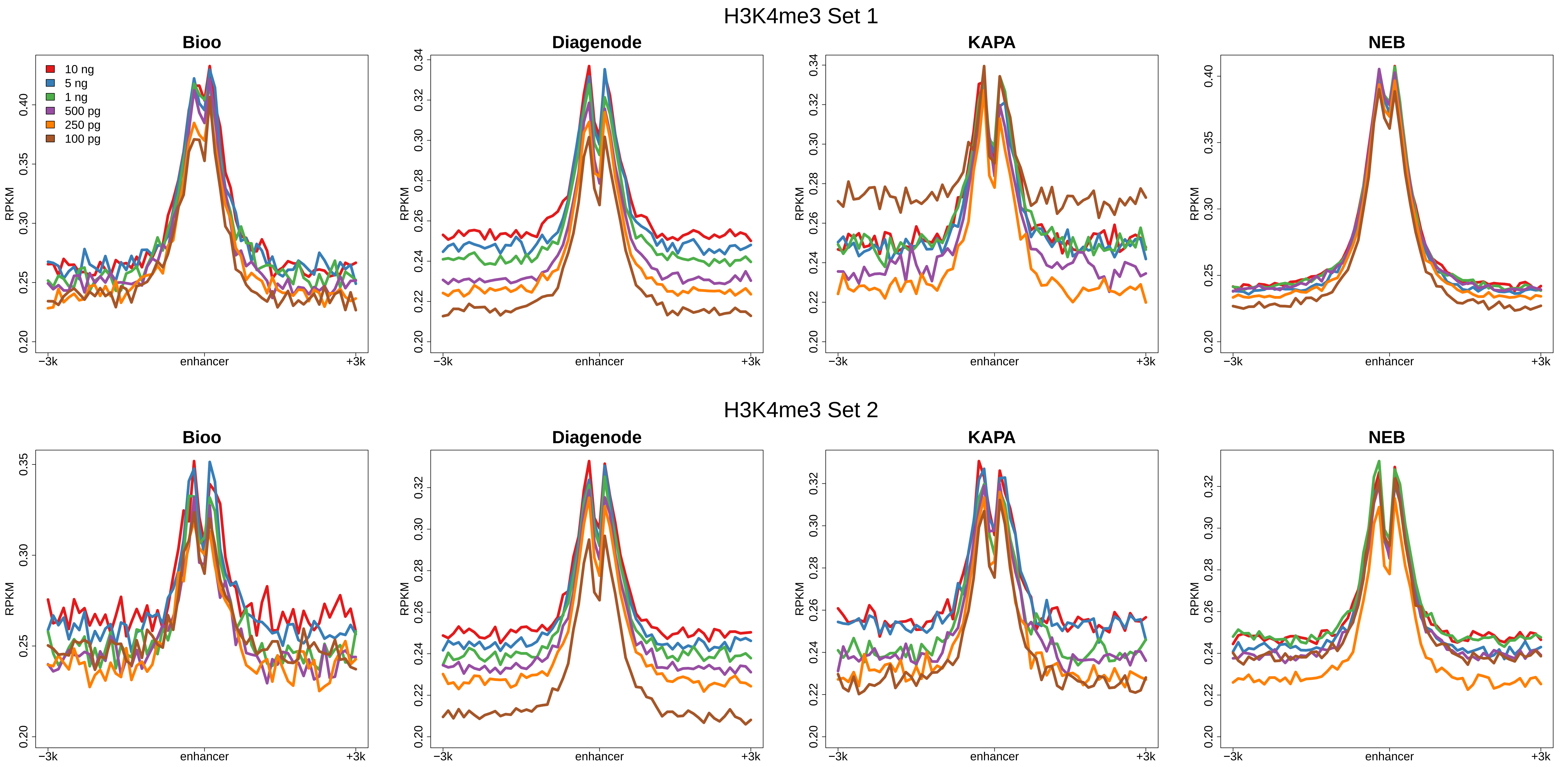

**Figure S13.** Aggregated signal curves around enhancers for H3K4me3 organized by protocol.
