## Supplemental Figure 1 for "Commercial ChIP-Seq library preparation kits performed differently for different classes of protein targets"

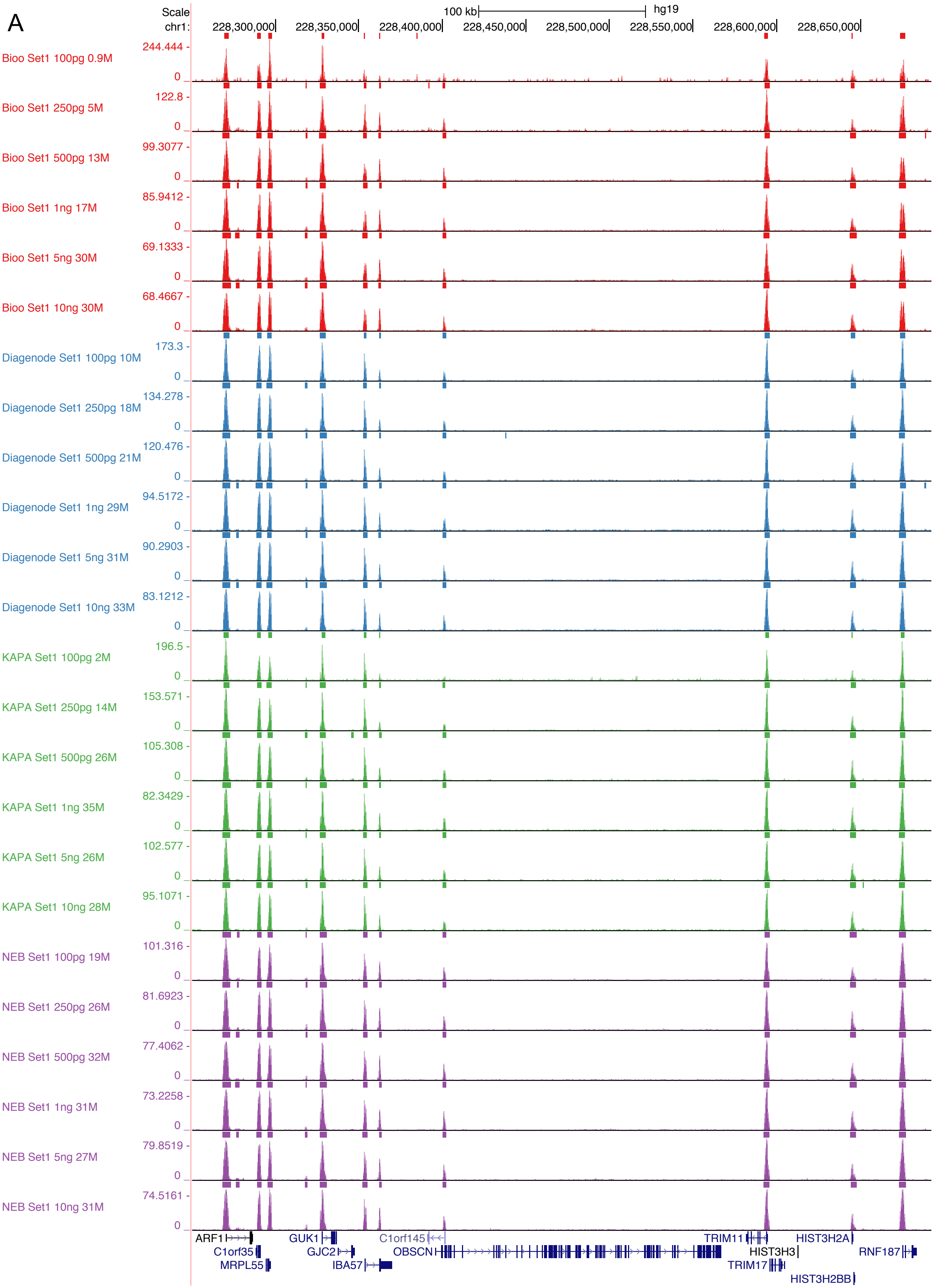
A
Scale
chr1:
100 kb
hg19
228,300,000
228,350,000
228,400,000
228,450,000
228,500,000
228,550,000
228,600,000
228,650,000
Bioo Set1 100pg 0.9M
244.444 -
Bioo Set1 250pg 5M
122.8 -
Bioo Set1 500pg 13M
99.3077 -
Bioo Set1 1ng 17M
85.9412 -
Bioo Set1 5ng 30M
69.1333 -
Bioo Set1 10ng 30M
68.4667 -
Diagenode Set1 100pg 10M
173.3 -
Diagenode Set1 250pg 18M
134.278 -
Diagenode Set1 500pg 21M
120.476 -
Diagenode Set1 1ng 29M
94.5172 -
Diagenode Set1 5ng 31M
90.2903 -
Diagenode Set1 10ng 33M
83.1212 -
KAPA Set1 100pg 2M
196.5 -
KAPA Set1 250pg 14M
153.571 -
KAPA Set1 500pg 26M
105.308 -
KAPA Set1 1ng 35M
82.3429 -
KAPA Set1 5ng 26M
102.577 -
KAPA Set1 10ng 28M
95.1071 -
NEB Set1 100pg 19M
101.316 -
NEB Set1 250pg 26M
81.6923 -
NEB Set1 500pg 32M
77.4062 -
NEB Set1 1ng 31M
73.2258 -
NEB Set1 5ng 27M
79.8519 -
NEB Set1 10ng 31M
74.5161 -
0
ARF1
C1orf35
MRPL55
GUK1
GJC2
IBA57
C1orf145
OBSCN
TRIM11
HIST3H3
TRIM17
HIST3H2A
HIST3H2BB
RNF187

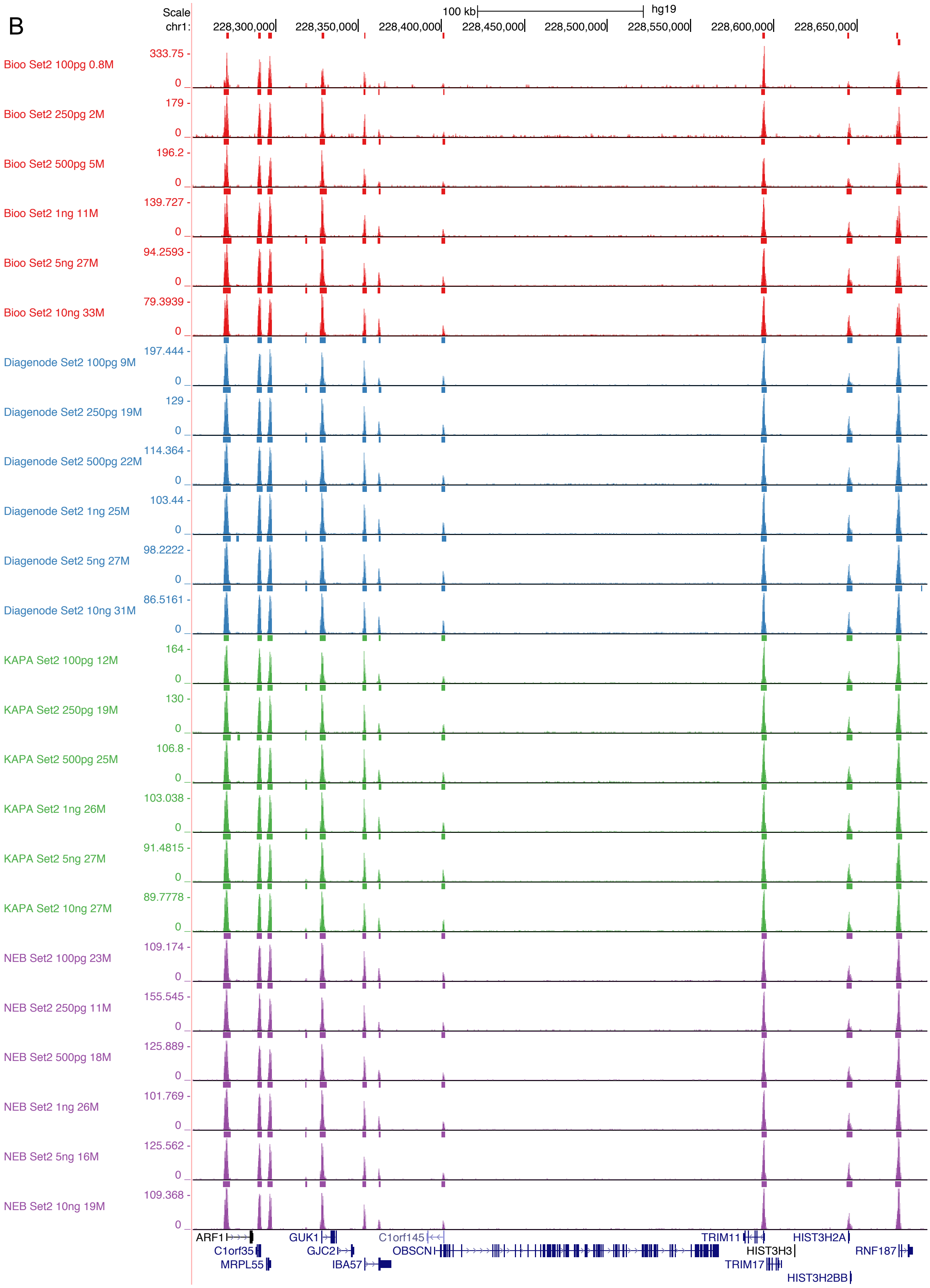
B
Scale
chr1:
100 kb
hg19
228,300,000
228,350,000
228,400,000
228,450,000
228,500,000
228,550,000
228,600,000
228,650,000
Bioo Set2 100pg 0.8M
Bioo Set2 250pg 2M
Bioo Set2 500pg 5M
Bioo Set2 1ng 11M
Bioo Set2 5ng 27M
Bioo Set2 10ng 33M
Diagenode Set2 100pg 9M
Diagenode Set2 250pg 19M
Diagenode Set2 500pg 22M
Diagenode Set2 1ng 25M
Diagenode Set2 5ng 27M
Diagenode Set2 10ng 31M
KAPA Set2 100pg 12M
KAPA Set2 250pg 19M
KAPA Set2 500pg 25M
KAPA Set2 1ng 26M
KAPA Set2 5ng 27M
KAPA Set2 10ng 27M
NEB Set2 100pg 23M
NEB Set2 250pg 11M
NEB Set2 500pg 18M
NEB Set2 1ng 26M
NEB Set2 5ng 16M
NEB Set2 10ng 19M
333.75
179
196.2
139.727
94.2593
79.3939
197.444
129
114.364
103.44
98.2222
86.5161
164
130
106.8
103.038
91.4815
89.7778
109.174
155.545
125.889
101.769
125.562
109.368
0
ARF1
C1orf35
MRPL55
GUK1
GJC2
IBA57
C1orf145
OBSCN
TRIM11
HIST3H3
TRIM17
HIST3H2A
HIST3H2BB
RNF187

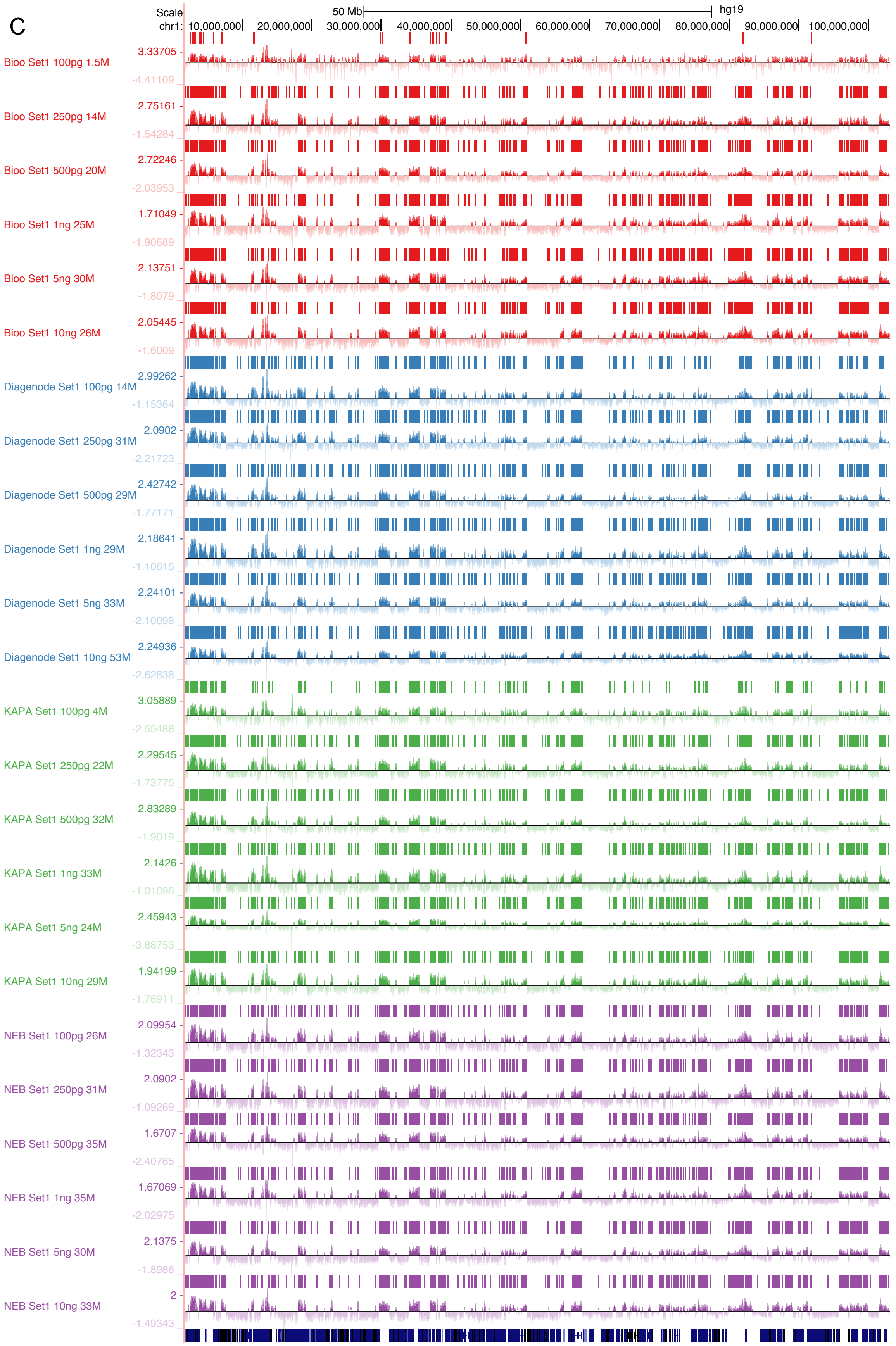

C
Scale
50 Mb
hg19
chr1: 10,000,000 20,000,000 30,000,000 40,000,000 50,000,000 60,000,000 70,000,000 80,000,000 90,000,000 100,000,000
Bioo Set1 100pg 1.5M
3.33705
-4.41109
Bioo Set1 250pg 14M
2.75161
-1.54284
Bioo Set1 500pg 20M
2.72246
-2.03953
Bioo Set1 1ng 25M
1.71049
-1.90689
Bioo Set1 5ng 30M
2.13751
-1.8079
Bioo Set1 10ng 26M
2.05445
-1.6009
Diagenode Set1 100pg 14M
2.99262
-1.15364
Diagenode Set1 250pg 31M
2.0902
-2.21723
Diagenode Set1 500pg 29M
2.42742
-1.77171
Diagenode Set1 1ng 29M
2.18641
-1.10615
Diagenode Set1 5ng 33M
2.24101
-2.10098
Diagenode Set1 10ng 53M
2.24936
-2.62838
KAPA Set1 100pg 4M
3.05889
-2.55488
KAPA Set1 250pg 22M
2.29545
-1.73775
KAPA Set1 500pg 32M
2.83289
-1.9019
KAPA Set1 1ng 33M
2.1426
-1.01096
KAPA Set1 5ng 24M
2.45943
-3.88753
KAPA Set1 10ng 29M
1.94199
-1.76911
NEB Set1 100pg 26M
2.09954
-1.32343
NEB Set1 250pg 31M
2.0902
-1.09269
NEB Set1 500pg 35M
1.6707
-2.40765
NEB Set1 1ng 35M
1.67069
-2.02975
NEB Set1 5ng 30M
2.1375
-1.8986
NEB Set1 10ng 33M
2
-1.49343

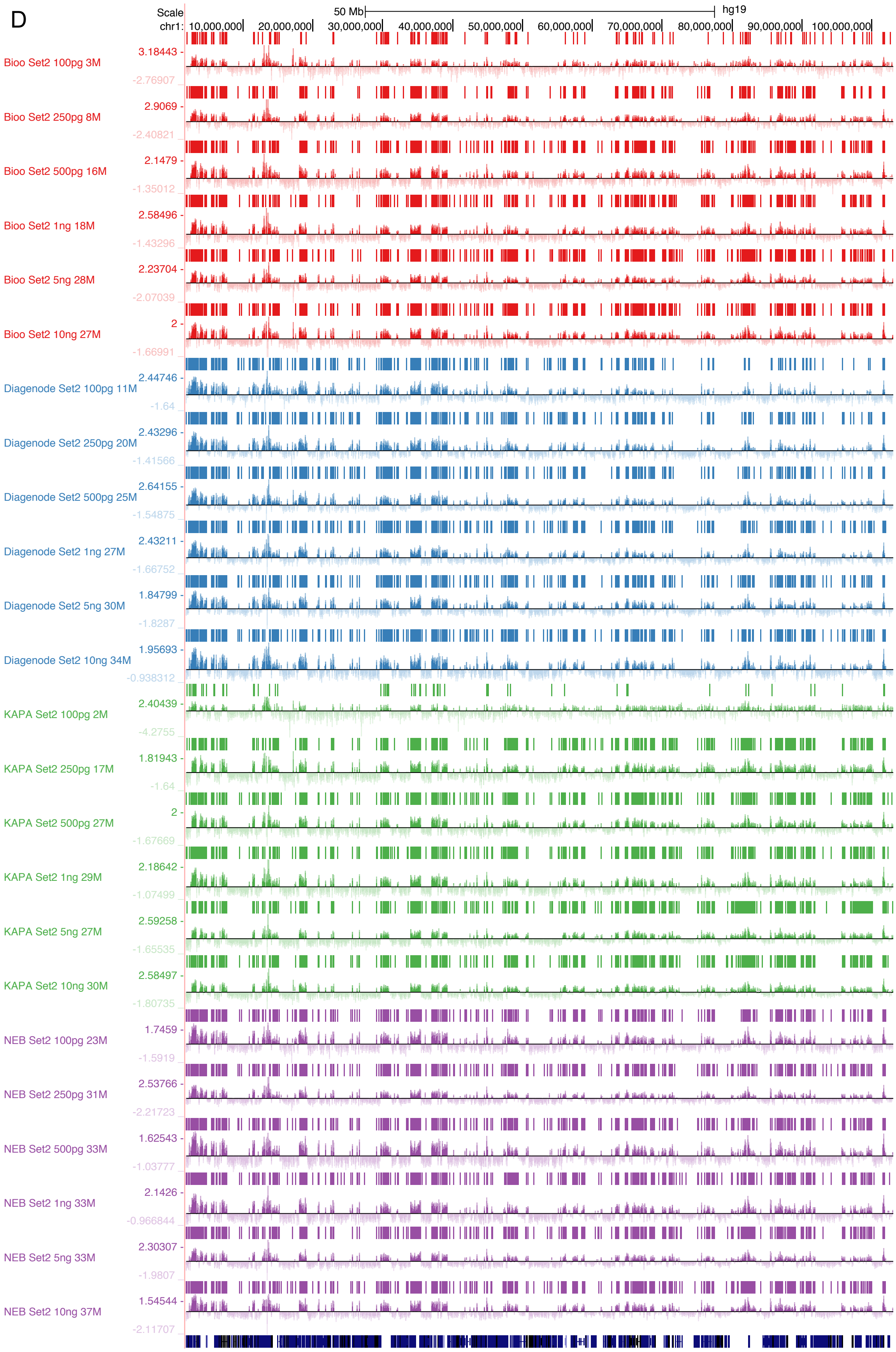

D
Scale
50 Mb
hg19
chr1: 10,000,000 20,000,000 30,000,000 40,000,000 50,000,000 60,000,000 70,000,000 80,000,000 90,000,000 100,000,000
Bioo Set2 100pg 3M
3.18443 -
-2.76907
Bioo Set2 250pg 8M
2.9069 -
-2.40821
Bioo Set2 500pg 16M
2.1479 -
-1.35012
Bioo Set2 1ng 18M
2.58496 -
-1.43296
Bioo Set2 5ng 28M
2.23704 -
-2.07039
Bioo Set2 10ng 27M
2 -
-1.66991
Diagenode Set2 100pg 11M
2.44746 -
-1.64
Diagenode Set2 250pg 20M
2.43296 -
-1.41566
Diagenode Set2 500pg 25M
2.64155 -
-1.54875
Diagenode Set2 1ng 27M
2.43211 -
-1.66752
Diagenode Set2 5ng 30M
1.84799 -
-1.8287
Diagenode Set2 10ng 34M
1.95693 -
-0.938312
KAPA Set2 100pg 2M
2.40439 -
-4.2755
KAPA Set2 250pg 17M
1.81943 -
-1.64
KAPA Set2 500pg 27M
2 -
-1.67669
KAPA Set2 1ng 29M
2.18642 -
-1.07499
KAPA Set2 5ng 27M
2.59258 -
-1.65535
KAPA Set2 10ng 30M
2.58497 -
-1.80735
NEB Set2 100pg 23M
1.7459 -
-1.5919
NEB Set2 250pg 31M
2.53766 -
-2.21723
NEB Set2 500pg 33M
1.62543 -
-1.03777
NEB Set2 1ng 33M
2.1426 -
-0.966844
NEB Set2 5ng 33M
2.30307 -
-1.9807
NEB Set2 10ng 37M
1.54544 -
-2.11707

E

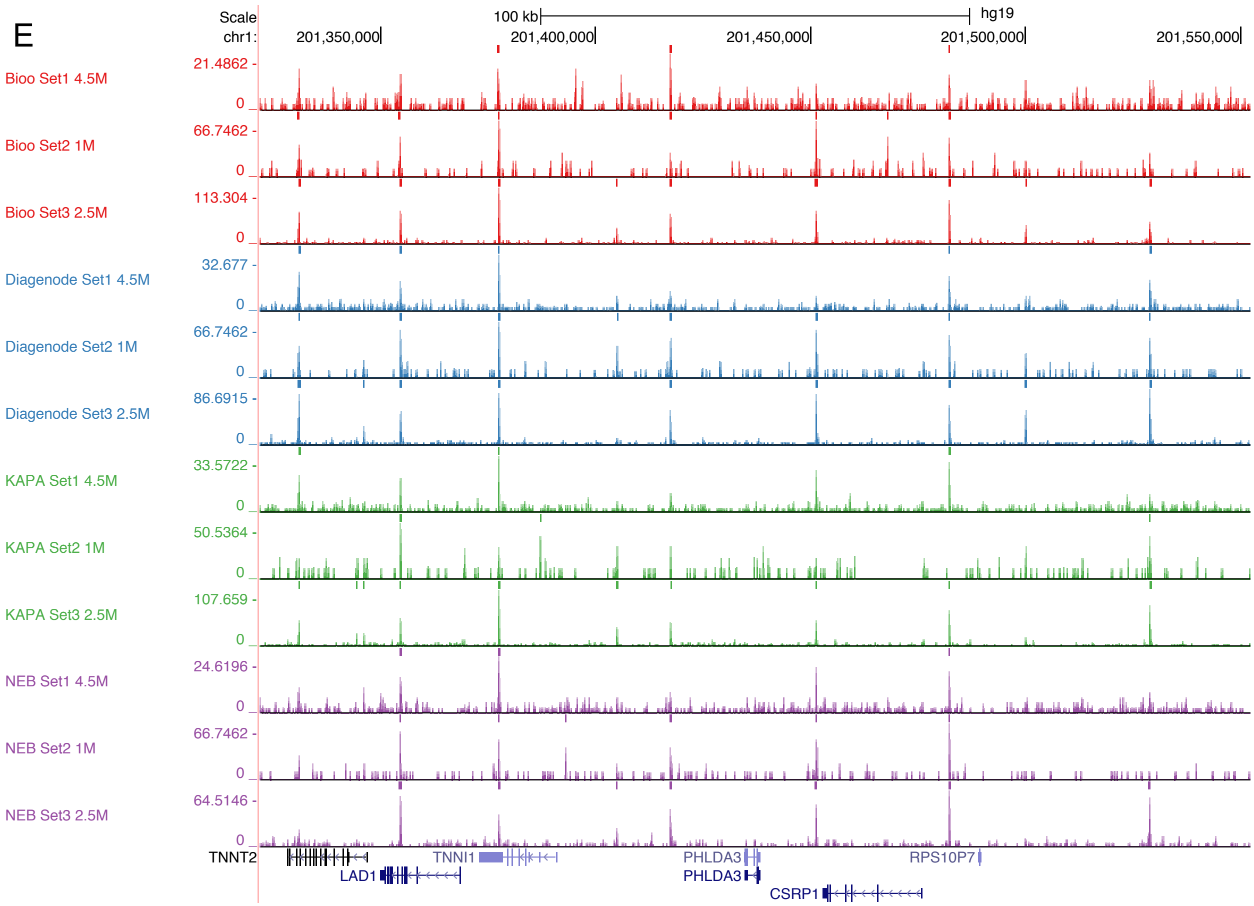
Scale
chr1:
100 kb
hg19
201,350,000
201,400,000
201,450,000
201,500,000
201,550,000
Bioo Set1 4.5M
21.4862 -
0
Bioo Set2 1M
66.7462 -
0
Bioo Set3 2.5M
113.304 -
0
Diagenode Set1 4.5M
32.677 -
0
Diagenode Set2 1M
66.7462 -
0
Diagenode Set3 2.5M
86.6915 -
0
KAPA Set1 4.5M
33.5722 -
0
KAPA Set2 1M
50.5364 -
0
KAPA Set3 2.5M
107.659 -
0
NEB Set1 4.5M
24.6196 -
0
NEB Set2 1M
66.7462 -
0
NEB Set3 2.5M
64.5146 -
0
TNNT2
TNNI1
PHLDA3
RPS10P7
LAD1
PHLDA3
CSRP1

**Figure S1.** Representative screenshots showing the overall signal landscape for each ChIP-Seq library. **A.** H3K4me3 Set 1 libraries. B. H3K4me3 Set 2 libraries. **C.** H3K27me3 Set1 libraries. D. H3K27me3 Set2 libraries. **E.** CTCF libraries. The small bars on top of each track indicate the peaks/broad domains called in each library. The number of reads used to generate the signal landscape of each library is indicated at the end of track name. Please note the purpose of this figure is to demonstrate the success of our libraries instead of a direct comparison among libraries, as different numbers of reads were used to generate these signal tracks.
