## Supplemental Figure 2 for "Commercial ChIP-Seq library preparation kits performed differently for different classes of protein targets"

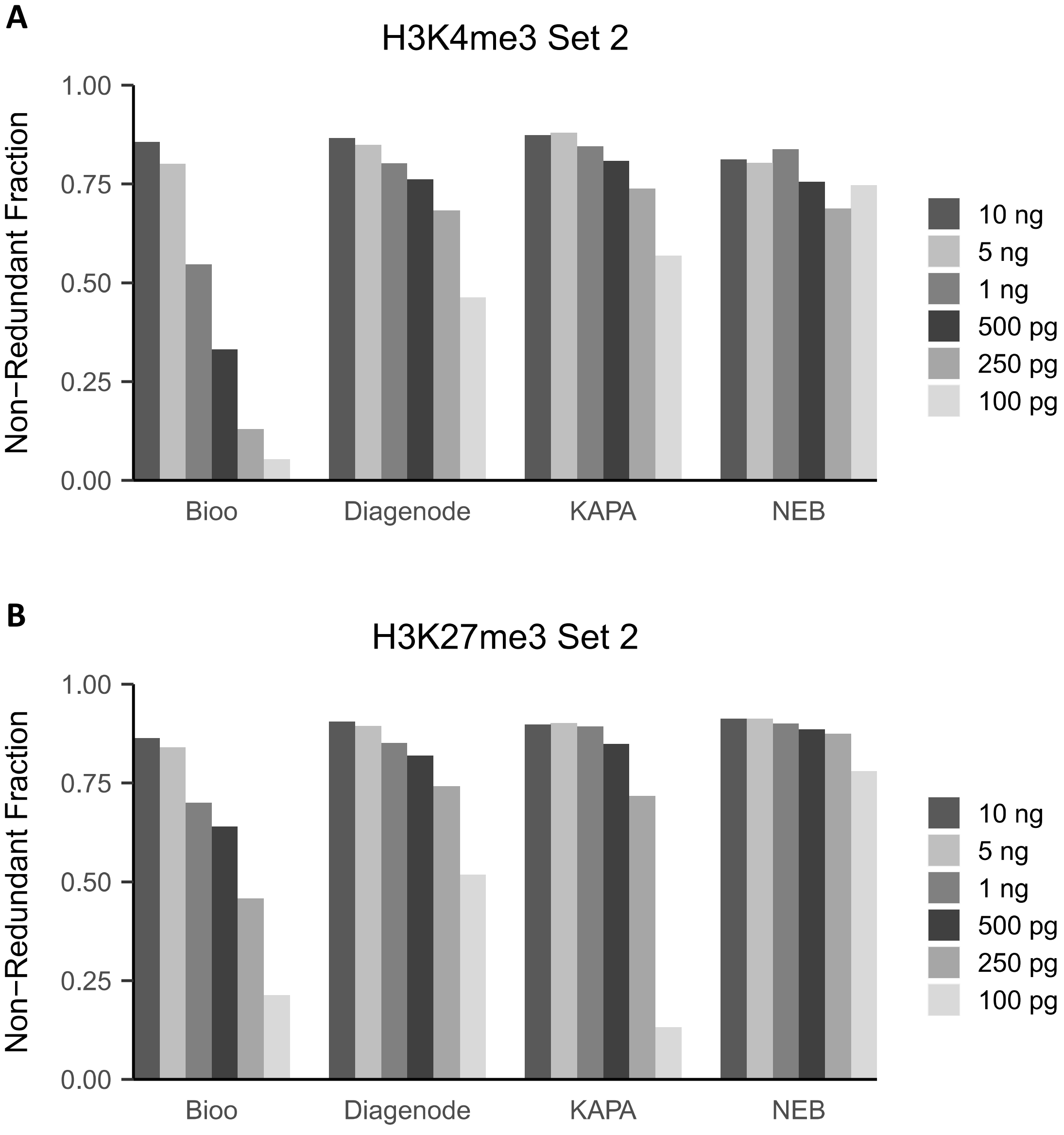
A
H3K4me3 Set 2
Non−Redundant Fraction
1.00
0.75
0.50
0.25
0.00
Bioo
Diagenode
KAPA
NEB
10 ng
5 ng
1 ng
500 pg
250 pg
100 pg
B
H3K27me3 Set 2
Non−Redundant Fraction
1.00
0.75
0.50
0.25
0.00
Bioo
Diagenode
KAPA
NEB
10 ng
5 ng
1 ng
500 pg
250 pg
100 pg

**Figure S2.** Library complexity measured as Non-Redundant Fraction (NRF). **A.** NRF for H3K4me3 set 2 libraries. **B.** NRF for H3K27me3 set 2 libraries.
