## Supplemental Figure 3 for "Commercial ChIP-Seq library preparation kits performed differently for different classes of protein targets"

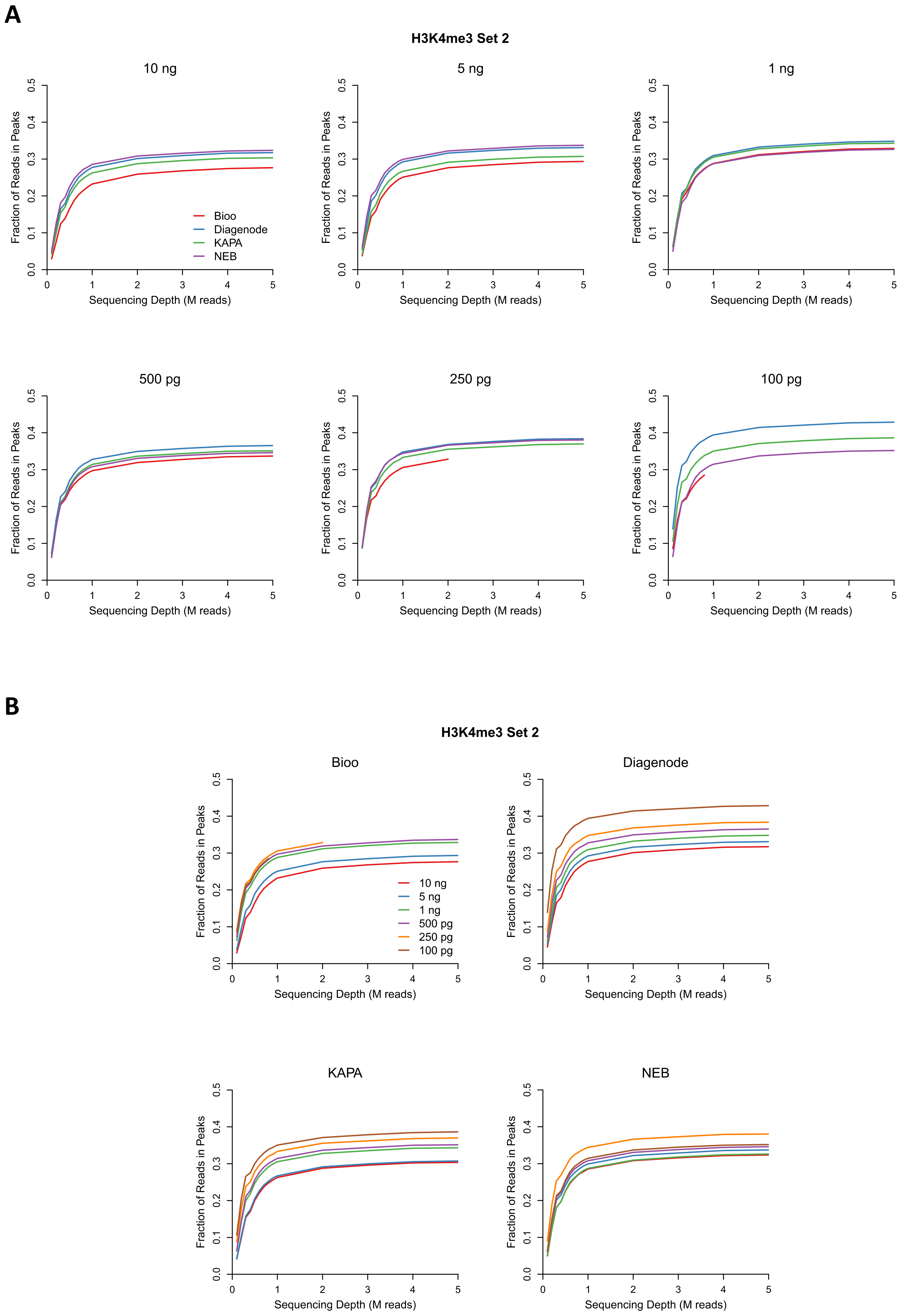
A
H3K4me3 Set 2
10 ng
5 ng
1 ng
500 pg
250 pg
100 pg
Fraction of Reads in Peaks
Sequencing Depth (M reads)
0.0
0.1
0.2
0.3
0.4
0.5
0
1
2
3
4
5
Bioo
Diagenode
KAPA
NEB
B
H3K4me3 Set 2
Bioo
Diagenode
KAPA
NEB
10 ng
5 ng
1 ng
500 pg
250 pg
100 pg

**Figure S3.** FRiP (fraction of reads in peaks) for H3K4me3 plotted against sequencing depth for H3K4me3 set 2 libraries. **A.** Graphs displayed to highlight differences in protocol performance at specific DNA inputs. **B.** Graphs displayed to facilitate comparisons of a single protocol at different DNA inputs.
