## Supplemental Figure 4 for "Commercial ChIP-Seq library preparation kits performed differently for different classes of protein targets"

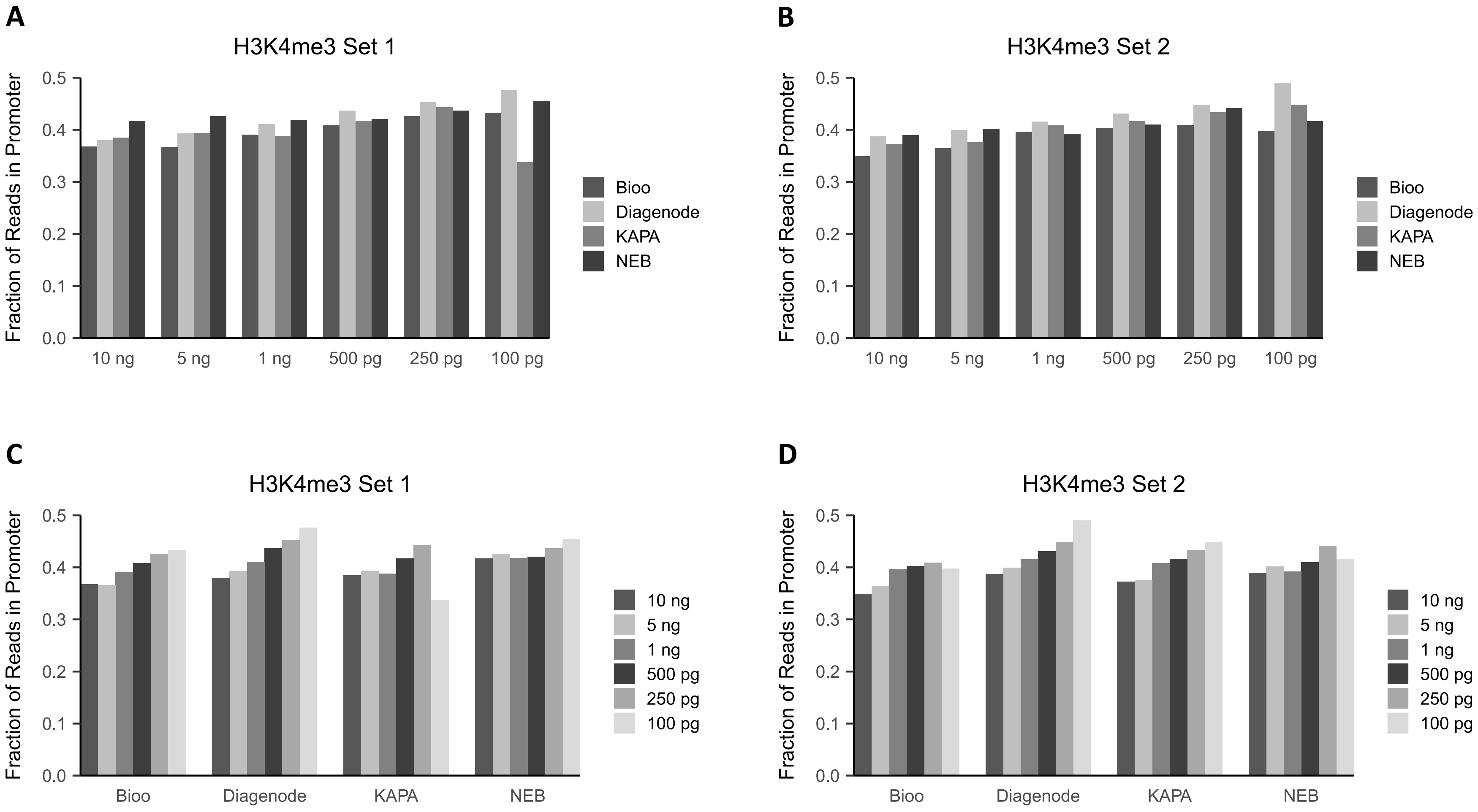
A
H3K4me3 Set 1
Fraction of Reads in Promoter
0.0
0.1
0.2
0.3
0.4
0.5
10 ng
5 ng
1 ng
500 pg
250 pg
100 pg
Bioo
Diagenode
KAPA
NEB
B
H3K4me3 Set 2
Fraction of Reads in Promoter
0.0
0.1
0.2
0.3
0.4
0.5
10 ng
5 ng
1 ng
500 pg
250 pg
100 pg
Bioo
Diagenode
KAPA
NEB
C
H3K4me3 Set 1
Fraction of Reads in Promoter
0.0
0.1
0.2
0.3
0.4
0.5
Bioo
Diagenode
KAPA
NEB
10 ng
5 ng
1 ng
500 pg
250 pg
100 pg
D
H3K4me3 Set 2
Fraction of Reads in Promoter
0.0
0.1
0.2
0.3
0.4
0.5
Bioo
Diagenode
KAPA
NEB
10 ng
5 ng
1 ng
500 pg
250 pg
100 pg

**Figure S4.** Fraction of H3K4me3 reads in promoter regions. **A-B.** Bar graphs displayed to highlight differences in protocol performance at specific DNA inputs. **C-D.** Graphs displayed to facilitate comparisons of a single protocol at different DNA inputs. H3K4me3 set 1 libraries are presented in A and C, and H3K4me3 set 2 libraries are presented in B and D.
