## Supplemental Figure 5 for "Commercial ChIP-Seq library preparation kits performed differently for different classes of protein targets"

H3K4me3 Set 1

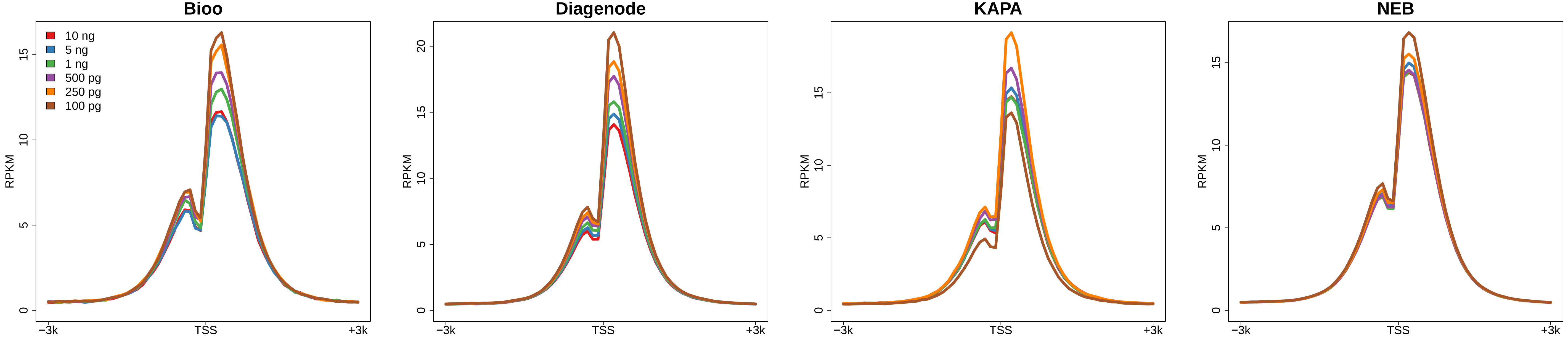


H3K4me3 Set 2

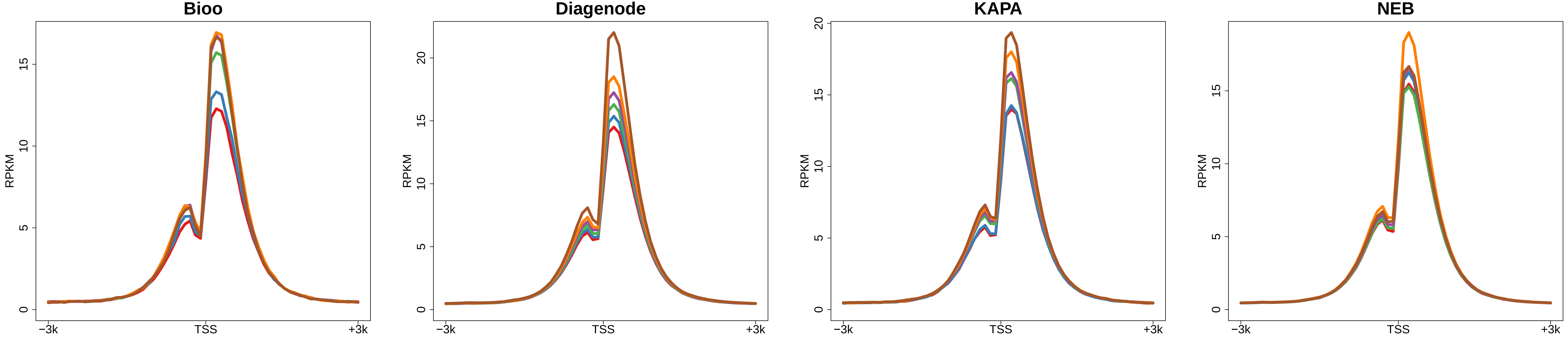

**Figure S5.** Aggregated signal curves around transcription start sites (TSSs) for H3K4me3 libraries organized by protocol.
