## Supplemental Figure 6 for "Commercial ChIP-Seq library preparation kits performed differently for different classes of protein targets"

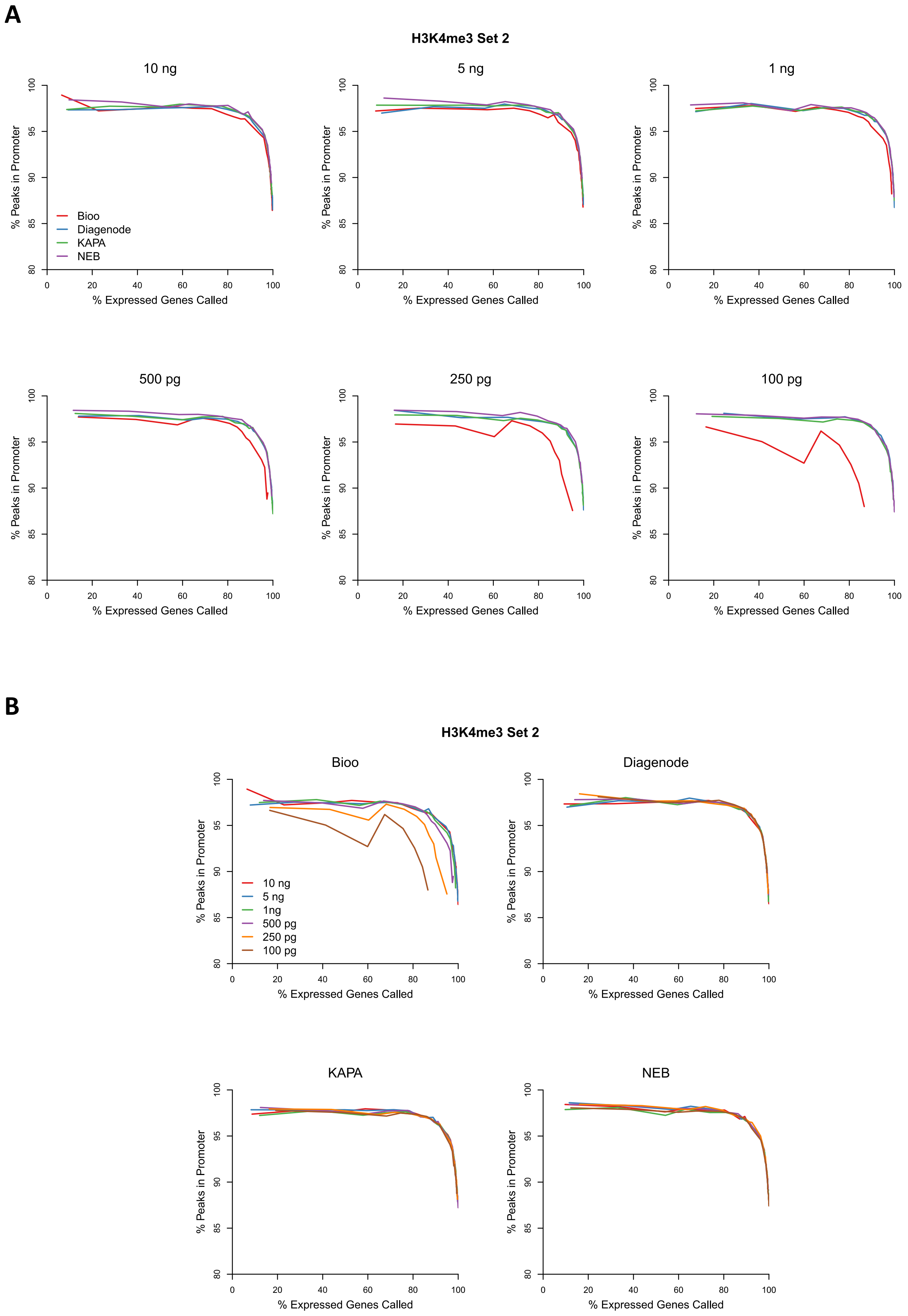
A
H3K4me3 Set 2
10 ng
5 ng
1 ng
500 pg
250 pg
100 pg
% Peaks in Promoter
% Expressed Genes Called
Bioo
Diagenode
KAPA
NEB
80
85
90
95
100
0
20
40
60
80
100
B
H3K4me3 Set 2
Bioo
Diagenode
KAPA
NEB
10 ng
5 ng
1ng
500 pg
250 pg
100 pg

**Figure S6.** The peak calling quality for H3K4me3 measured as the percentage of peaks in promoter regions vs. the percentage of expressed genes marked by peaks at varying sequencing depth for H3K4me3 set 2 libraries. **A.** Graphs displayed to highlight diffences in the performance of each protocol at specific DNA inputs. **B.** Graphs displayed to facilitate comparisons of a single protocol at different DNA inputs.
