## Supplemental Figure 8 for "Commercial ChIP-Seq library preparation kits performed differently for different classes of protein targets"

**A**

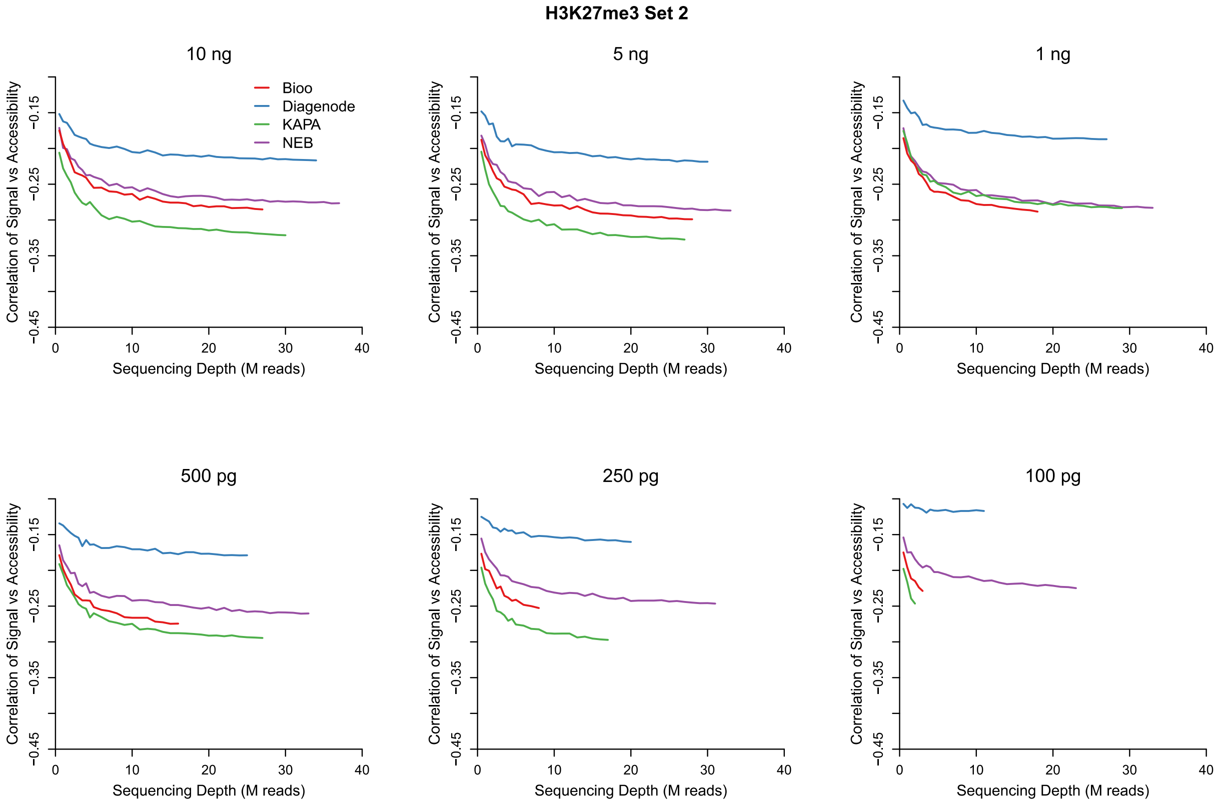

H3K27me3 Set 2
10 ng
5 ng
1 ng
500 pg
250 pg
100 pg
Bioo
Diagenode
KAPA
NEB
Correlation of Signal vs Accessibility
−0.45
−0.35
−0.25
−0.15
Sequencing Depth (M reads)
0
10
20
30
40


**B**

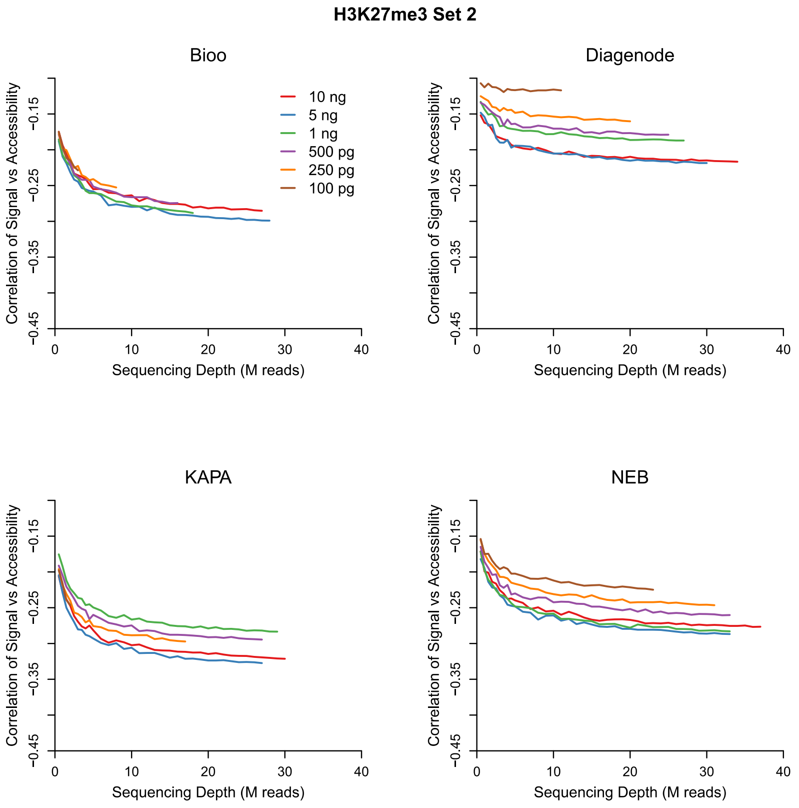

H3K27me3 Set 2
Bioo
Diagenode
KAPA
NEB
10 ng
5 ng
1 ng
500 pg
250 pg
100 pg
Correlation of Signal vs Accessibility
−0.45
−0.35
−0.25
−0.15
Sequencing Depth (M reads)
0
10
20
30
40

**Figure S8.** Spearman correlation of H3K27me3 signal intensity versus chromatin accessibility plotted against increasing sequencing depth for H3K27me3 set 2 libraries. The lower the curve, the better the negative correlation with gene expression. **A.** Graphs displayed to highlight diffences in the performance of each protocol at specific DNA inputs. **B.** Graphs displayed to facilitate comparisons of a single protocol at different DNA inputs.
