## Supplemental Figure 9 for "Commercial ChIP-Seq library preparation kits performed differently for different classes of protein targets"

**A**

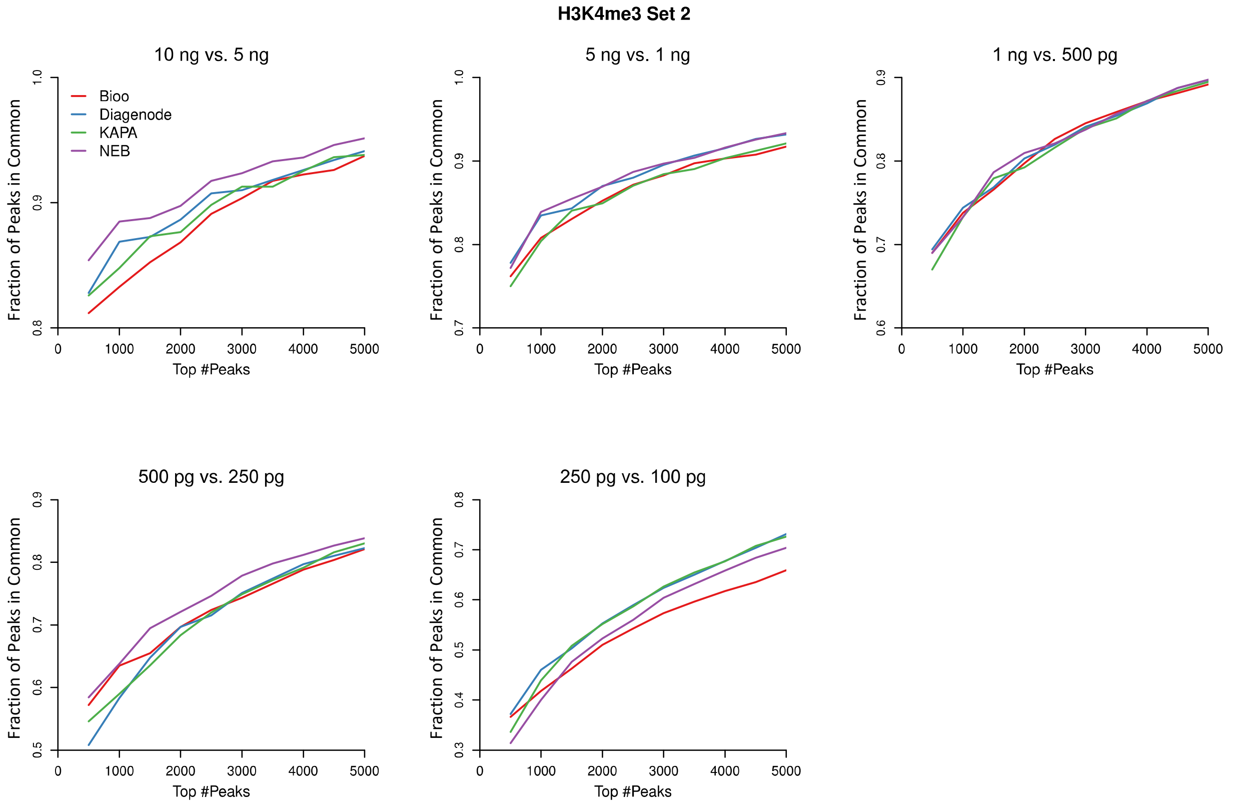
H3K4me3 Set 2
10 ng vs. 5 ng
5 ng vs. 1 ng
1 ng vs. 500 pg
500 pg vs. 250 pg
250 pg vs. 100 pg
Bioo
Diagenode
KAPA
NEB
Fraction of Peaks in Common
Top #Peaks

**B**

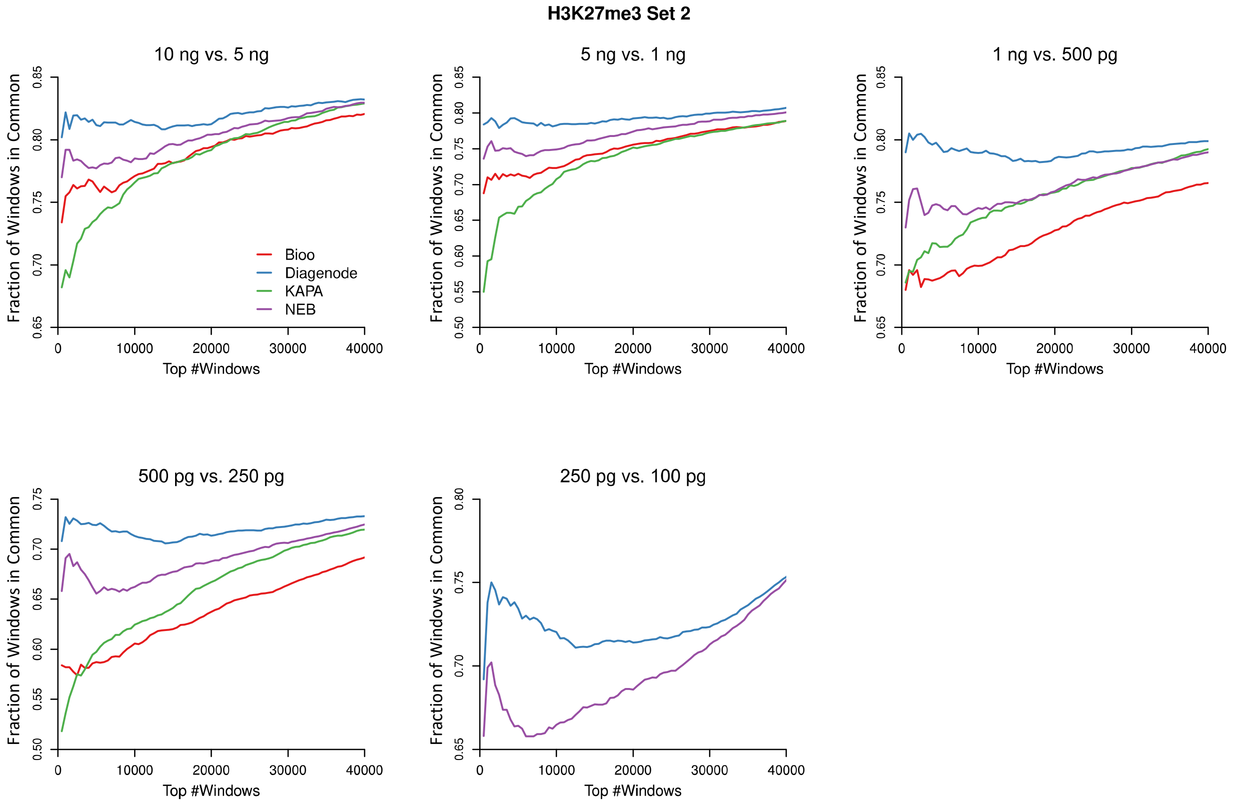
H3K27me3 Set 2
10 ng vs. 5 ng
5 ng vs. 1 ng
1 ng vs. 500 pg
500 pg vs. 250 pg
250 pg vs. 100 pg
Bioo
Diagenode
KAPA
NEB
Fraction of Windows in Common
Top #Windows

**Figure S9.** Reproducibility curves displayed to highlight differences in the performance of each protocol. The X-axis represents the number of top peaks (H3K4me3) or enriched 20 kb windows (H3K27me3) in pairwise comparisons of libraries produced from incremental changes in DNA input (i.e. 10 ng vs. 5 ng, 5 ng vs. 1 ng, 1 ng vs. 500 pg, 500 pg vs. 250 pg, and 250 pg vs. 100 pg). The Y-axis represents the fraction of top peaks or enriched windows that are in common between the two libraries that were compared. H3K4me3 (**A**) and H3K27me3 (**B**) set 2 data are plotted.
