## Supplemental Figure 10 for "Commercial ChIP-Seq library preparation kits performed differently for different classes of protein targets"

**A**

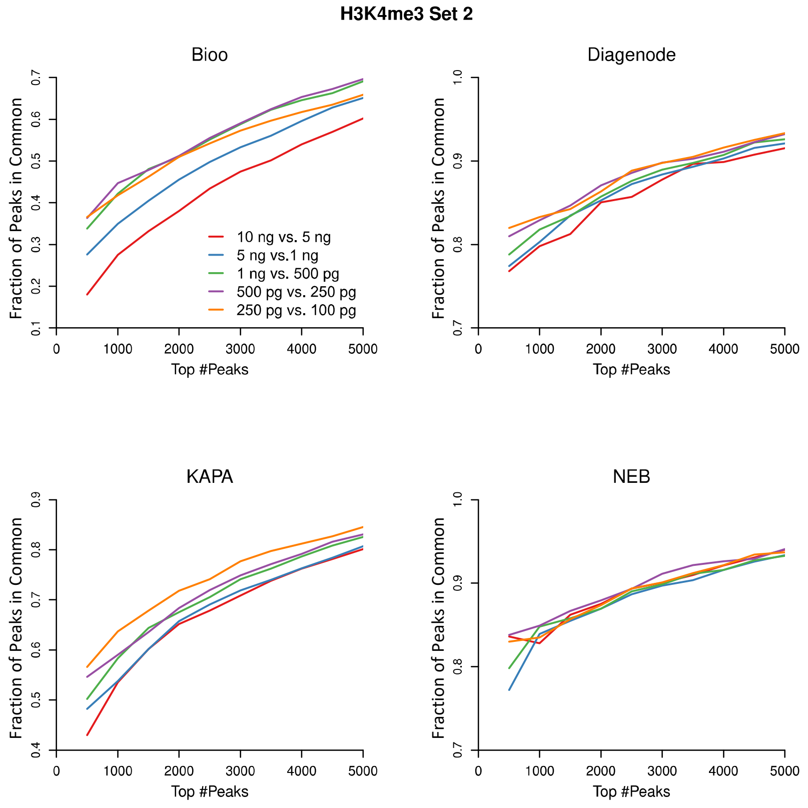
H3K4me3 Set 2
Bioo
Diagenode
KAPA
NEB
Fraction of Peaks in Common
Top #Peaks
10 ng vs. 5 ng
5 ng vs.1 ng
1 ng vs. 500 pg
500 pg vs. 250 pg
250 pg vs. 100 pg

**B**

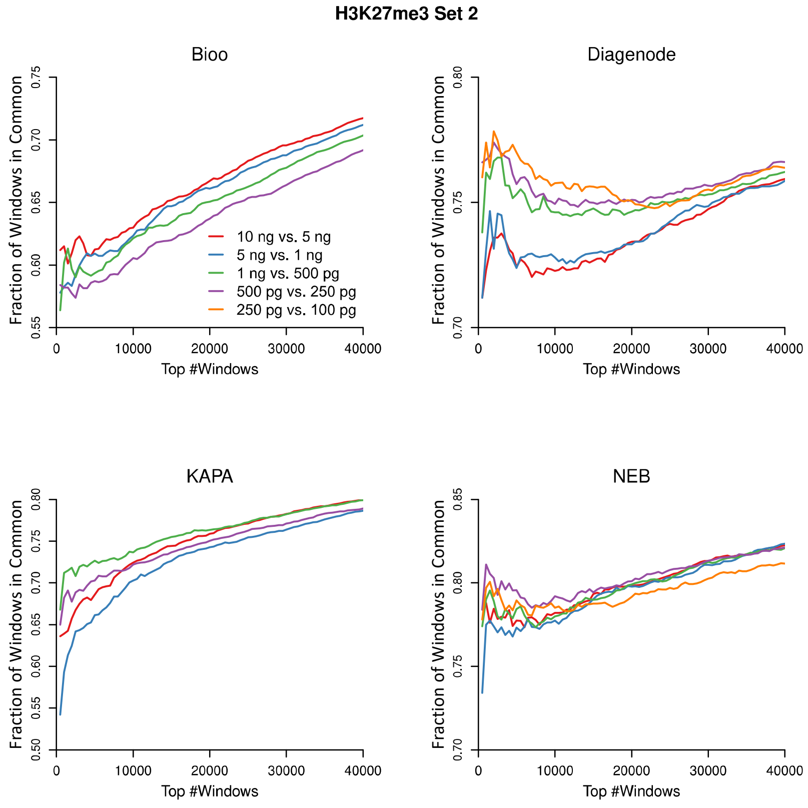
H3K27me3 Set 2
Bioo
Diagenode
KAPA
NEB
Fraction of Windows in Common
Top #Windows
10 ng vs. 5 ng
5 ng vs. 1 ng
1 ng vs. 500 pg
500 pg vs. 250 pg
250 pg vs. 100 pg

**Figure S10.** Reproducibility curves displayed to facilitate single protocol comparisons at incremental changes in DNA input. The X-axis represents the number of top peaks (H3K4me3) or enriched 20 kb windows (H3K27me3) in incremental, pair-wise comparisions of DNA input levels. The Y-axis represents the fraction of top peaks or enriched windows that are in common between the two libraries that were compared. H3K4me3 (**A**) and H3K27me3 (**B**) set 2 data are plotted
