## Supplemental Figure 11 for "Commercial ChIP-Seq library preparation kits performed differently for different classes of protein targets"

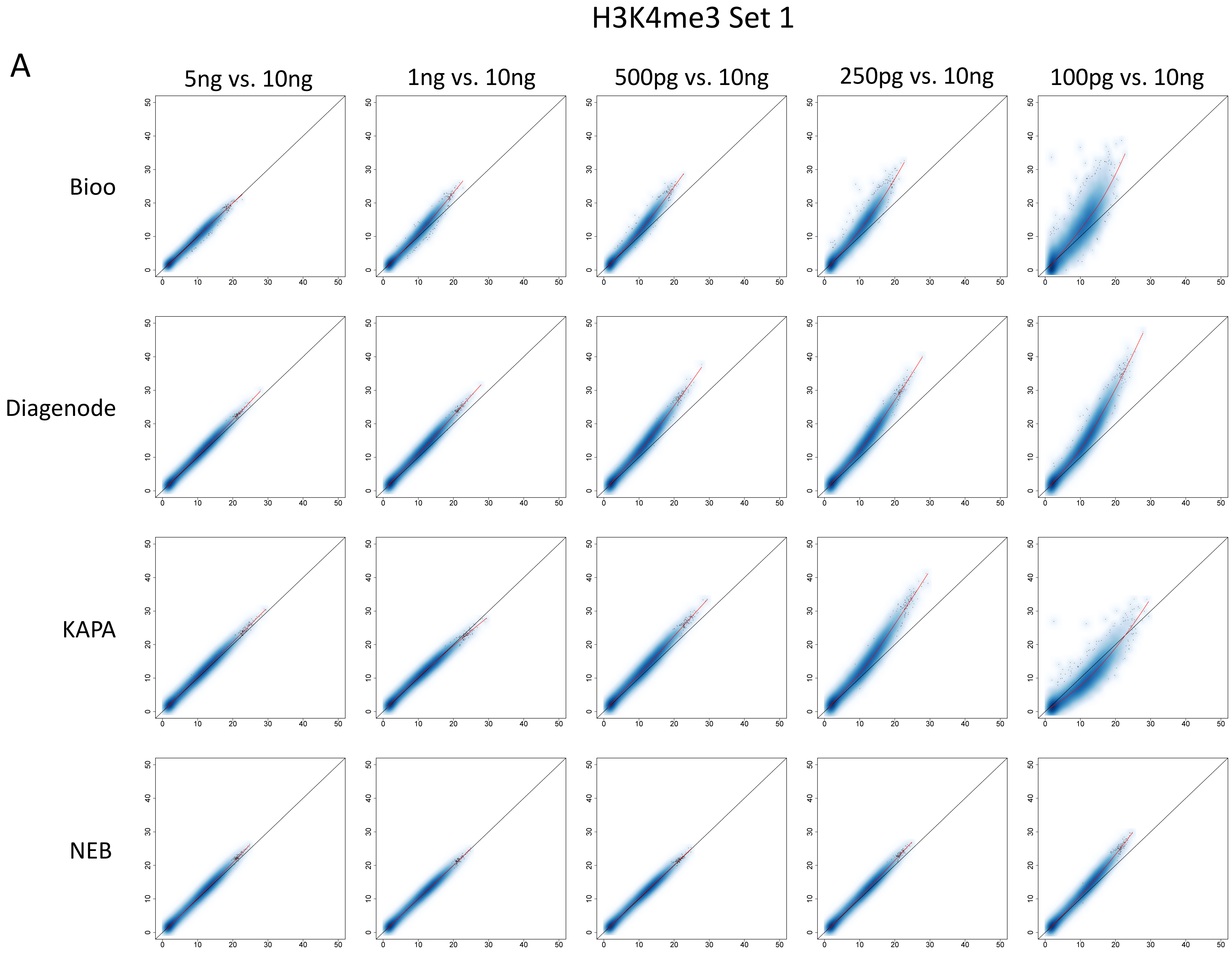
H3K4me3 Set 1
A
5ng vs. 10ng
1ng vs. 10ng
500pg vs. 10ng
250pg vs. 10ng
100pg vs. 10ng
Bioo
Diagenode
KAPA
NEB

### H3K4me3 Set 2

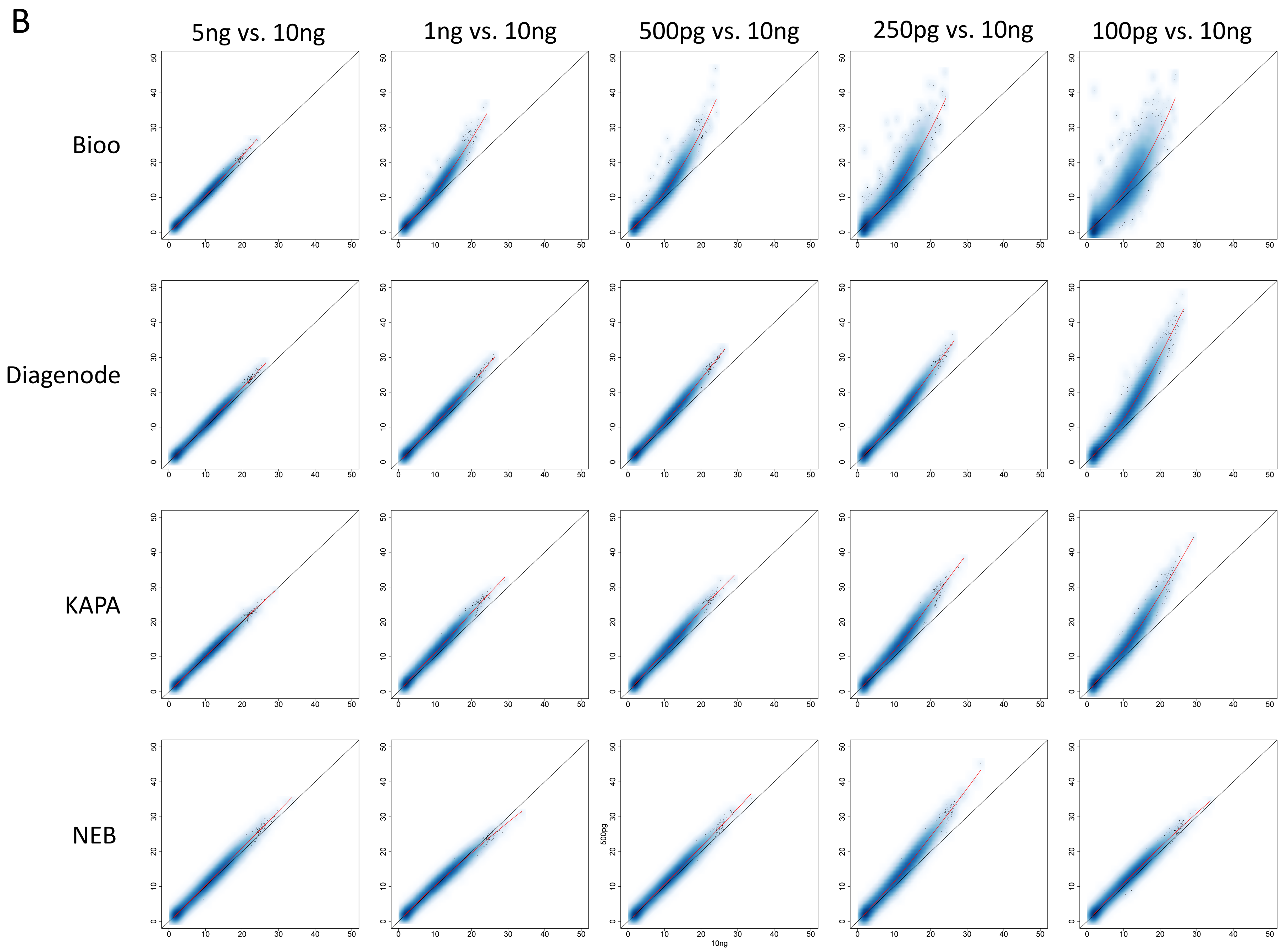

**Figure S11.** Scatter plots showing H3K4me3 signal intensity in peaks for each amount of input DNA (y-axis) compared to 10 ng of input DNA (x-axis) for each protocol. H3K4me3 set 1 (**A**) and set 2 (**B**) library data are presented as scatter plots with smoothed color density, and with red lines representing LOESS-fitted curves.
