## Supplemental Figure 12 for "Commercial ChIP-Seq library preparation kits performed differently for different classes of protein targets"

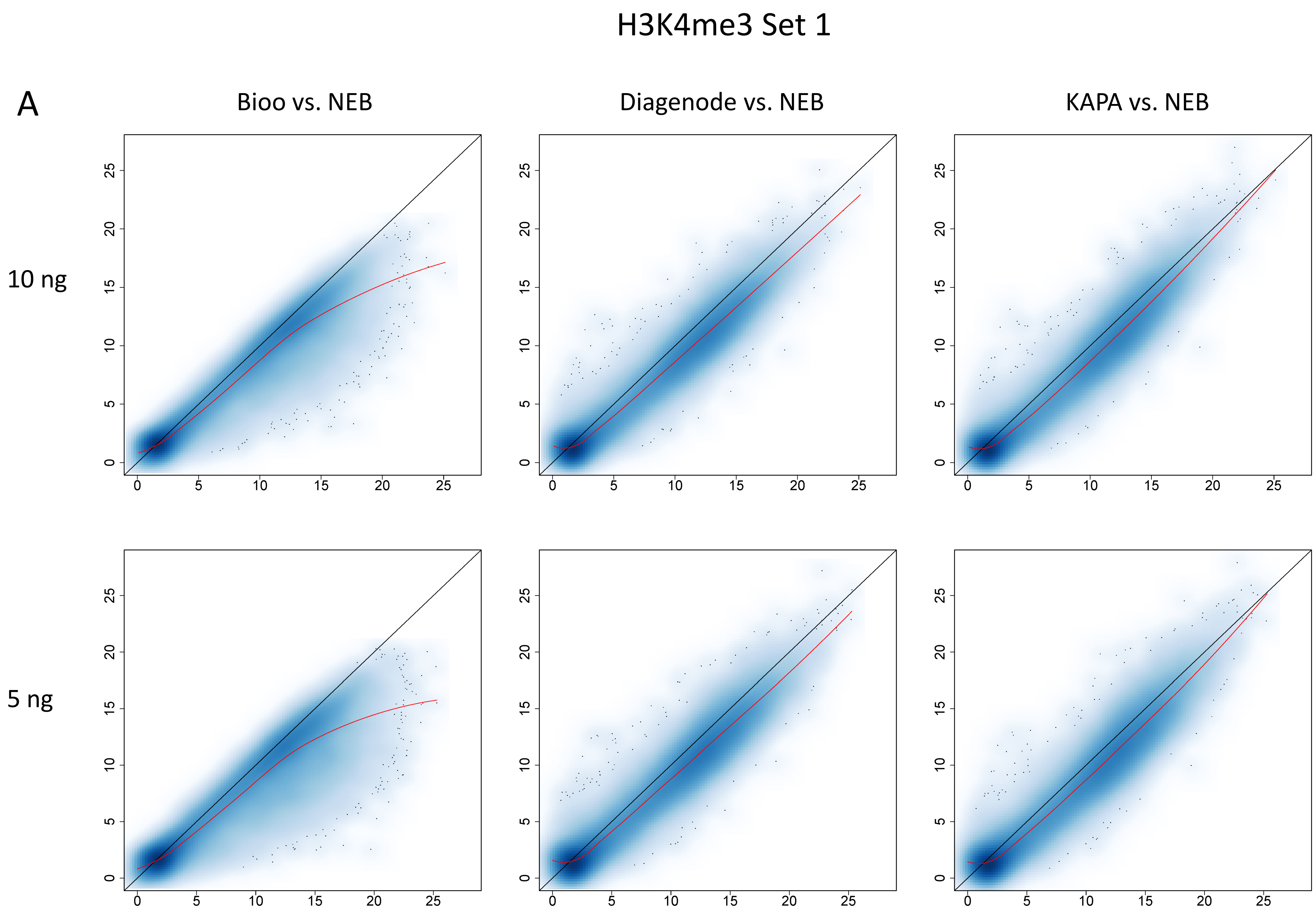

H3K4me3 Set 1
A
Bioo vs. NEB
Diagenode vs. NEB
KAPA vs. NEB
10 ng
5 ng

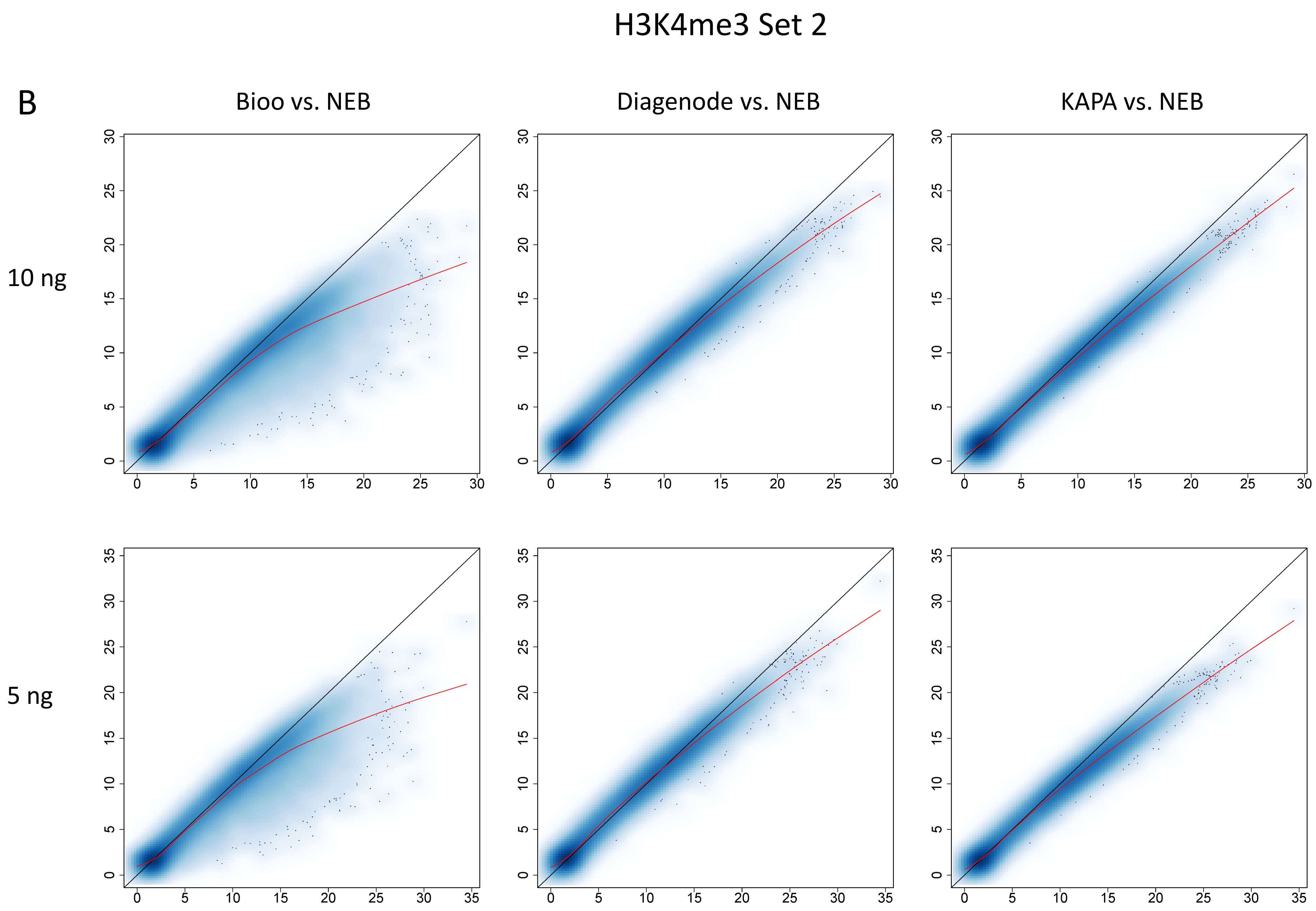
H3K4me3 Set 2
B
Bioo vs. NEB
Diagenode vs. NEB
KAPA vs. NEB
10 ng
5 ng
0
5
10
15
20
25
30
35

**Figure S12.** Scatter plots showing H3K4me3 signal intensity in peaks for each protocol (y-axis) compared to the NEB protocol (x-axis) at DNA inputs of 10 and 5 ng. H3K4me3 set 1 (**A**) and set 2 (**B**) library data are presented as scatter plots with smoothed color density, and with red lines representing LOESS-fitted curves.
