## Supplemental Figure 14 for "Commercial ChIP-Seq library preparation kits performed differently for different classes of protein targets"

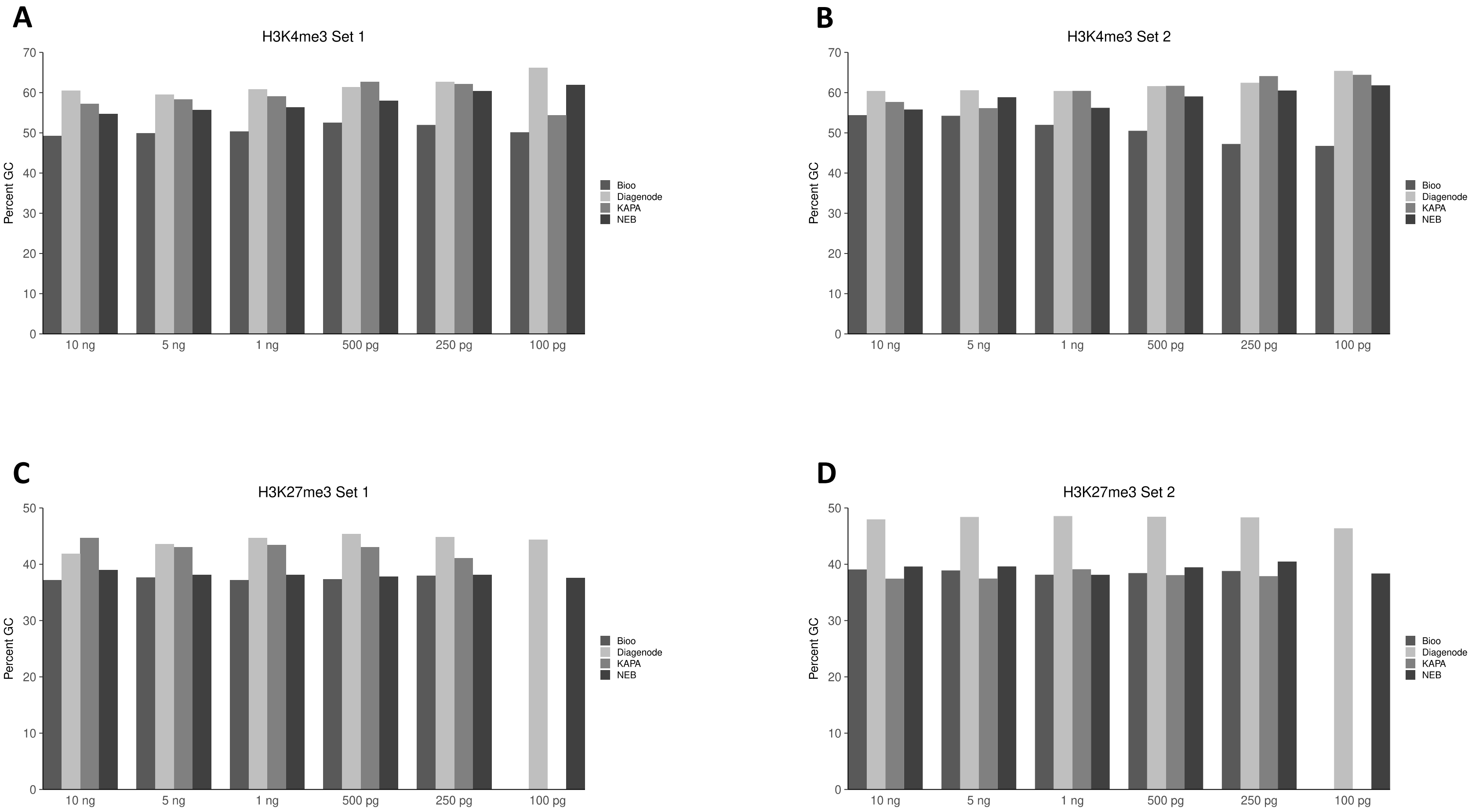
A
H3K4me3 Set 1
B
H3K4me3 Set 2
C
H3K27me3 Set 1
D
H3K27me3 Set 2
Percent GC
10 ng
5 ng
1 ng
500 pg
250 pg
100 pg
Bioo
Diagenode
KAPA
NEB

**Figure S14.** Bar graphs indicating the average GC content of the DNA sequences under peaks uniquely called to each protocol at specific DNA inputs. H3K4me3 set 1 and set 2 libraries are presented in **A** and **B**, respectively. H3K27me3 set 1 and set 2 libraries are presented in **C** and **D**, respectively. H3K27me3 Bioo 100 ng and KAPA 100 ng libraries didn't have enough reads for a meaningful peak calling, thus were skipped.
